# Indigenous peoples in eastern Brazil: insights from 19^th^ century genomes and metagenomes

**DOI:** 10.1101/2022.01.27.477466

**Authors:** Diana Ivette Cruz Dávalos, Yami Ommar Arizmendi Cárdenas, Miriam Jetzabel Bravo-Lopez, Samuel Neuenschwander, Silvia Reis, Murilo Q. R. Bastos, Jesper Stenderup, Fulya Eylem Yediay, Viridiana Villa-Islas, Carlos S. Reyna-Blanco, Claudia Rodrigues-Carvalho, Tábita Hünemeier, Morten E. Allentoft, Carlos Eduardo G. Amorim, J. Víctor Moreno-Mayar, María C. Ávila-Arcos, Anna-Sapfo Malaspinas

## Abstract

Although Brazil was inhabited by more than 3,000 Indigenous populations prior to European colonization, today’s Indigenous peoples represent less than 1% of Brazil’s census population. Some of the decimated communities belonged to the so-called “Botocudos” from central-eastern Brazil. These peoples are thought to represent a case of long-standing genetic continuity bearing a strong craniometric resemblance to that of the oldest Indigenous Americans (“Paleoamericans”). Yet, little is known about their origins and genetic relationship to other Native Americans, as only two “Botocudo” genomes have been sequenced so far and those were surprisingly of Polynesian ancestry. To deepen our knowledge on the genomic history of pre-contact Indigenous Americans and the pathogens they were exposed to, we carbon-dated and sequenced 24 ancient Brazilians (including 22 “Botocudos”) whose remains were hosted at the National Museum of Rio de Janeiro and recovered prior to the tragic 2018 fire. The resulting genomes’ depth of coverage ranged from 0.001× to 24×. Their genetic ancestry was found to be Indigenous American without gene flow from external populations such as Europeans, Africans or Polynesians. Unlike Mesoamericans, the “Botocudos” and Amazonians do not seem to have experienced a population expansion once in the Americas. Moreover, remarkably, their genomes exhibit amongst the lowest levels of heterozygosity worldwide and long runs of homozygosity, which could be explained by unique social practices or a very small effective size. Finally, whole genomes of likely ancient pathogens were recovered, including lineages of Human parvovirus B19 that were possibly introduced after the European contact.

**Significance statement:** To better understand the genetic relationship among Indigenous populations in Brazil, we sequenced the genomes of 24 ancient individuals (22 of which labelled as “Botocudos”, a term used to describe hunter-gatherer tribes) whose remains were hosted at the *Museu Nacional* of Rio de Janeiro prior to the tragic fire that consumed it in 2018. Unlike two previously published “Botocudo” genomes, the 22 “Botocudos” from this study have Indigenous American-related ancestry without any Polynesian-related ancestry, and they are similarly related to several Native Brazilian populations. Finally, unlike Eurasian hunter-gatherers, the “Botocudos” exhibit among the lowest heterozygosity and longest runs of homozygosity worldwide – compatible with a very small effective size and suggesting a unique social structure among hunter-gatherers in the Americas.

## Introduction

Despite decades of studies, the inferred human population history of South America and Brazil remains contentious. The number of human migration waves into the Americas, the population structure of the ancestral population(s) and the relationship between ancient and present-day people are yet to be fully characterized (Willerslev et al. 2021; Skoglund et al. 2016; Pickrell et al. 2014; Bolnick et al. 2016). For instance, it has been proposed that Indigenous Americans descend from a single founding population, and that ancient and present-day populations diverged by 23,000 years at most (Raghavan et al., 2015). Southern Indigenous Americans, including communities such as Yjxa (“Karitiana”, Storto et al. 2021), and northern Indigenous Americans, including Athabascans-speaking populations, were inferred to have split about 15,000 years ago (ya) (Moreno-Mayar, Potter, et al. 2018). Following this event, the populations constituting the southern Indigenous American branch seem to have diversified and moved rapidly across both North and South America within approximately two millennia (Moreno-Mayar, Vinner, et al. 2018; Posth et al. 2018).

The earliest humans in the archaeological record of Brazil, the Lagoa Santa people in Minas Gerais, were descendants of the southern Indigenous American expansion and their skeletal remains have been dated to about 10,000 years old (yo) (Neves 2017). When compared to Native Americans (such as Mixe and Wixárika or “Huichol”), the Lagoa Santa people exhibit a higher genetic affinity to Australasian populations (Moreno-Mayar, Vinner, et al. 2018). It is highly debated whether this Australasian signal is evidence for genetic structure in the founding Indigenous American populations or if it reflects multiple independent migration events into the Americas (Moreno-Mayar, Vinner, et al. 2018; Raghavan et al., 2015; Skoglund et al. 2015). Furthermore, although this “Australasian signal” was initially found in present-day Amazonians (such as Yjxa and Paiter or “Surui”; Raghavan et al., 2015; Skoglund et al. 2015), a recent study (Castro e Silva et al. 2021) has suggested that it is widespread in South America.

At the time of the European contact around 1500 CE, it is estimated that about 3,000 indigenous communities inhabited Brazil’s territory (Pagliaro et al. 2005), covering an extensive linguistic diversity. For instance, Moore (2006) has classified 150 present-day languages into six major stocks: Tupi, Macro-Jê, Karib, Arawak, Pano and Tukano. Tupi and Macro-Jê speakers stand for a large part of the people speaking an Indigenous language in Brazil, with the Tupi-speakers embodying 32% of the population, and Macro-Jê speakers representing 23% of the surveyed population in 2010 (IBGE 2010). Tupi-speaking communities have been associated with agricultural practices and a sedentary lifestyle, while Jê-speaking peoples were organized in nomad or semi-nomad bands subsisting on a hunting-fishing-gathering strategy. Tupi-speaking communities, especially those of the Tupi-Guarani branch, are better represented in archaeology, linguistics and genomics, with studies suggesting that their ancestors expanded from northwestern Amazon between 2,000 and 3,000 years before present (BP) (Castro e Silva et al. 2020; Galucio et al. 2015; Ramallo et al. 2013; Silverman et al. 2008; Walker et al. 2012), with the Tupi-Guarani branch reaching southwest, northeast and coastal Brazil, becoming the predominant communities on the Atlantic coast (Paraíso 1992). During this expansion, gene flow has been identified between Tupi-speaking and Jê-speaking populations from central Brazil (Castro e Silva et al. 2020). Less is known about the genomic history of Jê-speaking communities, but they have been reported to be more closely related to the 5,800-year-old Moraes (associated to “*sambaquis*” or shellmound constructions in southeast Brazil) and the 2,000-year-old Laranjal people (associated to sambaquis in southern Brazil) than the Tupi-speaking peoples are (Castro e Silva et al. 2020; Posth et al. 2018).

Of the 3,000 pre-contact nations, it is estimated that only about 250 Indigenous communities, and approximately 70 uncontacted populations in Brazil still exist in 2021, altogether comprising between 0.2% and 0.4% of the total Brazilian population (Azevedo 2018; IBGE 2010; Pagliaro et al. 2005). The decline in Indigenous peoples has been in part attributed to the introduction of infectious diseases and armed conflicts with the European conquerors (Paraíso 1992; Steward 1949). During the European colonization of Brazil, many communities, mostly in Eastern Brazil, were referred to as “Botocudos” by the European settlers (Ehrenreich 2014; Paraíso 1992), and as “Aymorés” (meaning wandering enemies) by the Tupi-speakers of the Brazilian coast (Ehrenreich 2014; Paraíso 1992). It is thought that the so-called “Botocudos” (or “Aymorés”) were Jê-speaking hunter-fisher-gatherers (Ehrenreich 2014; Paraíso 1992). Although the term “Botocudo” was coined from the use of wooden disks (“*botoques”* in Portuguese) in lips and ears, it was used in a broad sense around the 18^th^ century to refer to peoples that had not yet “assimilated” the European regime (Paraíso 1992). Thus, many descriptions of central-eastern Native Brazilians from the historical records are inaccurate or unspecific due to the generalized use of the term “Botocudo”. For instance, some authors relate, among others, the Pojixá, Naknanuk, Nakrehé, Krenak and Aranã as “Botocudos” (Ehrenreich 2014; Paraíso 1992). Except for the Krenak and Aranã peoples, the aforementioned “Botocudos” have been considered “extinct” following the introduction of harsh “assimilation” or “integration” policies in the 19th century (Bieber 2014; Missagia De Mattos 2017; Paraíso 1992). However, despite a prevalent narrative of “Botocudos” not having present-day descendants (Gonçalves et al. 2010; Langfur 2002; Paraíso 1992), 3,159 people identified themselves as belonging to the “Botocudo” ethnicity in the 2010 Brazilian demographic census (whereas 359 identified as “Krenak”, and 210 as “Aranã”; (IBGE 2010).

The study of past and present populations’ genomes has helped to throw light on their genetic history (Nielsen et al. 2017; Willerslev et al. 2021), as well as to demonstrate links to present day peoples (Schroeder et al. 2018), and to open up the possibility of repatriation of the remains of the studied individuals to their community of origin (Moreno-Mayar, Vinner, et al. 2018; Phillips 2019; Rasmussen et al. 2015; Wright et al. 2018). In addition to uncover aspects of human history, ancient DNA has been key to pinpoint the identity and spread routes of microbes across time (Bos et al. 2014; Majander et al. 2020; Mühlemann, Jones, et al. 2018; Rascovan et al. 2019). Nevertheless, vast regions in South America are underrepresented in genomic and metagenomic studies of present-day and ancient populations. For instance, in Brazil, genome-wide data has been generated for a few Indigenous nations, and only a handful of them have been shotgun-sequenced. Notably, most of those Indigenous populations are located along the Amazon basin (northern Brazil) and south of the Atlantic Forest (southern Brazil). As a result, the genetic history of Native populations from central-eastern Brazil is largely unknown. Similarly, routine metagenomic screens from shotgun-sequencing experiments, which inform on the commonly circulating pathogens in a determined context, are scarce.

The National Museum of Rio de Janeiro (*Museu Nacional*) hosted the skulls of 35 individuals which were collected mainly in the 19^th^ century and labelled as “Botocudos”. Two “Botocudo” genomes from the collection were previously sequenced, surprisingly revealing that the individuals’ genetic ancestry was Polynesian without any gene flow from Indigenous Americans nor any other populations (Malaspinas et al., 2014). Yet, as only two genomes among the 35 were sequenced, several questions remained open, such as: (i) how common Polynesian ancestry was among the “Botocudos”. (ii) what is the genetic relationship between “Botocudos” and ancient and present-day Native Americans, such as Tupi-speaking and Jê-speaking populations?; (iii) what are the patterns of genetic diversity in “Botocudos” and the resulting population and social structure of the “Botocudos” inferred from them?; (iv) is there any evidence for “Australasian” gene flow in the “Botocudos”, as reported in ancient (Lagoa Santa) and present-day (Yjxa, Paiter, Xavante) Indigenous Brazilians?; and (v) which ancient pathogens can be identified among the Indigenous Brazilians from the anthropological collection?

To answer these questions, we sequenced the genome, radiocarbon dated and generated isotope data for 24 ancient individuals from Brazil whose remains were hosted at the *Museu Nacional*. This set comprises one individual excavated from a sambaqui construction in Santa Catarina state, one mummy from Minas Gerais state, and 22 “Botocudos” from Minas Gerais, Bahia and Espírito Santo states. We conducted population genetic analyses and assessed their relationship to previously studied present-day and ancient populations, and examined the sequence data for evidence of ancient microbes. This study represents the characterization of a part of the anthropological collection from the *Museu Nacional* prior the tragic fired that consumed the museum (including the skeletal remains analyzed here) in September 2018.

## Results and discussion

### Description of the individuals studied

Prior to the fire, the anthropological collection from the *Museu Nacional* had 35 skulls labeled as “Botocudo” from central-eastern Brazil, of which 22 were selected for DNA and isotope analyses (Datasets S1, S2, Table 1 and Fig. 1), and we refer to them as “Botocudos” hereafter. In addition to the “Botocudo” individuals, we sequenced the genomes of two other individuals from the anthropological collection, including a mummified individual (MN1943, Table 1) of unknown cultural affiliation found in a cave in Minas Gerais (Brazil), and an individual recovered from the shell mound Sambaqui de Cabeçuda (MN01701, Table 1) in the state of Santa Catarina (Brazil). We refer to these individuals as mummy (MN1943) and Sambaqui (MN01701) hereafter. Thus, a total of 24 ancient individuals were sampled for this study (Datasets S1 and S2). One tooth was sampled for 20 ”Botocudos”, the Sambaqui and the mummy. For the remaining two “Botocudos” (MN00019 and MN0008), different portions of the skull were sampled, including the petrous bone for MN0008 (Table 1 and Dataset S2).

**Figure 1.**
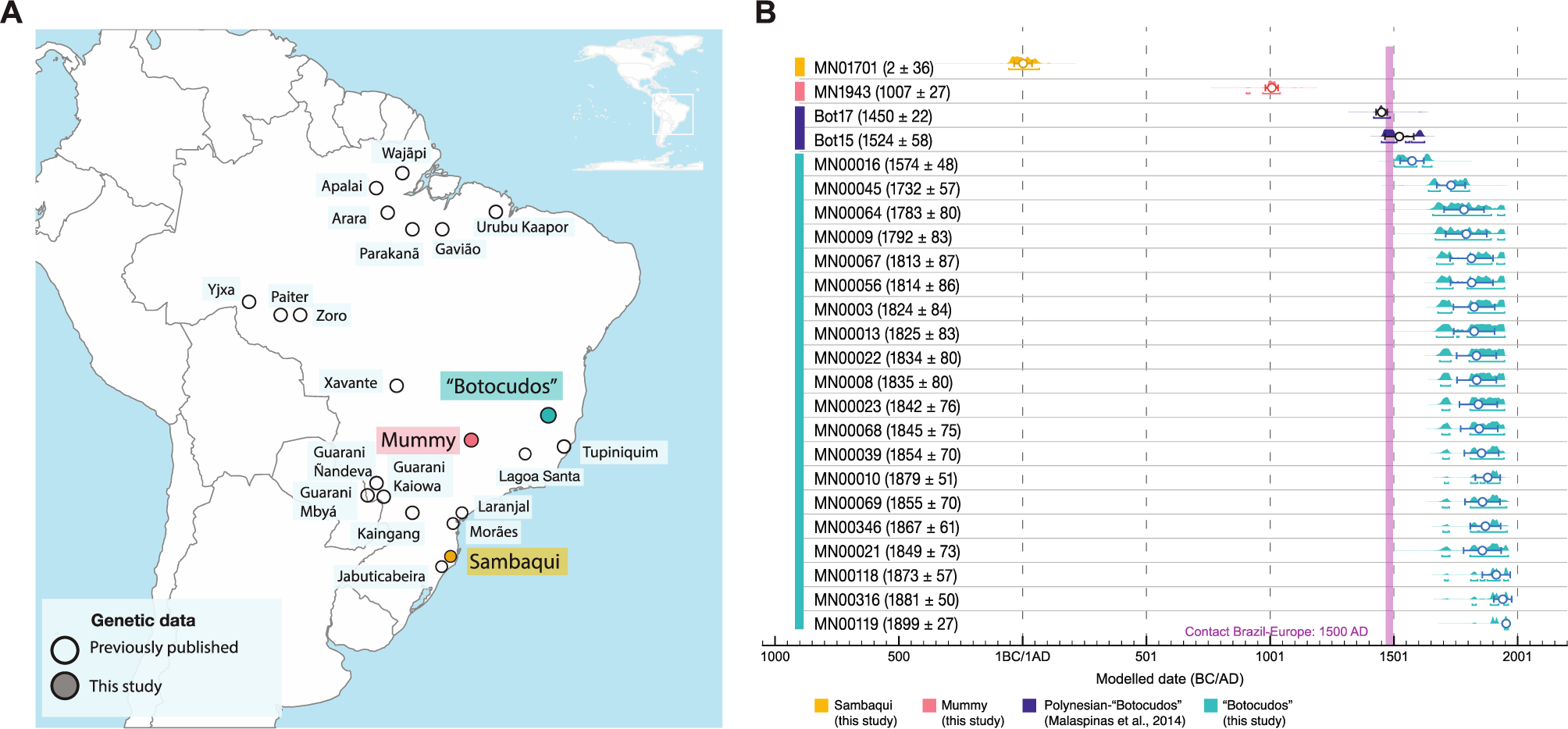
Geographical origin and calibrated radiocarbon dates.| (A) Map showing the provenance of the 24 individual remains sequenced in this study, and the location of previously published ancient and present-day populations from the region. (B) Highest posterior densities of the calibrated radiocarbon-dates for 22 individuals dated in this study, and the two Polynesian-“Botocudo” individuals analyzed in Malaspinas et al. (2014). Color bars on the left highlight labels used throughout the study: yellow for the Sambaqui individual, pink for the mummy, cyan for the “Botocudos” from this study, and dark blue for the two “Botocudos” individuals published in Malaspinas et al., (2014). The colored bars are followed by the individuals’ IDs. Numbers in parentheses indicate mean calibrated date (CE) plus-minus one standard deviation. The mean is also indicated with a white circle underneath the distributions. Brackets below the distributions indicate the highest posterior density interval that encompasses approximately 95.4% of the total area of the distribution. The populations Paiter and Yjxa have been labeled as “Surui” and “Karitiana”, respectively, in previous publications.

**Table 1.**
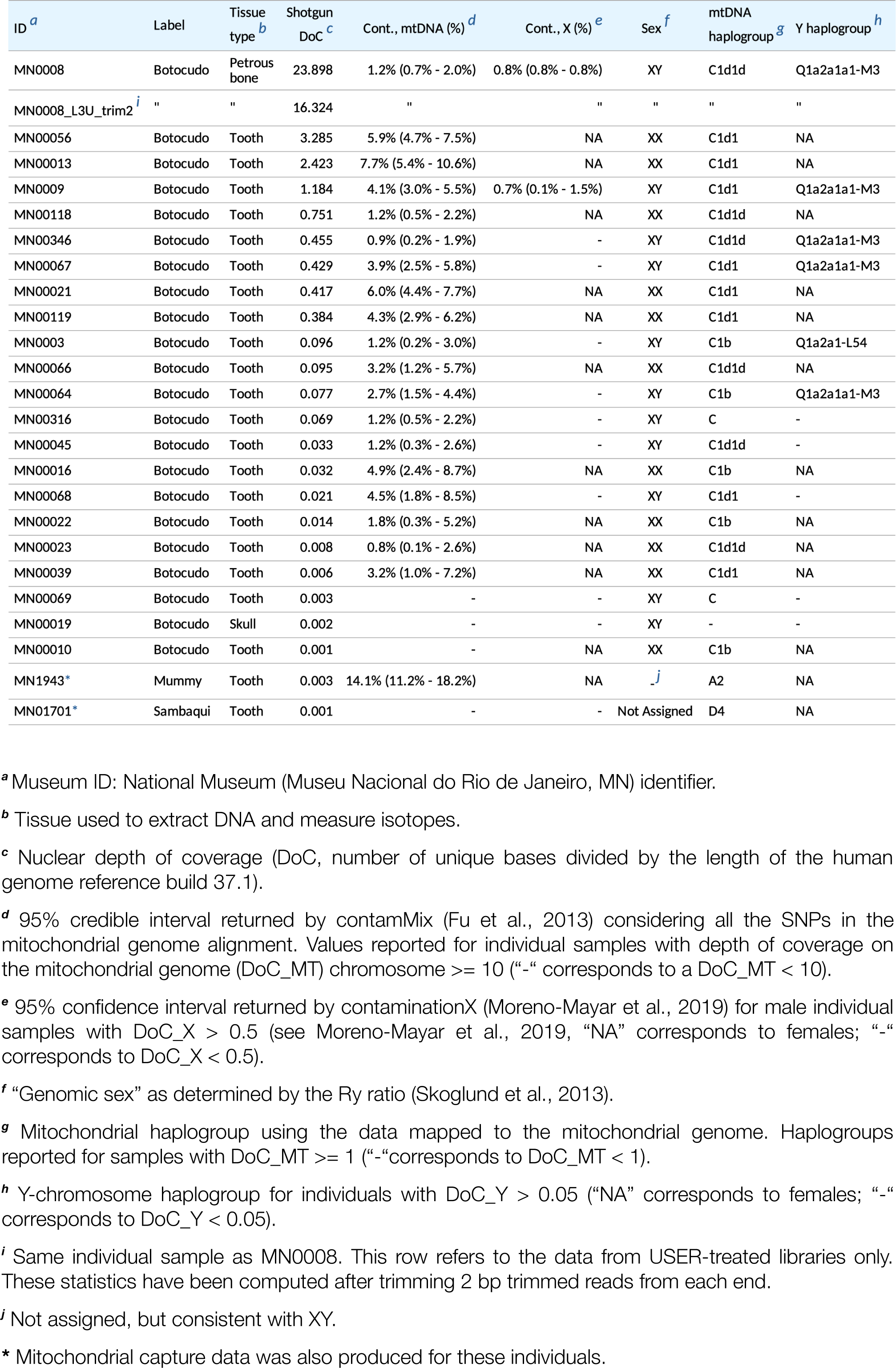
Description of the individuals sequenced in this study.

### Radiocarbon dates and ancient genomes

Twenty-two samples (teeth crown and petrous bone) from 20 “Botocudos”, the Sambaqui and the mummy individuals, were radiocarbon dated at the 14Chrono Laboratory (Queens University, Belfast, UK). The dates for the Sambaqui and the mummy were calibrated to pre-European contact (around 2±36 AD and 1007±27 AD, respectively). The mean calibrated dates for the “Botocudos” range from the 16^th^ to the 19^th^ century (Table 1, Fig. 1B). Hence, all “Botocudos” from this study date to the post-European contact period, and they are more recent than the two previously studied 16^th^-17^th^ century Polynesian-“Botocudos” (Malaspinas et al., 2014).

For the 24 individuals, DNA was extracted, built into Illumina libraries, and sequenced in dedicated ancient DNA clean-room facilities at the GLOBE Institute at the University of Copenhagen. For 20 individuals, the human DNA content ranged from 0.02% to 8% for (Dataset S2), as expected for tropical environments (Allentoft et al. 2012; Hughey et al. 2008; Smith et al. 2003), while the depth of coverage (DoC; the average number of reads covering each position of the genome) ranged between 0.001× and 0.75× (Table 1). The remaining four individuals had better preservation, with human DNA contents between 11% to 47%. The highest value (47%) was found for the DNA extracted from petrous bone (MN0008). For those four well-preserved “Botocudos”, the DoC ranged from 1.2× to 3.3× for MN0009, MN00013 and MN00056 and reached 23.9× for MN0008. For the latter individual, one library (out of three) was prepared from a partially USER-treated DNA extract to reduce post-mortem damage; the data from this USER-treated library achieved a 16.3× DoC (Table 1) after trimming 2 bp at read termini (see Dataset S2 for statistics prior to trimming).

In addition to genome-wide shotgun sequencing data, mitochondrial DNA (mtDNA) was enriched with targeted capture probes for the 1,000-year-old mummy and the 2,000-year-old Sambaqui individuals. As a result, the mtDNA DoC increased from 1.0× to 4.8× for the mummy and from 1.2× to 19.4× for the Sambaqui (Dataset S2).

For all libraries, the sequenced reads showed the expected ancient DNA signatures including molecular damage and fragmentation with deamination rates ranging from 5% to 30% at the read termini, as well as short read lengths averaging from 42 bp to 65 bp per individual (Figs. S1 and S2 and Datasets S2-S4). The average error rates ranged between 0.4% and 2.0% across all ancient individuals (Fig. S3). We note that error rates are driven upwards by an excess of transitions due to post-mortem damage (Figs. S4). The average error rate for MN0008 decreased from 1.14% to 0.08% after USER treatment and trimming 2 bp at the read termini (Fig. S5).

The genetic sex of the individuals was determined by comparing the number of reads mapped to the Y chromosome relative to those mapped to the X and Y chromosomes as in (Skoglund et al. 2013). Eleven “Botocudo” individuals were classified as XY, and eleven “Botocudo” individuals as XX (Table 1). The data for the 1,000-year-old-mummy was consistent with an XY karyotype (male), while the sex for the Sambaqui individual could not be determined (Table 1).

We estimated DNA contamination based on haploid chromosomes (X chromosome in males, and mitochondrial genome for all the individuals), restricting the analyses to individuals with a depth of coverage above 0.5× on the X chromosome (Moreno-Mayar et al. 2020) for males (n = 2 males), and above 10× on the mitochondrial genome (n = 20 individuals). Based on X-chromosome data, we estimated less than 1.5% of contamination for the two males analyzed (MN0008 and MN0009, Table 1). Furthermore, we obtained point contamination estimates based on mitochondrial data below 5% for 16 out of 20 individuals. The remaining 4 individuals with mitochondrial DoC >= 10× (MN00056, MN00013, MN00021, and MN1943, the latter including enriched mtDNA data) had higher contamination estimates (with point estimates at 5.9%, 7.7%, 6.0% and 14.1%, respectively, Table 1). Since the four individuals with the highest contamination estimates (>5%) did not stand out in any of the performed analyses based on nuclear DNA, they were analyzed together with the other individual samples below. However, caution should be applied when including the data from these individuals in future studies.

Depending on the analyses that were performed, we used either of the following five sets: (i) the 24 newly sequenced ancient individuals (22 “Botocudos”, the 1,000-year-old-mummy and the 2,000-year-old Sambaqui); (ii) the 22 “Botocudo” individuals; (iii) nine “Botocudos” with a DoC above 0.1×; (iv) the two highest depth “Botocudos” (MN00056 and MN0008); (v) the high-depth USER treated 16× “Botocudo” genome (MN0008). For the latter, to further reduce the effect of ancient DNA damage, for some analyses, only the data from the USER-treated library were used after trimming the read termini for 2 bp. For all sets we considered pseudo-haploid calls or genotype likelihoods for every individual, except for set v, where we called diploid genotypes (see “Mapping and variant calling” in Materials and Methods).

### No further evidence of Polynesian-South American contact among the “Botocudos”

Two previously published “Botocudo” genomes were found to be of Polynesian ancestry (Goncalves et al. 2013; Malaspinas et al., 2014) without evidence of gene flow from Indigenous Americans (and hereafter referred to as Polynesian-”Botocudos”). To assess whether there were additional individuals with Polynesian ancestry among the 24 sequenced individuals, we compared the newly sequenced ancient genomes to a set of genomes from worldwide populations (including Polynesian and Indigenous American populations, Malaspinas et al., 2014; Wollstein et al. 2010; Xing et al. 2010) with a dimension-reduction technique (classical multidimensional scaling, MDS; see, e.g., Cox et al. 2001) as well as a genetic clustering approach (NGSadmix and fastNGSadmix, Jørsboe et al. 2017; Skotte et al. 2013).

All newly sequenced ancient genomes group together with the Native American populations in the MDS analyses (Fig. 2A and Datasets S5 and S6). In contrast, the two previously-sequenced Polynesian-“Botocudos” (Fig. 2A, blue diamonds) group with Polynesian populations, as reported before (Malaspinas et al., 2014). This shows that – unlike the previously published Polynesian-“Botocudos” genomes — the twenty-four individuals in this study have a closer genetic affinity to people from the Americas than to any other population in this panel.

**Figure 2.**
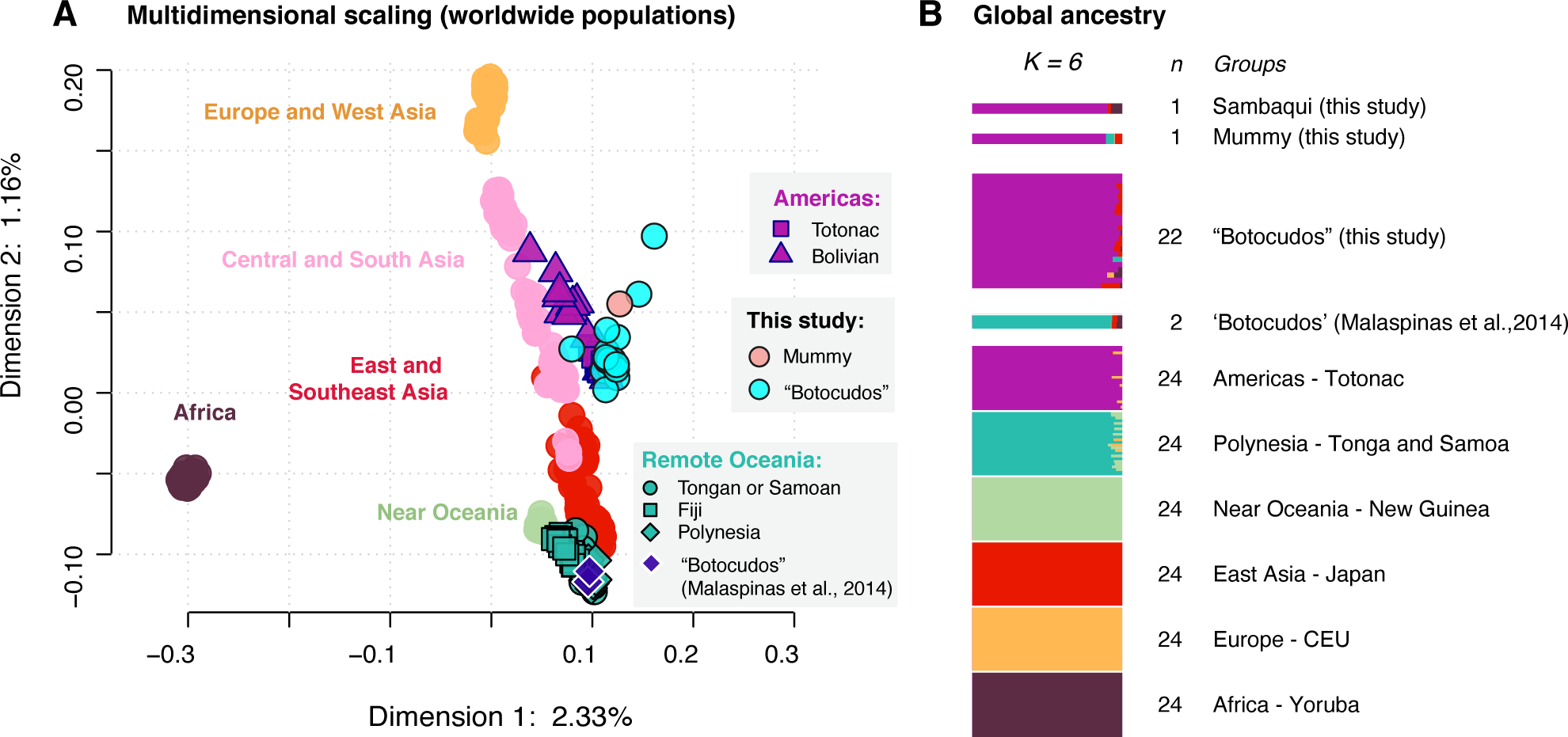
Genomic ancestry for the 24 sequenced individuals and 586 individuals from present-day worldwide populations (transversions only). (**A**) Multidimensional scaling combining 21 “Botocudos” and the mummy (MN01943). The individuals MN00010 (“Botocudo”) and MN01701 (Sambaqui) are not included in this plot as there was no overlap in SNPs between them and at least one other ancient individual (see “Multidimensional scaling” in Materials and Methods). MDS analyses including one ancient individual at a time for all individuals sequenced in this study can be found in Datasets S5 and S6. ( **B**) Admixture proportions estimated using NGSadmix (Skotte et al., 2013) for six groups (Yoruba, CEU, Japanese, New Guineans, Tongan and Samoans, and Totonacs), and fastNGSadmix (Jørsboe et al., 2017) for the 22 “Botocudos” from this study (see “Clustering analyses - Ancestry estimation” in Materials and Methods). Each horizontal bar represents the genome of a single individual, colored in proportion to their ancestry components. “Botocudos” are ordered by depth of coverage, with the individuals with the highest depth at the top. The number of individuals per group is indicated next to the bars. Ancestry proportions were estimated in two steps. First, NGSadmix was used to estimate the allele frequency proportions and ancestry in the following six (K=6) groups: Yoruba, CEU, Japanese, New Guineans, Tongans and Samoans, and Totonacs. Those inferred ancestry proportions and the allele frequencies were then used as input for fastNGSadmix to infer the ancestry of the following individuals: 22 “Botocudos”; MN01701 (Sambaqui); MN1943 (mummy); the two Polynesian “Botocudos” published in Malaspinas et al., 2014. See “Reference panels and dataset merging” in Materials and Methods for a description of the panels “MDS_NGSadmix_Wollstein_Xing_Malaspinas” and “NGSadmix_fastNGSadmix_Wollstein_Xing” used in this analysis.

To assess whether there is evidence of gene flow from populations outside the Americas, a two-step clustering analysis was performed with the tools NGSadmix (Skotte et al. 2013) and fastNGSadmix (Jørsboe et al. 2017). This analysis allowed us to infer clusters of ancestries predominant in six populations representing different geographical regions: Yoruba (Africa), CEU (Utah residents with Northern and Western European ancestry), Japanese (East Asia), Papuans (New Guinea), Tongans and Samoans (Polynesian Islands), and Totonacs (Americas). After identifying the allele frequencies per cluster (assuming six clusters), we inferred admixture proportions in the two Polynesian-“Botocudos”, the 22 “Botocudos”, the mummy and the Sambaqui (Fig. 2B).

All the newly sequenced individuals with a DoC above 0.005× (n = 19 individuals) showed >90% Indigenous American ancestry with the remaining ancestry being assigned mainly to the East Asian cluster (Fig. 2B). In these individuals, the estimates for Polynesian ancestry ranged between 0.0% (standard deviation, SD: 0.0%) and 4.2% (SD: 3.5%). For the remaining 5 individuals with DoC below 0.005×, the Indigenous American ancestry ranged between 83.7% (SD: 6.5%) and 96.2% (SD: 4.8%) with the remaining ancestry assigned to East Asian (maximum assigned: 11.2%, SD: 11.3%), African (max.: 6.9%, SD: 5.7%), European (max.: 6.1%, SD: 4.5%), and Polynesian clusters (max.: 4.2%, SD: 5.5%) (Fig. S6 and Dataset S7). These higher proportions of non-Indigenous American ancestry can be attributed to noise due to the low-coverage of those genomes. For instance, despite using a panel with twice as many sites, the authors of fastNGSadmix (Jørsboe et al. 2017) also report a decrease in Indigenous American ancestry for a Yjxa individual after downsampling the genome to low DoC. In summary, our results are consistent with a scenario in which the 22 herein sequenced “Botocudos” as well as the mummy and the Sambaqui individuals did not receive recent gene flow from populations outside the Americas. Thus, we assume that the previously documented (Malaspinas et al., 2014) presence of people of Polynesian ancestry among “Botocudos” was rare.

### Relationship to Indigenous Americans

Having established that the 24 individuals in this study are Indigenous Americans, we investigated their relationship to other ancient and present-day populations from the Americas. Based on cranial morphology, it has been suggested that the “Botocudos” are direct descendants of the Lagoa Santa people (Imbelloni 1938; Lacerda et al. 1876; Paul 1942; Pucciarelli et al. 2003) while historical records suggest that the “Botocudos” could be related to other Jê-speaking populations (Ehrenreich 2014; Paraíso 1992).

### Uniparental markers

The Y-chromosome and mitochondrial haplogroups of the individuals studied here (Table 1) are characteristic of Indigenous American populations (Kivisild 2017; Alves-Silva et al. 2000; Ramallo et al. 2013; Colombo et al. 2022). The Y-chromosome haplogroups of six out of eleven “Botocudo” males could be typed and were determined to be Q1a2a1-L54 and Q1a2a1a1-M3 (Table 1). Regarding maternal lineages, the 16^th^ – 19^th^ century “Botocudos” were distinct from the older individuals sequenced in this study (Sambaqui and mummy). For the “Botocudo” individuals, all mitochondrial haplogroups (Table 1) were determined to be either C or a subclade of it (Table 1). In contrast, the 1,000-year-old mummy and the 2,000-year-old Sambaqui individual were found to harbor A2 and D4 haplogroups, respectively. All these haplogroups (A2, C and D4) are commonly found in present-day Brazilians and they are widespread across present-day and ancient Indigenous Americans.

### Outgroup-*f*3 statistics

We then identified the most closely related populations to the 22 “Botocudos” at the nuclear genomic level. For this purpose, we compared the newly sequenced “Botocudos” to previously published genomic data from Indigenous Americans (Moreno-Mayar, Vinner, et al. 2018; Posth et al. 2018; Raghavan et al., 2015; Skoglund et al. 2015) using outgroup-*f_3_* statistics of the form (Yoruba; “Botocudos”, Indigenous Americans).

Among all Indigenous American populations tested, we observe the lowest *f_3_* values (i.e., the lowest shared genetic drift) for ancient and present-day populations from Greenland (i.e., Saqqaq and East and West Greenlanders) (Figs. 3 and S7 – S9), indicating the lowest affinity to the “Botocudos”. This is expected since they represent relatively recent migrations to the New World Arctic dating to the last 6,000 years, when the Arctic started to be peopled (Raghavan et al., 2014). We then observe a gradient in *f_3_* values with low scores for populations in the north and higher scores for populations in South America, with the highest values observed for populations to the east of the Andes, including Brazil. When comparing to populations in Brazil, the “Botocudos” have the highest affinities with the Xavante (central Brazil), Kaingang (southern Brazil), Zoró (west Amazonia), Urubu Ka’apor (east Amazonia), and Arara (east Amazonia) peoples (Figs. 3 and S7 – S11). We note that the Jê-speaking Xavante people record the highest values for the outgroup-*f_3_* tests across different subset of SNPs (Fig. 3 and Figs. S10 and S11); however, these scores are not significantly different from those obtained for other populations in Brazil or South America (for instance, the Tupiniquim in East Brazil, the Kaingang, Zoró, Paiter and Yjxa, or even the Aymara from Bolivia). In summary, the 22 “Botocudos” present a close affinity to Indigenous populations in Brazil and surrounding territories. However, no Indigenous Brazilian population analyzed here is significantly more closely related to the “Botocudos” (Fig. 3).

**Figure 3.**
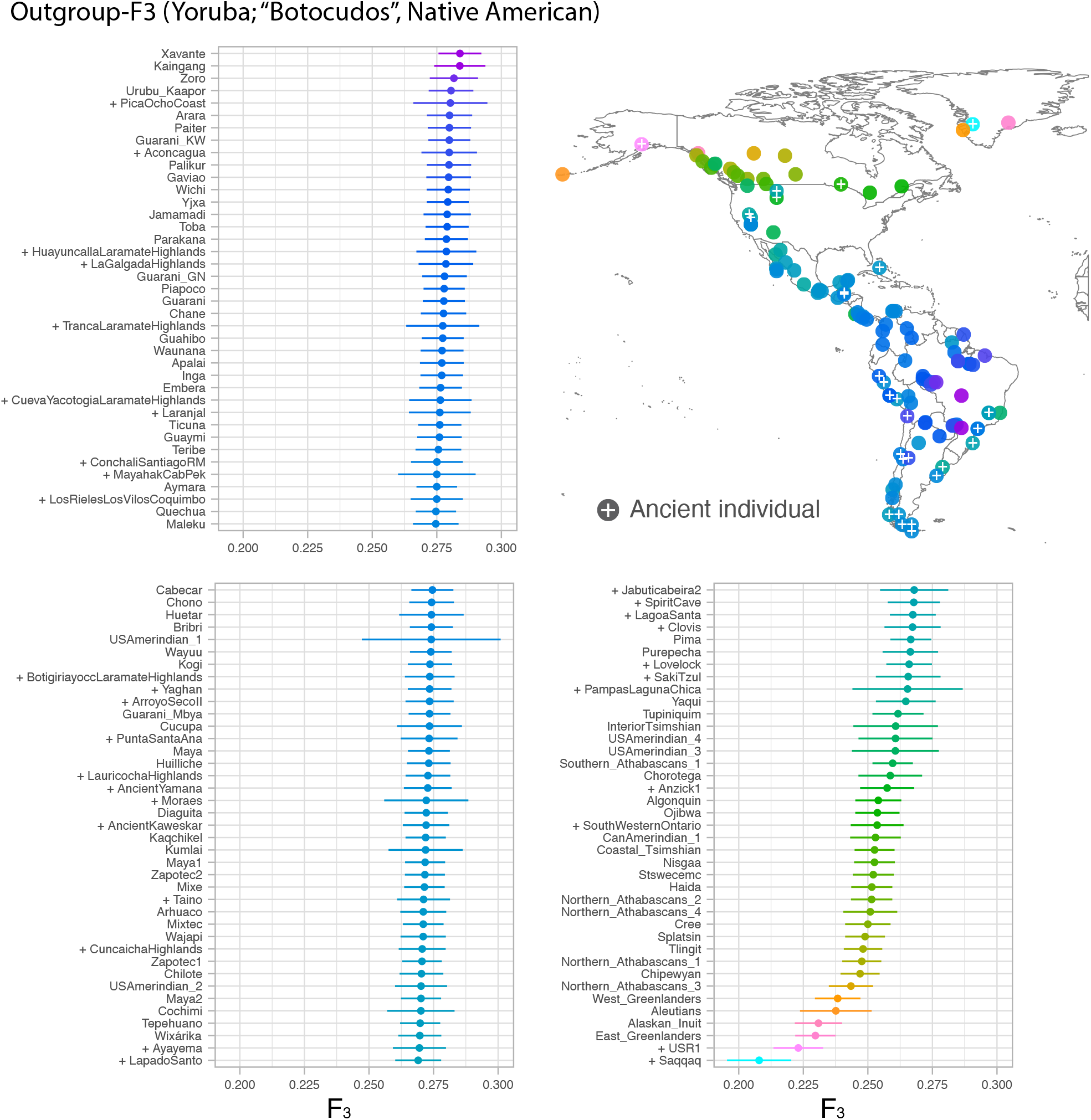
Shared genetic drift between the 22 “Botocudo” individuals and other Native Americans. X-axis: Outgroup-f3 value for the test *f*3(Yoruba; “Botocudos”, Native American) (Patterson et al., 2012); y-axis: Native American populations. Higher values correspond to higher similarity between the Native American population tested and the 22 “Botocudos”. To help with readability on the y-axis, the values are split into three panels. The values are ordered from highest to lowest (top to bottom and left to right). Ancient populations are labeled with a “+”. Error bars correspond to three standard errors. Locations of the Native American populations are displayed on top-right panel. Dots are colored according to the point estimate of the *f*3-test. Ancient populations or individuals are marked with a “+”. See “Reference panels and dataset merging” in Materials and Methods for a description of the panel F3_Raghavan_Skoglund_Castro_AncAmericas used in this analysis. The populations Paiter, Yjxa and Wixárika have been labeled as “Surui”, “Karitiana” and “Huichol”, respectively, in previous publications.

### Genetic structure in South America

To further investigate the relationship between the “Botocudos” and other Native American populations, we estimated admixture proportions with NGSadmix (Skotte et al. 2013) (Fig. 4). Using a panel enriched for Indigenous American populations and “outgroups”, we can identify different clusters outside and within the Americas. For instance, at K = 13 (Fig. 4)–the K at which we distinguish a Jê-related (Xavante) and a Guarani-related (Guarani Kaiowá) cluster–we observe four expected clusters for the outgroups (Yoruba, Papuans, French and Dai) and nine clusters within the Americas: Eskimo-Aleut and Na-Dene speakers (East- and West-Greenlanders, Coastal Tsimshian, Nisga’a, Tlingit, Haida and Chipewyan), Aridoamerican (Pima), Mesoamerican (Tepehuano, Mixe and Zapotec), Andean (Aymara and Quechua), Chibchan-Paezan (Cabecar), Yjxa (Tupi speakers), Paiter (Tupi speakers), Xavante (Jê speakers), and Guarani Kaiowá (Tupi speakers) (see Dataset DS10 for runs with 22 “Botocudos” and other K values with this panel).

**Figure 4.**
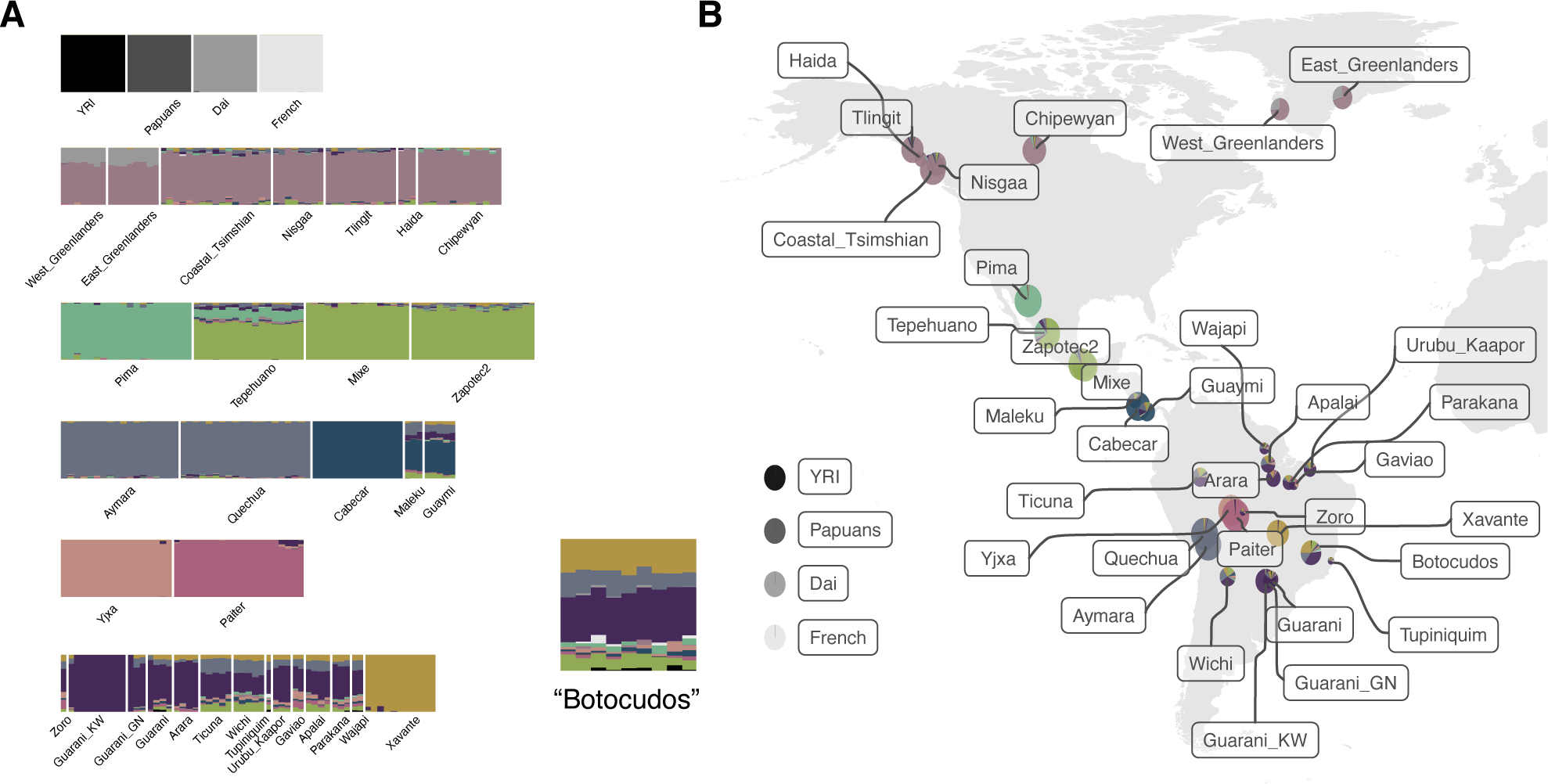
Admixture proportions for nine “Botocudos”, 34 Native American groups and four outgroups (K = 13). (**A**) Admixture proportions using a clustering analysis relying on genotype likelihoods (NGSadmix Skotte et al., 2013) estimated for a panel with ∼56,000 SNPs and assuming 13 ancestral components. Each horizontal bar represents the genome of a single individual, colored following their inferred ancestry proportions. Nine “Botocudos” from this study (the genomes with a depth of coverage above 0.1×) are included in this analysis and are ordered by depth of coverage, with the individual with the highest depth on the left (MN0008, MN00056, MN00013, MN0009, MN00118, MN00346, MN00067, MN00021 and MN00119). (**B**) Admixture proportions and geographical locations. Pie charts shows the average admixture proportions as inferred in (A). For the populations in the Americas, the pie charts are placed on their location on the map. The charts corresponding to YRI, Papuans, Dai and French are placed on the left. The radius of each pie chart is proportional to the sample size of the corresponding population. See “Reference panels and dataset merging” in Materials and Methods for a description of the panel “NGSadmix_Raghavan_Skoglund_Castro” used in this analysis. Plots for other K values and 22 “Botocudos” are shown in Dataset S10. The populations Paiter and Yjxa have been labeled as “Surui” and “Karitiana”, respectively, in previous publications.

At this K value (K = 13), the “Botocudo” ancestry is depicted as a mix of the Guarani-(purple) and Xavante-related (yellow) components, followed by the Andean- and Mesoamerican-related components (violet and green, respectively) in smaller proportions. Similarly, several other populations in Brazil (i.e., Ticuna, Urubu-Kaápor, Gavião, Apalai, Parakanã and Wajãpi) are also modeled as a mixture of the Guarani-, Xavante-, Mesoamerican- and Andean-related components. However, we observe that the “Botocudos” carry a larger proportion of the Jê-related Xavante component compared to these populations (see also Datasets DS11 – DS14 for runs with other panels).

We subsequently explored the relationship between the ancient “Botocudos” and other present-day Indigenous populations from Brazil using admixture graphs (Patterson et al. 2012). First, we considered a tree similar to that in (Castro e Silva et al. 2020), where Tupi-speaking groups (Yjxa, Paiter, Urubu Ka’apor and Guarani Kaiowá) form a clade to the exclusion of the Jê-speaking Xavante. We grafted the 16× ancient “Botocudo” (MN0008) to all possible branches of this tree and found that the topology where the ancient “Botocudo” is most closely related to the Jê-speaking Xavante (Fig. 5A, Fig. S12) resulted in the best fit score and the lowest *f_4_*-residual Z-scores. By contrast, when the “Botocudo” was added to the Tupi clade, all trees yielded higher residuals (Z-score > 3, Fig. S12) between the observed and predicted *f_4_*-statistics, suggesting a poor fit when adding the “Botocudo” to this clade. These results were consistent when we grafted the ancient individual to a different starting graph with a significantly better fit score (p-value > 0.05), where, for the base tree, Tupi-Guarani-speakers (Urubu Ka’apor and Guarani Kaiowá) are modelled as a mixture of populations related to the Tupi-speaking Paiter and the Jê-speaking Xavante (Fig. 5B, Fig. S12).

**Figure 5.**
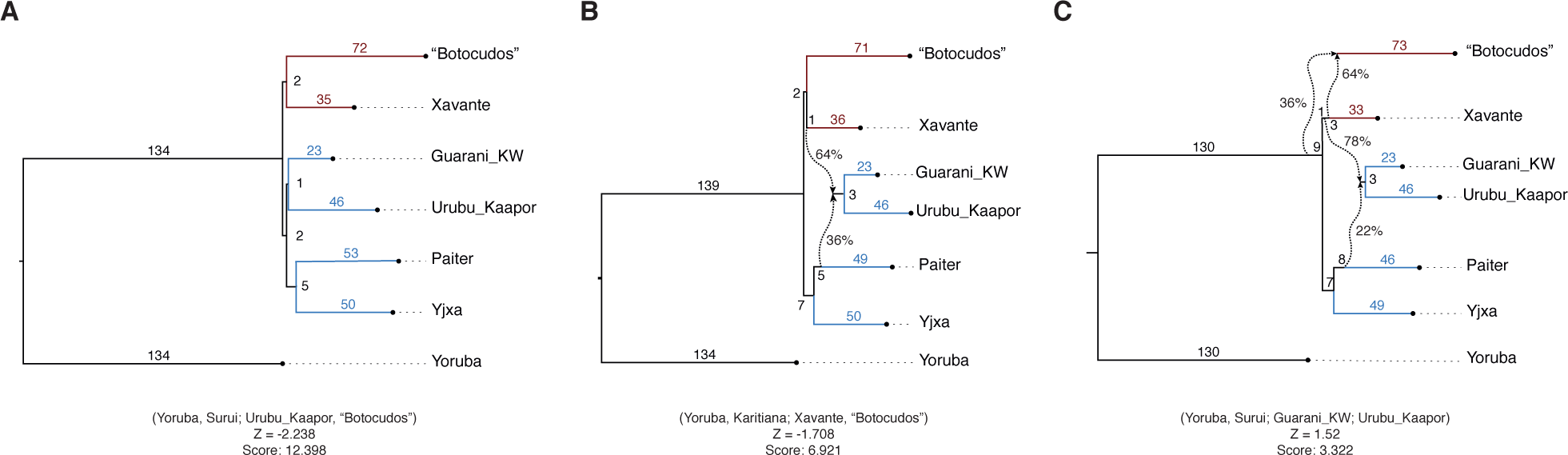
Admixture graphs modelling the relationship between the “Botocudos” and other indigenous peoples from Brazil. We considered a graph including Jê- (Xavante and “Botocudos”, colored in red) and Tupi-Guarani-speaking (Paiter, Yjxa, Urubu_Kaapor and Guarani_KW, colored in blue) groups. We show the best-fitting graph resulting from adding the 16× “Botocudo” genome (USER treated libraries, trimmed data and diploid genotypes) to all possible branches of (**A**) a simple tree similar to that in (Castro e Silva et al., 2020), and (**B**) a graph where Guarani_KW and Urubu_Kaapor bear Xavante- and Paiter-related admixture. In (**C**), we show the best-fitting graph where we include the “Botocudo” individual as an admixed leaf to the graph in (B). For each panel, we indicate below the graph the combination of four populations that give rise to the worst residual between the observed and the expected *f*-statistics, the Z-score corresponding to that residual and its fit score. The branches leading to Native American populations are colored according to their linguistic family (Jê in red and Tupi in blue). In Fig. S12 we also show graphs with suboptimal fit scores for each model. See “Reference panels and dataset merging” in Materials and Methods for a description of the panel “qpGraph_Skoglund_Castro” used in this analysis. The populations Paiter and Yjxa have been labeled as “Surui” and “Karitiana”, respectively, in previous publications.

We then assessed if modelling the “Botocudo” as a mixture of two different branches in the graph yielded a significantly better fit. We found that models where the “Botocudos” carry Xavante-related ancestry and a contribution from an outgroup to the Indigenous peoples from Brazil provided significantly better fit scores (Fig. 5C, Fig. S12). We thus proceeded to assess whether this “outgroup admixture” represents a population east or west of the Andes, or basal to South American populations. As our initial graph included only Indigenous South Americans located on the East of the Andes, we first grafted the Andean Aymara to all possible branches of the best-fitting graph. However, there were different models that were similarly good (as determined by the fit scores and residuals) for the placement of the Aymara on the graph (Fig. S12). Thus, in this case we could not find a model that clearly represented a better fit over the others. We surmise that the lack of power to find a distinctly good placement of the Aymara in the topology is related to the resolution of the dataset and the radiation-like split patterns of South American populations after initial peopling of the continent (Moreno-Mayar, Vinner, et al. 2018).

Altogether, these results suggest that, of all the Indigenous peoples for which genetic data are analyzed here, the “Botocudos” showed closest genetic relationship with the present-day Jê-speaking Xavante. However, we note that establishing a direct connection between populations is challenging since—similar to other populations East of the Andes—the “Botocudos” and the Xavante have experienced substantial population-specific drift suggesting they diverged a long time ago. Furthermore, the genetic relationship between the Xavante and “Botocudos” is not captured by simply assuming a split between sister populations, as the ancestry of the latter is better explained by including a component originating from a Native American outgroup.

### Population and social structure among the “Botocudos”

As pointed out above, the “Botocudo” label was given to different Indigenous Brazilian nations, therefore the individuals sequenced in this study may have belonged to one or more nations. For example, the label was used by German explorers to describe the social organization of Indigenous populations living next to the rivers Jequitinhonha and Doce (separated by about 400 km) in the beginning of the 19^th^ century (Ehrenreich 2014; Paraíso 1992; Wied 1820). However, it is unknown whether this vast area corresponds to their original geographical range around the time of European colonization, or if they engaged in large migrations due to the occupation of the territory by the European colonizers (Ehrenreich 2014; Paraíso 1992). Social practices and customs related to some peoples referred to as “Botocudos” have been documented, but it is unclear if those apply to all tribes under this denomination during the 18^th^ and 19^th^ century. For instance, several scholars report that the “Botocudos” were subsisting on a hunting-fishing-gathering strategy and were organized in small nomad or semi-nomad bands characterized by constant group fragmentation (Ehrenreich 2014; Paraíso 1992), possibly related to fission-fusion dynamics observed in other Indigenous South American populations (Neel et al. 1967). It has also been suggested that marriage between cross-cousins was preferred, while marriage between parallel cousins was forbidden, and exogamous marriages were allowed (Ehrenreich 2014; Paraíso 1992). To gain insight from a genomic perspective into the population diversity and size, as well as marital practices among the “Botocudos”, we inferred the levels of heterozygosity, runs of homozygosity and changes in effective population size (Ne) over time.

### Conditional heterozygosity

We assessed heterozygosity levels for all pairs of “Botocudos” and compared them to those of other worldwide populations, including ancient and present-day Native Americans (Fig. 6A). To reduce the effect of DNA damage, we restricted this analysis to sites that are heterozygous in an African genome, and we sampled one allele per individual in order to generate a pseudo-haploid genome (following Skoglund et al. 2014). The pseudo-haploid genomes of two individuals are then compared (thus reducing the impact of inbreeding within individuals) and the conditional heterozygosity is computed as the ratio of heterozygous sites to the number of sites with data for both individuals.

**Figure 6.**
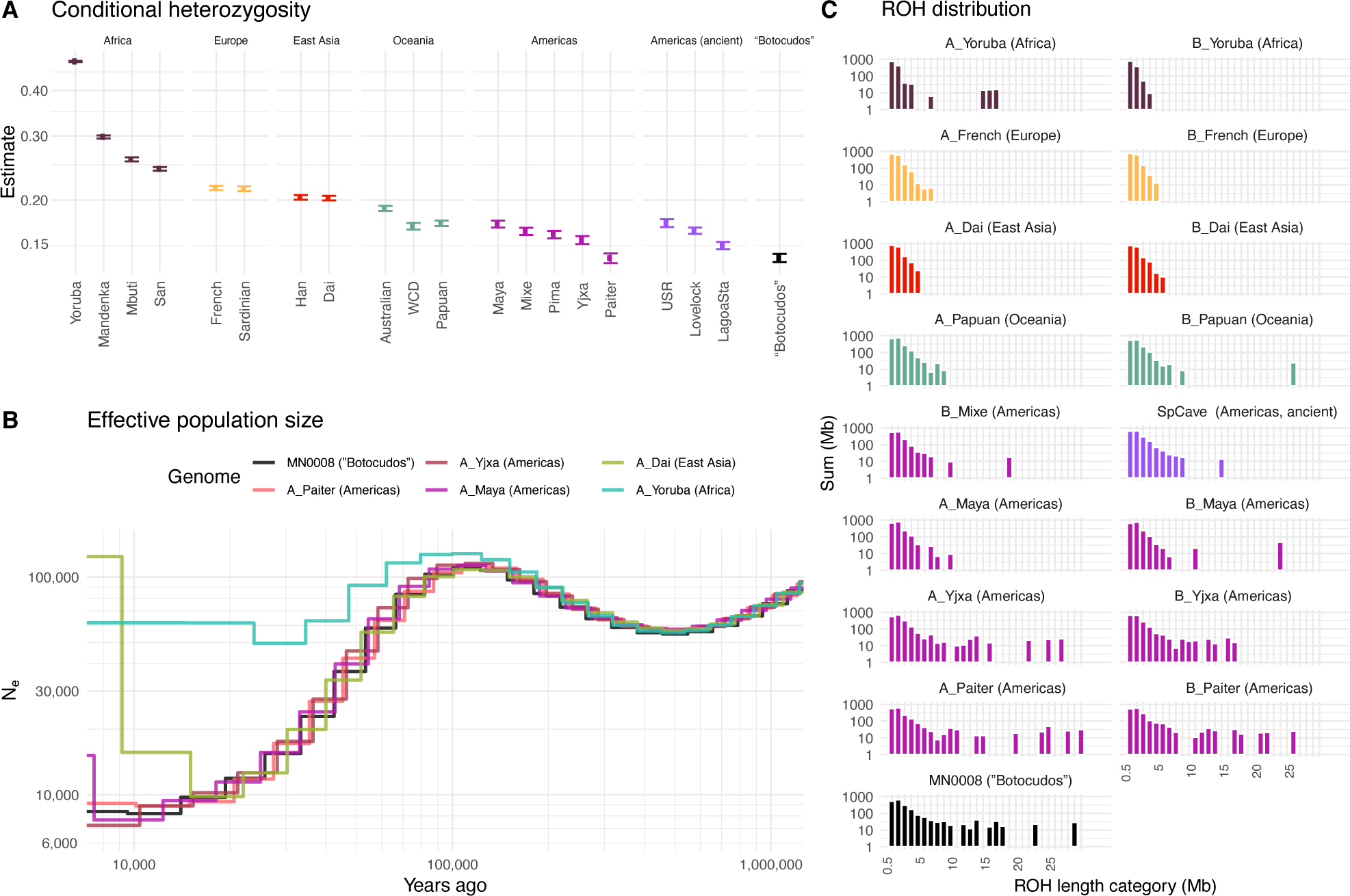
Heterozygosity, runs of homozygosity (ROH) and effective population size. (**A**) Heterozygosity values conditioned on heterozygous sites (transversions only) ascertained in a Yoruba genome. Exactly two individuals were selected for each population (x-axis), and an allele was sampled for each of them. “Botocudos” MN0008 and MN00056 were selected to be displayed in this figure. Each dot corresponds to the average estimate per pair of individuals, and the error bars represent the 95% confidence interval obtained from a jackknife resampling approach over 5 Mb blocks (Skoglund et al., 2014). Refer to Fig. S13 for the estimates with all pairs of “Botocudos”. ( **B**) Effective population size over time inferred with PSMC (Li & Durbin, 2011). The inference was done on genotype calls (transversions only) from a single genome per population. ROH and effective population size estimations for the “Botocudo” individual MN0008 were done on genotype calls from trimmed reads (2 bp on each end) from the USER-treated library only. (**C**) Runs of homozygosity distribution per individual. X-axis: length category; y-axis: sum of ROH lengths. ROH were inferred with PLINK (Purcell et al., 2007) on genotype calls (transversions only) from genomes with a minimum depth of coverage of 15×. See “Reference panels and dataset merging” in Materials and Methods for a description of the panel “Conditional_heterozygosity” used in this analysis. The populations Paiter and Yjxa have been labeled as “Surui” and “Karitiana”, respectively, in previous publications.

Conditional heterozygosity was then computed for every pair of “Botocudos” (Fig. S13 shows the distribution of the conditional heterozygosity point estimates among pairs). The point estimates for the “Botocudos” range from 0.10 to 0.18 across comparisons, with an average value of 0.14 and no outliers. As expected, there is higher variation in the estimates for pairs of individuals including less data (i.e., lower-coverage genomes). Specifically, when considering only pairs of genomes with a depth of coverage above 2× (MN00013, MN00056 and MN0008), the heterozygosity range is reduced to values between 0.139 and 0.140.

The “Botocudos’” heterozygosity was then compared with that in worldwide populations including Africans, Europeans, Oceanians, and ancient and present-day Native Americans. For this comparison, medium to high depth present-day genomes were selected and compared to the two “Botocudo” individuals with the highest depths of coverage (MN0008 and MN00056, Table 1). As expected (Henn et al. 2016; León et al. 2018), the conditional heterozygosity values decrease as the distance from Africa increases, with present-day and ancient Indigenous Americans exhibiting the lowest values; in particular, the “Botocudos” have the lowest conditional heterozygosity worldwide together with the Paiter people, also from Brazil. This could suggest that the ancient “Botocudos” analyzed here had a small effective population size, similar to that of the present-day Paiter population.

### Effective population size

We inferred changes in effective population size with a pairwise sequentially Markovian coalescent approach (PSMC, Heng Li et al. 2011) to gain insights into the demographic history of the “Botocudos” through time. The genomes of one Yoruba (Africa), one Dai (East Asia), one Maya (Mesoamerica), one Yjxa (South America), one Paiter (South America), and the high-coverage (DoC = 16×) “Botocudo” individual (MN0008) were analyzed with PSMC (Fig. 6B). All genomes follow the same trajectory until 50,000 to 100,000 years ago (ya), when the Yoruba population maintained a relatively high Ne, while all other out-of-Africa genomes showed a population decline. This has been interpreted as an out-of-Africa migration bottleneck (Heng Li et al. 2011). Next, the effective population size of the East Asian population starts increasing from 20,000 to 10,000 ya in contrast to Indigenous American effective population sizes, which decrease during the same timeframe. Within the Americas, the population size of the Mesoamerican Maya starts increasing a few thousand years ago, as reported in previous studies (Bergström et al. 2020) but also at a timeframe where the method is limited by lack of resolution. In contrast, the effective size of the ”Botocudo” people follows a trajectory similar to that of the Amazonian Yjxa and Paiter, with an effective size that remains small (<10,000) during the last 20,000 years. We note, however, that recent inbreeding in the individual MN0008 would impact those estimates (see also below).

### Runs of homozygosity

Runs of homozygosity (ROHs) represent segments of a genome with contiguous homozygous states. ROHs are expected to arise more frequently under various scenarios, including reduction in population size (Kirin et al. 2010; Pemberton et al. 2012), endogamy (Kirin et al. 2010; McQuillan et al. 2008; Yengo et al. 2019), and natural selection (Pemberton et al. 2012). We inferred ROHs for the “Botocudo” MN0008 (DoC = 16×) and 14 other high-coverage genomes (DoC > 20×) from Africa, Europe, East Asia, Oceania, and the Americas to get a sense of the extent of homozygosity distribution in the genomes of the “Botocudos” compared to other populations.

The total lengths of ROHs segments in each length category for 15 individuals are shown in Fig. 6C. The out-of-Africa genomes have longer ROHs as reported before (Kirin et al. 2010; Pemberton et al. 2012), while present-day and ancient Native Americans have larger portions of their genome within ROHs longer than 5 Mb than Africans, Eurasians and Oceanians, possibly reflecting the serial population bottlenecks experienced by the first peoples to enter the Americas (Pemberton et al. 2012). Lastly, we observe even longer (>10 Mb) ROH in present-day South American (Yjxa and Paiter) groups and in the ancient “Botocudo” individual MN0008.

Comparisons of individuals from different subpopulations would lead to higher heterozygosity estimates (compared to that of the individual subpopulations). Yet the “Botocudos” show very low levels of heterozygosity without any outliers among estimates for pairs of individuals (Fig. 6A) and no ancestry clusters formed within the “Botocudos”, suggesting little differentiation between them, a single population or gene flow between subpopulations. Moreover, the “Botocudos” display a small effective population size and long ROHs in par with present-day Indigenous Brazilians (Fig. 6BC). Taken together, these results suggest that the analyzed “Botocudos” have a population and inbreeding history similar to those of the Amazonians Paiter and Yjxa. These two populations have experienced collapses in their population sizes following contact and introduction of pathogens (Fleming-Moran et al. 1991; Kanindé Associação de Defesa Etnoambiental et al. 2021; Mindlin 1985; Monteiro 1984; Velden 2004); regarding social practices, the Paiter and Yjxa also encourage both exogamic and cross-cousins marriages (Kanindé Associação de Defesa Etnoambiental et al. 2021; Storto et al. 2021). Note however that as the ROH and effective population size results for the “Botocudos” are based on a single individual (i.e., the high DoC genome of MN0008), more high coverage genomes and demographic modelling are needed to get a more comprehensive view on the demography of other ancient Indigenous Brazilians and exclude scenarios such as selection to explain the data.

### Prevalence of Australasian ancestry among South Americans

Recently, it has been shown that, compared to other Native Americans (like the Mixe from Mexico), some present-day Native Brazilians–including the Paiter, the Yjxa and the Xavante (Skoglund et al. 2015)–and peoples in the Pacific coast like Chotuna (Castro e Silva et al. 2021) share more alleles with

Australasians (e.g., Australians, Papuans or Onge from the Andaman Islands) than expected by chance. As this increased affinity is not consistently widespread in large regions of the Americas, it has been hypothesized that this signal could be attributed to population structure in the ancestral population that peopled the Americas (Moreno-Mayar, Vinner, et al. 2018; Raghavan et al., 2015; Skoglund et al. 2015) with more recent gene flow having potentially reduced the signal in some populations (Moreno-Mayar, Vinner, et al. 2018). In agreement with an ancestral population structure hypothesis, this excess allele sharing was only replicated in some ancient individuals in Brazil, such as the 10,000-year-old Lagoa Santa individual (Moreno-Mayar, Vinner, et al. 2018), but not in others, such as the 10,000-year-old Lapa do Santo individuals (Posth et al. 2018).

To assess whether Australasians represent a strict outgroup to 19^th^ century Brazilian “Botocudos” and the Mixe from Mexico, D-statistics (Green et al. 2010) of the form D(H_1_, Mixe; H_3_, Yoruba) based on whole genomes were computed (Fig. 7). These statistics allowed us to test whether the Native American in H_1_ (e.g., the “Botocudos”) share more or less alleles with H_3_ (an Australasian or a Eurasian population) than the Mixe does. For the tests that we implemented, H_1_ was a Native American population (Brazil: “Botocudos”, Paiter, Yjxa, Lagoa Santa; Bolivia: Aymara; Mexico: Wixárika), and H_3_ an Australasian (Australian, Papuan, historical Andamanese) or a Eurasian (Han or French) population. To reduce the impact of post-mortem molecular damage, we adopted three strategies: (i) USER-treated data were used when possible for the ancient individuals, (ii) the data were restricted to sites containing transversions only and (iii) the D-statistics were corrected by taking into account the inferred error rates for the ancient and present-day genomes (Soraggi et al. 2018).

**Figure 7.**
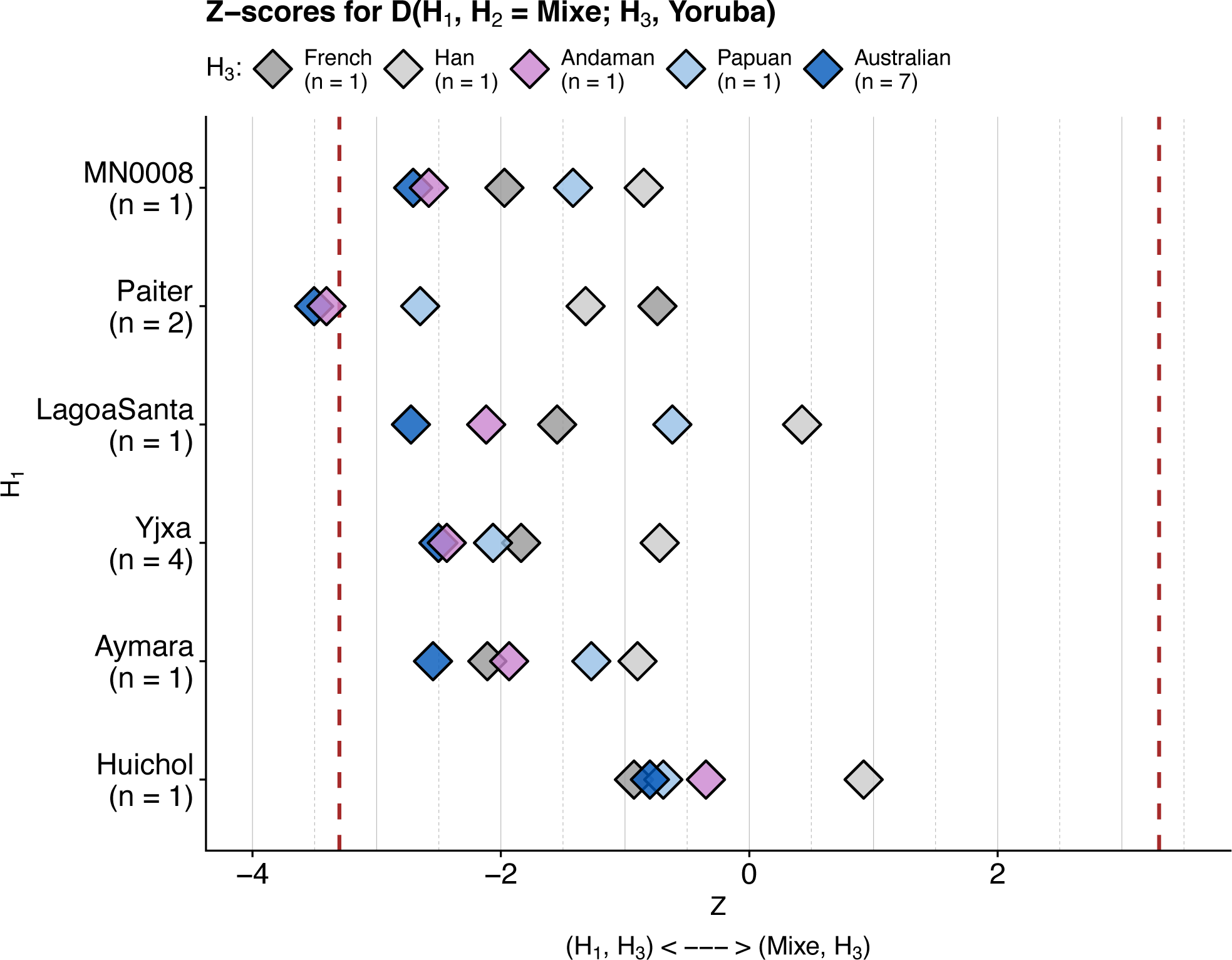
Z-scores for error-corrected D-statistics D(H_1_, H_2_ = Mixe; H_3_, Yoruba). Y-axis: population used in the test as H_1_. Color code: population used in the test as H_3_. When H_3_ is closer to H_1_ than to Mixe, the Z-scores become negative (left-hand side of the plot). In contrast, positive Z-scores (right-hand side of the plot) indicate a stronger proximity between Mixe and H_3_ than between H_1_ and H_3_. The number of genomes per population is indicated between parentheses. D-statistics were corrected according to the error rates estimated per genome following (Soraggi et al., 2018). Z-scores were obtained after a jackknife resampling approach of D-statistic for 5 Mb-blocks. Vertical dotted lines are placed at |3.3| values and correspond to a *p*-value of approximately 0.005 (Green et al., 2010). For the “Botocudo” individual MN0008, only reads from the USER-treated library were analyzed. Reads from ancient genomes were trimmed either 2 bp (MN0008) or 5 bp (LagoaSanta, Andaman) on each end. See “Reference panels and dataset merging” in Materials and Methods for a description of the panel “ErrorCorr_Dstat” used in this analysis. The populations Paiter and Yjxa have been labeled as “Surui” and “Karitiana”, respectively, in previous publications..

As in previous studies, the affinity to Australasian populations was stronger when considering the Paiter in H_1_ (Fig. 7), the latter population sharing significantly more alleles with Australasian populations (Australian and Andaman). For the 10,000-year-old Lagoa Santa and the present-day Yjxa and Aymara, we found a trend similar to the one reported in (Moreno-Mayar, Vinner, et al. 2018), in which the Z-scores associated with the D-statistics are more negative when considering an Australasian population in H_3_ than a Eurasian population in H_3_. This indicates that there are more shared alleles between Lagoa Santa and Australasians and between Yjxa and Australasians than between Australasians and Mixe. Yet, we cannot reject the null hypothesis (similar proportions of shared alleles between those Native Americans and Australasians) at a 0.001 level (the Z-score are higher than-3.3). In contrast, when placing the Wixárika from Mexico in H_1_, the tests including an Australasian population in H_3_ were of a similar magnitude than the tests with a Eurasian population in H_3_. This suggests that for the Wixárika, indeed, Australasian gene flow cannot be detected when comparing to Mixe.

The Z-scores for the D-statistics with the “Botocudo” MN0008 (Fig. 7) as well as the combined 22 “Botocudos” (Fig. S14) in H_1_ are similar to those seen for Yjxa and Lagoa Santa in H_1_ (Fig. 7), suggesting that, similarly to Yjxa and Lagoa Santa, the “Botocudos” also share more alleles with Australasian than the Mixe does. In contrast, this asymmetry between the “Botocudo” and Mixe is not as marked when considering a Eurasian population in H_3_ (the Z-scores are lower for H_3_ = Australasian than for H_3_ = Eurasian). Yet, as for the Yjxa, the D-statistic values are not significant at a 0.001 level.

“Botocudo” populations have been thought to represent a case of long-standing genetic continuity bearing a strong craniometric resemblance to that of Australasians and to that of the Lagoa Santa people (Strauss et al. 2016; Neves 2017; Pucciarelli et al. 2003). We find that the “Botocudos” and the 10,000-year-old Lagoa Santa are similarly related to Australasians, but not more so than present day Native Brazilians. Overall, our results are in line with previous studies, as we observe a trend for some ancient and present-day South Americans to present a higher affinity to Australasians when comparing them to Mesoamericans (like Mixe). Likewise, this affinity to Australasians is higher than that to Eurasian populations (like the French and the Han Chinese). Yet, these results are only significant for the Paiter population. It could well be that a subtle signal is there for the “Botocudos” as well (and also the Yjxa and Lagoa Santa, as reported before), but that we lack statistical power for this test. Alternatively, the signal could truly be absent in the “Botocudos”. This suggests that either the craniometric resemblance to Australasians is due to chance or that the “Botocudos” are also the descendants of a structured population with close ties to Australasia as has been suggested for present-day Native Brazilians.

### Screening for ancient bacteria and viruses

Studies describing the microbial and viral diversity among pre- and post-European contact people in the Americas are scarce, but they can inform us on the microbes that were commonly circulating in the past and potentially played an important role in an individual’s health (Barquera et al. 2020; Bos et al. 2014; Guzmán-Solís et al. 2021; Bravo-Lopez et al. 2020; Vågene et al. 2018). We aimed at discovering bacteria and viruses that could have infected any of the 24 sequenced individuals by analyzing the shotgun data generated for them. This approach allowed us to identify several candidate disease-causing agents, and to reconstruct whole genomes for the most abundant bacteria and viruses with molecular signature typical of ancient DNA.

### Presence of ancient bacteria associated with periodontal disease

We used Kraken2 (Wood et al. 2019) to carry out a metagenomic characterization of the non-human reads by identifying ancient human bacteria that might have infected the 24 individuals (Dataset S15). Since all but two samples were teeth, we decided to focus on those associated with the aerodigestive tract (Escapa et al. 2018), and identified eight species: *Acinetobacter lwoffii*, *Actinomyces* sp. oral taxon 414, *Anaerolineaceae* bacterium oral taxon 439, *Comamonas testosteroni*, *Fusobacterium nucleatum*, *Pseudomonas stutzeri*, *Stenotrophomonas maltophilia* and *Tannerella forsythia*. In agreement with previous studies (Mann et al. 2018; Philips et al. 2017; Bravo-Lopez et al. 2020), we identified DNA from oral microbes present in high abundance (>10% of reads) in the dentin of the 20 out of 22 individuals for which teeth were sampled (Dataset S15). The two pieces of skull (MN0008 and MN00019) and the remaining two teeth samples (MN0003 and MN00064) also had hits to the eight oral bacteria, but in lower proportions (less than 2% of their reads). Among the eight oral bacteria, *Anaerolineaceae* bacterium oral taxon 439, *Actinomyces* sp. oral taxon 414 and *Tannerella forsythia* have been previously reported in archeological remains from Europe and the Americas (Eisenhofer et al. 2020; Kazarina et al. 2021; Wada et al. 2018; Ottoni et al., n.d.; Velsko et al. 2019; Warinner et al. 2014; Bravo-Lopez et al. 2020).

We then reconstructed the genomes of each of those eight bacteria, attaining different read depths in the 24 individual samples (Dataset S15) as follows: *A. lwoffii* (0.0006×-51.5×; n = 4 above 1×), *Actinomyces* sp. oral taxon 414 (0.004×- 8.2×; n = 2 above 1×), *Anaerolineaceae* bacterium oral taxon 439 (0.0003×- 30.0×; n = 2 above 1×), *C. testosteroni* (0.003×- 49.0×; n = 3 above 1×), *F. nucleatum* (0.0005×- 4.2×; n = 1 above 1×), *P. stutzeri* (0.008×- 3.9×; n = 6 above 1×), *S. maltophilia* (0.008×- 41.1×; n = 6 above 1×) and *T. forsythia* (0.0004×- 23.2×; n = 1 above 1×). To verify the authenticity of the twenty-five reconstructed bacterial genomes with a DoC above 1× (Dataset S15), we examined their damage patterns and coverage evenness. We observed that the deamination rates at the end of the reads were relatively high (10% to 17%) for the three taxa belonging to genera isolated mostly from the oral cavity in humans (*F. nucleatum, T. forsythia*, and *Anaerolineaceae* bacterium oral taxon 439; Escapa et al. 2018; Hofstad 2006; Beall et al. 2018), although the breadth of coverage (BoC) was variable, with 43% to 94% of the genome covered at least once. The remaining five taxa showed lower deamination rates (between 2% and 5%) on the terminal bases, and varied BoC (15% to 83%). These five taxa belong to the *Acinetobacter*, *Actinomyces*, *Comamonas*, *Pseudomonas* and *Stenotrophomonas* genera, which encompass several species commonly found in soil and aqueous environments (Towner et al. 1991; Willems et al. 2006; Palleroni 2010; Ryan et al. 2009). Therefore, further analyses would be required to disentangle endogenous microbial genomes from those of environmental microbes. Future phylogenetic analyses of these and other candidate species identified in the 24 individuals would help elucidate patterns of molecular diversification in the individual taxa.

### Presence of ancient human Parvovirus B19

Following Mühlemann et al., (2018), we screened the non-human reads of all 24 sequenced individuals for ancient viruses using the classification tool DIAMOND (Buchfink et al. 2014)and the NCBI viral proteins database. Fifty viruses (Dataset S16) that are known to infect humans (VirusHost database, Mihara et al. 2016) were identified. We then mapped all non-human reads to each of the 50 viral reference genomes with BWA (Li et al., 2009) and recovered different fractions of each viral genome, as indicated by their breadths of coverage (Dataset S17). As false positives are expected to be common in ancient virome studies (Arizmendi Cárdenas et al. 2021), we further analyzed the 15 viruses for which BoC was above 1% after mapping (Fig. 8A and Dataset S17). Based on genome coverage, damage patterns and read length distribution, we inferred the presence of ancient human parvovirus B19, but were unable to confirm the remaining 14 putative ancient viruses due to low read counts.

**Figure 8.**
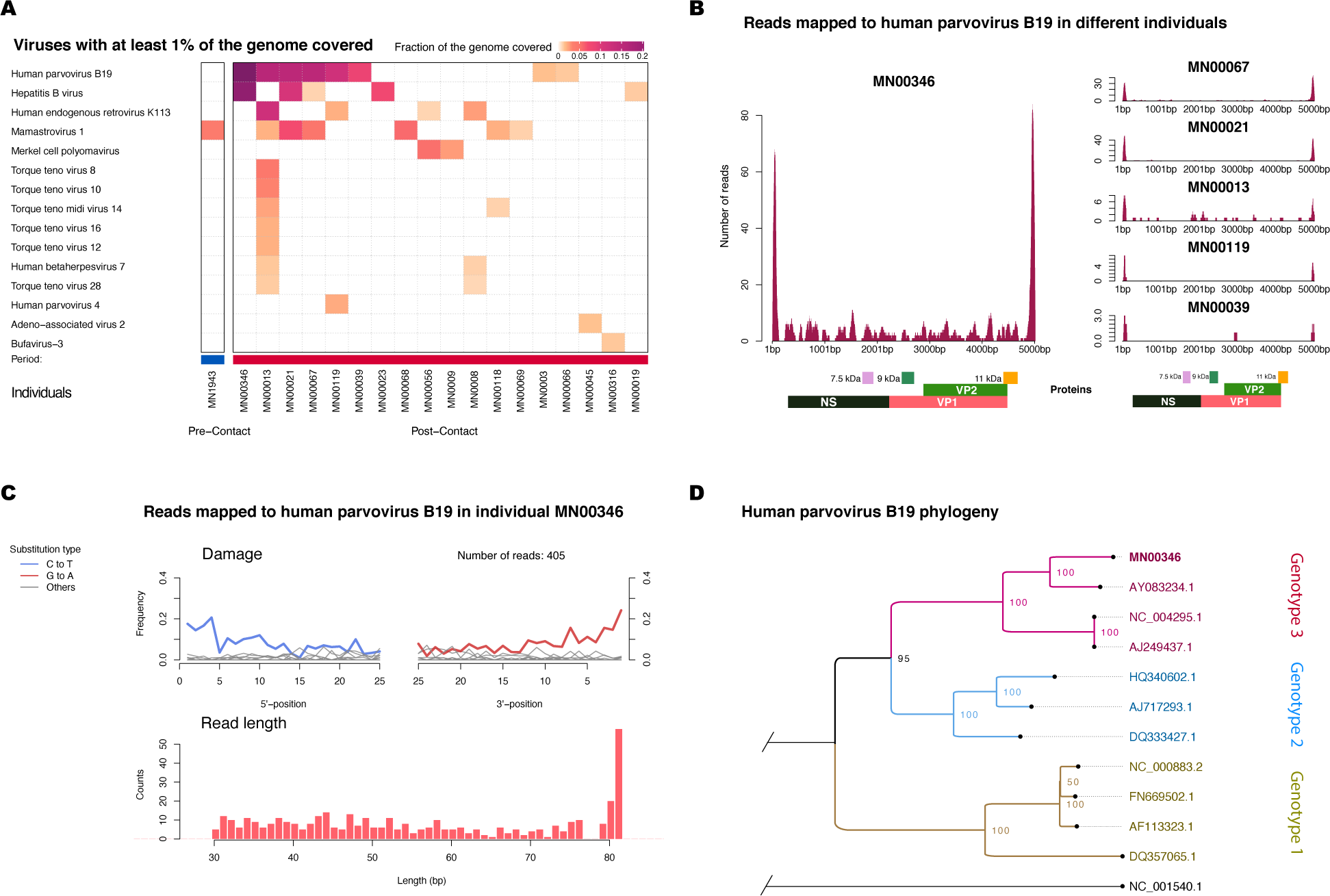
Top viruses detected in the sequenced data and human parvovirus B19 genome reconstruction. (**A**) Heatmap depicting the fraction of the viral genomes covered at least once (breadth of coverage, BoC) after mapping the non-human reads to the virus reference sequences. Each column represents the data for one individual and each row represents the viral genome used as a reference. Note that five “Botocudos” and the Sambaqui individual are not shown as none of their viruses had more than 1% of their genome covered by one read. (**B**) Genome coverage along the human parvovirus B19 genome (AY083234.1, genotype 3) per individual. For these six individuals, parvovirus B19 was the virus with the highest BoC. The title of each subplot indicates the individual’s ID. Boxes under the plots represent the viral proteins of the human parvovirus B19. At the end of the genome (3’ and 5’), the regions that are not annotated include the inverted terminal repeats. The higher achieved depth of coverage at the beginning and end of the genome can be explained by the palindromic nature of these regions. (**C**) Damage plot (top) and read length (bottom) for the MN00346 reads mapping to the human parvovirus B19 (AY083234.1). (**D**) Neighbor-joining tree of the consensus sequence obtained from the MN00346 reads mapping to AY083234.1 (labeled as MN00346 in the tree), together with ten human parvovirus B19 sequences representing three different genotypes (3 sequences per genotype), including the human parvovirus B19 sequence used in DIAMOND’s database (NC_000883.2, genotype 1), and an outgroup sequence (bovine parvovirus, NC_001540.1). Bootstrap values were obtained by running 100 replicates and are indicated at each node.

Human parvovirus B19 is responsible for fifth disease in children, persistent anemia in immunocompromised patients, transient aplastic crises, hydrops fetalis in pregnant women, and arthropathy (Qiu et al. 2017). This DNA virus has been previously identified in aDNA studies (Guzmán- Solís et al. 2021; Mühlemann, Margaryan, et al. 2018). In our analyses eight “Botocudo” individuals had between 1 and 726 sequencing reads mapped to the parvovirus genome (NC_000883.2, genotype 1, Dataset S17), corresponding to a BoC between 1.5% and 24.8% (Fig. 8A). We detected a higher DoC (for AY083234.1, genotype 3, see below) at the 3’- and 5’-termini of the viral genome for six out of eight individuals (MN00346, MN00067, MN00021, MN00013, MN00119, MN0039), as expected for authentic Human parvovirus B19 genetic data, since the termini are known to be “hairpin” regions, formed by double-stranded DNA (Fig. 8B, see also Guzmán-Solís et al. 2021). To further support the authenticity of the ancient viral DNA data, we evaluated the read length distribution and observed short mapped reads, as expected for degraded aDNA molecules (see Fig. 8C for MN00346). Among the individuals with human parvovirus B19, MN00346 is the individual with the highest number of reads (namely 405 reads, reaching an average DoC of 4.5×). Furthermore, the characteristic C-to-T and G-to-A aDNA damage substitutions at the read termini were around ∼20%, (Fig. 8C), making it the most promising candidate for further analyses. We thus used MN00346 data to reconstruct an ancient parvovirus genome.

To assess the genotype of the ancient parvovirus recovered from MN00346, the non-human reads were mapped independently to 11 parvoviral genomes as in (Mühlemann, Margaryan, et al. 2018). A consensus sequence was built with the reads mapped to the genome with the highest breadth of coverage (84%, genotype 3, AY083234.1) and, together with the 11 previously published parvovirus genomes, a neighbor-joining tree was reconstructed. We observed that the MN00346 consensus sequence clustered with genotype 3 (Fig. 8D), suggesting that the parvovirus that infected MN00346 belonged to this genotype. Genotype 3 was previously observed in other post-contact age individuals in the Americas (Guzmán-Solís et al. 2021) and is considered today as endemic to Ghana (although it has been reported in different countries, Qiu et al. 2017). Following (Guzmán-Solís et al. 2021), we hypothesize that it might have been introduced into the “Botocudo” population in Brazil upon European colonization, possibly through the transatlantic slave trade.

### Stable isotope dietary analysis

Diet can vary across individuals and populations due to a number of factors including their customs, the environment and individual preferences. The comparative study of stable isotopes such as δ^13^C and δ^15^N can inform us on whether an individual’s diet was constituted mainly of C3 (such as rice, wheat, manioc and yam) or C4 (such as maize, sugar cane and sorghum) plants and give us an idea on the levels of animal protein consumption. In South America, Indigenous peoples had access to both C3 (manioc, yam) and C4 (maize) local plants (Scheel-Ybert et al. 2003; Wesolowski et al. 2010). In central-eastern Brazil, several “Botocudo” peoples subsisted on a fisher-hunter-gathering strategy (Ehrenreich 2014; Paraíso 1992), until some of them were forcibly relocated to European-founded settlements during the European colonization (Paraíso 1992), where they were given access mainly to corn and fish (Ehrenreich 2014; Oliveira 2016).

Carbon and nitrogen stable isotopes were measured in the collagen of the Sambaqui (n = 1) and the “Botocudos’” (n = 14) permanent teeth and contrasted to publicly available data (Dataset S18). The datasets spanned different timepoints (from the Early Holocene to present day), archaeological and present-day contexts, and regions in Brazil. The δ^13^C and δ^15^N values for the 2,000-year-old- Sambaqui individual (MN01701) were high and similar to those of published 3,000-1,000 yo fisher-gatherers buried in coastal sambaquis in Santa Catarina state, indicating a high consumption of marine fauna. In contrast, the isotope values for the “Botocudos” were quite distinct from those of the coastal fisher-gatherers, with much lower δ^13^C and δ^15^N values. Their δ^13^C values were compatible with a diet based mainly on C3 plants, similar to those of the mid-Holocene riverine hunter-gatherers from São Paulo, and the Early Holocene hunter-gatherers from Minas Gerais (Fig. S15). Furthermore, on average, the “Botocudos” presented significantly higher δ^15^N values than the Early and mid-Holocene hunter-gatherers and the present-day individuals in Rio de Janeiro and Brasília (Tukey’s range test, p-value < 0.05). Overall, this suggests that the diet of the “Botocudos” did not include a high proportion of marine fauna, that it was mostly based on C3-plants (as attested by the low δ^13^C values) and that they had access to more animal protein sources than Early and mid-Holocene hunter-gatherers and present-day individuals (as attested by their δ^15^N values).

### Conclusions and future directions

The *Museu Nacional*, founded in 1818, is one of the oldest and largest museums in Latin America. In September 2018, the Museum was consumed by a tragic fire, and the human remains preserved in its anthropological collection were largely lost. Our results represent the characterization of a part of the Museum’s anthropological collection recovered prior to the fire, including Indigenous individuals from Eastern Brazil, in particular the “Botocudos”. The term “Botocudo” is a generic name that was first given by Portuguese colonizers to Indigenous peoples wearing wooden disks as facial ornaments, and “Botocudos” likely comprised several distinct populations whose trace was lost in the historical record. Nowadays, some Indigenous people identify as “Botocudo” (IBGE 2010) yet very little was known about the genomic history different “Botocudo” peoples as only two genomes of Polynesian ancestry had been sequenced so far. Here, we present isotope, radiocarbon date, and shotgun sequencing data for 22 post-contact “Botocudos” (including a 24× coverage whole genome sequence), in addition to a pre-contact sambaqui-associated individual and a 1,000-year-old mummy.

Out of the 35 “Botocudo” individuals previously housed at the National Museum of Rio de Janeiro, 22 were found to be of Indigenous American ancestry (present study), and two, slightly older, were found to be of Polynesian ancestry (Malaspinas et al., 2014). At this point, we are unable to further disentangle the different scenarios proposed in 2014 to explain the presence of Polynesian individuals in the “Botocudo” collection. We have no further evidence to corroborate or disprove that the two previously sequenced Polynesian-“Botocudo” were either the result of transpacific contact between Polynesia and the Americas, a European-mediated voyage, or a mislabel in the collection.

The 22 newly generated ancient “Botocudo” and the Sambaqui and mummy genomes indicate that these people were all Indigenous American harboring genetic ancestry common in present-day Indigenous populations in Brazil. Furthermore, the “Botocudo” exhibit one of the lowest heterozygosity levels and some of the longest runs of homozygosity among worldwide populations sequenced to date. This could suggest, as has been hypothesized (Barbieri et al. 2018; Ceballos et al. 2018; Neel et al. 1967; Pemberton et al. 2012), that Native American hunter-gatherers generally had different social and marital practices than Eurasian hunter-gatherers (Sikora et al. 2017).

Among Natives from Brazil, “Botocudos” were found to be distantly related to Tupi-speaking populations, whose ancestors expanded to most of South American lowlands during the Late Holocene. The individuals sequenced in this study were inferred to be more closely related to the Xavante, a present-day Jê-speaking population, suggesting that the “Botocudos” may also have spoken a Jê language as recorded in the 18^th^ and 19^th^ century literature (Ehrenreich 2014; Paraíso 1992). Yet, in our analyses, the Xavante were not significantly closer to the “Botocudos” than other Indigenous populations in Brazil and the genetic drift that the Xavante and “Botocudos” experienced after their split was high, suggesting that these groups have substantially differentiated from each other.

All in all, the “Botocudos” are Indigenous Americans, and the genetically closest populations (sequenced to date) are found in South America, on the east side of the Andes. All these Indigenous populations presented a similar relationship to the “Botocudo” individuals analyzed here, but a strong affinity to any particular population could not be established. Thus, although we observe a shared genomic history between these “Botocudos” and South Native Americans, the lack of a strong affinity to a specific population indicates that we are recovering some of the genetic diversity that has not been explored so far in South America. It would be of particular interest to establish the genetic relationship between surviving “Botocudo” communities, like the Krenak and Aranã people in Minas Gerais or other present-day populations self-identifying as “Botocudos”, and the ancient “Botocudos” analyzed here. Genetic studies of additional present-day Indigenous populations should allow to resolve this question in the future.

### Ethics Statement

This project started off as a collaboration following the publication of Malaspinas et al., (2014) between Brazilian researchers at the National Museum in Rio de Janeiro and researchers who are now based at the University of Lausanne. To facilitate the collaboration and to favor knowledge exchange, several visits at the National Museum in Rio de Janeiro took place. During those visits, preliminary data were presented to the staff and students working at the National Museum. These visits and the regular exchanges with local researchers helped to better define the research questions to be addressed within the study, to present this work to broader audiences, and to prepare a report that will be presented in lay language to the Indigenous communities.

Following Brazilian law, prior to any sampling, a project proposal was submitted for authorization from the Instituto do Patrimônio Histórico e Artístico Nacional (IPHAN) by National Museum in Rio de Janeiro and the IPHAN delivered the research permits for this work (IPHAN numbers 01500.001759/2016-15 and 01500.001824/2018-66). Note that the remaining bone will be returned to the Museum at the end of the project (01500.001824/2018-66). Human research in Switzerland is only permissible upon review and approval by the responsible ethics committee (Human Research Act, HRA). Hence, the project proposal was also submitted (Req-2018-00993) to the Commission Cantonale d’Ethique de la Recherche sur l’être humain (CER-VD), responsible ethics committee for the research projects taking place in the Canton de Vaud, Switzerland. The CER-VD ruled that the project was not falling within the scope of the HRA and thus did not require an authorization for the analyses to be carried out in Switzerland. Microbial species discovered in this study were registered at the Sistema Nacional De Gestão Do Patrimônio Genético E Do Conhecimento Tradicional Associado (registration number: ADC5681).

While conducting research following the local laws is obviously a necessity, community engagement should become an integrant part of ancient DNA research (Wagner et al. 2020). However, many Brazilian Indigenous communities (including those who donated samples, Castro e Silva et al. 2020; Skoglund et al. 2015) could not be contacted by non-Indigenous people during the pandemic. To reach out, we have summarized our study in a report in Portuguese that will be included in a compilation of results that will be presented to Indigenous populations (Skoglund et al. 2015; Castro e Silva et al. 2020) who have asked to remain informed about the scientific work that involves their data.

## Author Contributions

Conceived the project: ASM. Designed the study: ASM, DICD with input from SR, MQRB, CRC. Supervised the study: ASM, MCAA. Provided archival and contextual information: SR, MQRB, CRC. Performed the laboratory work: JS, FEY, VVI under the supervision of MEA and MCAA. Performed the isotope analysis: MQRB and DICD. Analyzed the genetic data: DICD, YOAC, MJBL, JVMM, SN, CEGA under the supervision of ASM and MCAA. Wrote the supplementary information: DICD, YOAC, MJBL, MCAA, ASM with critical input from all coauthors. Wrote the main text: DICD, YOAC, MJBL, SR, MQRB, JVMM, CEGA, MCAA, ASM with critical input from SN, JS, FEY, VVI, CSRB, CRC, TH and MEA.

## Acknowledgments

We thank Peter de Barros Damgaard, Federico Sánchez-Quinto, Barbara Mühlemann, Nicolás Rascován, Alexandre Reymond, Laurent Excoffier and Ludovic Orlando for helpful discussions. We thank Alejandra Castillo and Carina Díaz for technical support at LIIGH, and to Luis Alberto Aguilar Bautista, Alejandro de León Cuevas, Carlos Sair Flores Bautista and Jair Garcia Sotelo from the “Laboratorio Nacional de Visualización Científica Avanzada” (LAVIS-UNAM) for providing computational resources and support. We would also like to thank Vital-IT and UNIL-DCSR for computer infrastructure and support and the GeoGenetics Sequencing Core at the University of Copenhagen for sequencing and basecalling. ASM, DICD, YOAC, SN and CEGA were funded by an SNSF and an ERC grant to ASM. JS and FEY were funded by a Swedish Foundation for Humanities and Social Sciences grant (M16-0455:1, The Rise III) to Kristian Kristiansen. MJBL was funded by project IA201219 PAPIIT-DGAPA-UNAM. MCAA was funded by the Wellcome Trust (WT_208934/Z/17/Z).

## Materials and Methods

### DNA extraction, library preparation, and sequencing

#### Shotgun data

Laboratory work was performed in dedicated clean laboratory facilities at the Centre for GeoGenetics, Natural History Museum, University of Copenhagen. Twenty-four DNA extracts were prepared: one from petrous bone, one from a piece of skull and twenty-two from teeth. Between 100 and 200 mg of sample material were pulverized, pre-digested and digested using established protocols for ancient DNA (Damgaard et al. 2015). The DNA was then purified using the method described by (Allentoft et al. 2015). The extract for one library of the individual MN0008 was treated with 5 μl of the USER^TM^ enzyme for 3 hours at 37°C, and end-repaired using NEBnext end-repair module. Between one and three double-stranded DNA libraries were built per individual sample, using Illumina-specific adaptors, and following (Kircher et al. 2012) for double indexing. The libraries were amplified, and the number of PCR cycles was determined by qPCR. The indexed and amplified libraries were purified and quantified on an Agilent 2200 TapeStation before being pooled in equimolar amounts. The libraries were then sequenced (100 bp, single-end, Dataset S2) on Illumina HiSeq2500 and Hiseq4000 platforms and basecalled using CASAVA (version 1.8.2) at the GeoGenetics Sequencing Core at the University of Copenhagen.

#### Mitochondrial DNA capture

Capture-enrichment of human mitochondrial genome was performed using the single indexed libraries of the samples MN01701 (Sambaqui) and MN1943 (mummy) with the commercial kit myBaits Mito from Arbor biosciences. Capture was done following the manufacturer’s protocol v.4, setting the hybridization time to 42 hours. After capture, we performed a qPCR analysis using the Maxima SYBR Green/rROX qPCR Master Mix to determine the number of cycles to reamplify the captured libraries. Then, both libraries were reamplified for 11 cycles using the enzyme Phusion U Hot Start DNA Polymerase from Thermo Fisher Scientific. Purification of libraries was made with the Magnetic beads SPRISelect from Beckman Coulter. To determine the concentration and average length of the DNA, libraries were analyzed in the Bioanalyzer 2100 from Agilent. The libraries were then sequenced (75 bp, paired-end, Dataset S2) on Illumina NextSeq 500 platforms at the INMEGEN (National Institute of Genomic Medicine, Mexico City) genomics facility.

### ^14^C dating and stable isotopes measurements

Twenty-two samples (20 “Botocudos”, the mummy and the Sambaqui, Dataset S18) were dated at the 14CHRONO Centre radiocarbon dating facility from the Queen’s University, Belfast. Pretreatment, combustion and graphitization, AMS dating, and measurement of C:N ratios, δ^13^C and δ^15^N were performed following (Reimer et al. 2015).

As the isotope data suggested a potential high consumption of marine fauna for the Sambaqui-MN01701 (Fig. S15), we inferred the protein marine diet proportion for this individual using Bayesian mixing models in Fruits 3.0 (Fernandes et al. 2014), with δ^13^C and δ^15^N values of three different food sources as published in (Pezo-Lanfranco et al. 2018). The food sources consisted of marine and terrestrial fauna (archaeological terrestrial herbivores from the southern coast of Brazil (Colonese et al. 2014), and modern C3 plants from the southeastern Atlantic Forest in Brazil (Galetti et al. 2016).

We then calibrated the dates with OxCal 4.4 (Bronk Ramsey 2009) for the 20 “Botocudos”, the mummy-MN1943, the Sambaqui-MN01701 and the two Polynesian-“Botocudos” (Bot15 and Bot17, Malaspinas et al., 2014) (Fig. 1). For the individuals with a high marine protein consumption (Sambaqui-MN01701 and Bot17, cite), the dates were calibrated using mixed curves consisting of a Southern Hemisphere (Hogg et al. 2020) and a Marine (Heaton et al. 2020) calibration curves. Protein marine proportions of 0.70±0.09 (as described above for Sambaqui-MN01701) and 0.60±0.16 (Bot17, as in Malaspinas et al., 2014) were specified for the Marine curves. A local R offset of ΔR = 111±43 years (calib.org/marine) was used for Sambaqui-MN01701, whereas a ΔR = 0±20 years was used for the Polynesian-“Botocudo” Bot17 as proposed in (Malaspinas et al., 2014). Finally, for the 20 “Botocudos”, the mummy-MN1943 and the Polynesian-“Botocudo” Bot15, the radiocarbon dates were calibrated using a Southern Hemisphere calibration curve (Hogg et al. 2020).

### Mapping and variant calling

#### Mapping the genomic data to the human genome

Illumina adapter sequences, low-quality bases and nucleotides reported as “N” were trimmed from the reads with AdapterRemoval version 2.1.7 (Schubert et al. 2016). Reads of 30 bp length and above were mapped to the human reference genome *built* 19 with BWA aln version 0.7.15 (Li et al., 2009), disabling the seed to avoid mapping bias due to ancient DNA damage at the 5’ termini of the reads (Schubert et al. 2012). Reads with a mapping quality score equal or greater than 25 (for the two mitochondrial capture libraries only) or 30 (for the other 38 libraries) were retained. Duplicate reads were identified and removed with Picard tools MarkDuplicates version 2.9.0 (http://broadinstitute.github.io/picard/), and indel realignment was performed with GATK version 3.7 with default options (McKenna et al. 2010). MD tags were then recomputed with SAMtools version 1.8 (Danecek et al. 2021). Molecular damage parameters were obtained with mapDamage2 (Jónsson et al. 2013). Statistics associated to trimmed and mapped reads were computed using BEDTools (Quinlan et al. 2010) and customized python (version 3.6) scripts. Mapping statistics are reported in Table 1 and Dataset S2.

#### Genotype likelihoods

Genotype likelihoods were computed with ANGSD (Korneliussen et al. 2014), selecting the SAMtools model (-GL 1), and using bases with a minimum PHRED-score of 20 (-minQ 20) unless otherwise specified.

#### Genotype calling

Genotypes were called with SAMtools mpileup and BCFtools call commands (Danecek et al. 2021) as follows. The mapping qualities of the reads were adjusted (-C 50, value recommended by the SAMtools authors for reads mapped with BWA), and the reads with a minimum mapping quality of 30 (-q 30) after the readjustment were used for the mpileup step. Genotypes were then called using the original consensus caller (-c) implemented in BCFtools/SAMtools. These “raw” genotype calls were then filtered in four main steps with BCFtools (Danecek et al. 2021), retaining only genotype calls that meet the following four conditions: the genotype overlaps with the 1000 Genomes strict mask (Altshuler et al. 2012); it is located outside of the RepeatMask regions (available at UCSC Table Browser, http://genome.ucsc.edu/, Karolchik et al. 2004); its depth of coverage is above one-third and below twice the average depth of that of the individual genome analyzed; its genotype quality score is equal or greater than 30.

### Error rate estimation

Error rates were estimated for the 24 individual samples (Figs. S3-S5) by comparing the sequenced genomes to an outgroup and an error-free individual (an approach introduced in Orlando et al. 2013). To estimate error rates, the mapped reads were compared to the consensus sequence of an outgroup (chimpanzee), and to that of a high-coverage present-day human genome (Dinka individual; ID: SS6004480; depth of coverage: 37×; Prüfer et al. 2014). The consensus sequence for the Dinka individual was inferred using ANGSD (-doFasta 2) (Korneliussen et al. 2014) from reads with a minimum mapping quality of 30 (-minMapQ 30) and bases with a minimum base quality of 30 (-minQ 30). The error rates were then estimated using all the reads (-doAncError 1) with ANGSD (Korneliussen et al. 2014) and bases with a minimum quality score of 30 (-minQ 30).

### Molecular sex determination and uniparental markers

Molecular sex was determined by computing the ratio of reads mapping to the Y chromosome with respect to those mapping to the sex chromosomes, and following the threshold established in (Skoglund et al. 2013) to identify females or males. To call Y-chromosome and mitochondrial haplogroups, we first used ANGSD version 0.921 (Korneliussen et al. 2014) to generate the consensus sequence by taking the most common base (-doFasta 2 -doCounts 1 -minQ 20) across the Y or mitochondrial chromosome. Y-chromosome haplogroups were called by querying the consensus sequence against the Y-chromosome variants reported in the phase 3 of the 1000 Genomes Project following (Poznik 2016). Mitochondrial haplogroups were obtained by uploading the consensus sequence in FASTA format to James Lick’s website (https://dna.jameslick.com/mthap/). Molecular sex and uniparental haplogroups are presented in Table 1 and Dataset S2.

### Contamination estimation

Contamination was estimated with (i) contamMix (Fu et al. 2013) for the mtDNA for mitochondrial data with a DoC above 10×–as this method assumes the consensus is the truth–and (ii) with contaminationX (Moreno-Mayar et al. 2020) for X chromosome data in males with an average depth of coverage above 0.5× on that chromosome (as suggested in Moreno-Mayar et al. 2020). Both estimates are reported in Table 1 and Dataset S2.

#### Estimation with contamMix

The mitochondrial consensus sequence was generated per individual with ANGSD (-doFasta 2 - doCounts 1) (Korneliussen et al. 2014). The mitochondrial reads were then remapped with BWA aln version 0.7.15 (Li et al., 2009) to the individual’s consensus sequence, realigned with GATK v3.7 (McKenna et al. 2010), and their MD tags were recomputed with SAMtools version 1.8 (Danecek et al. 2021) as described above. The consensus sequence of each individual was added to a database of 311 worldwide mitochondrial sequences (Green et al. 2008) and aligned with MAFFT (Katoh et al. 2013). contamMix (Fu et al. 2013) was run per individual, with the reads aligned to the consensus sequence and the multiple alignment of 312 mitochondrial sequences as input. The Monte Carlo Markov chain (MCMC) was run for 3 chains, with 100,000 samples each and discarding the first 10,000 iterations. Note that the potential-scale reduction factor values were below 1.05, indicating that the variance over chains versus pooled variance was less than 5% (Gelman-Rubin estimator R, 1<= R <= 1.05) for all runs which suggests that the MCMC chains may have converged.

#### Estimation with contaminationX

Following (Moreno-Mayar et al. 2020) allele counts were generated on the X chromosome with ANGSD (Korneliussen et al. 2014) for bases with a minimum quality score of 20 (-doCounts 1 -iCounts 1 -minQ 20). The counts were then filtered with contaminationX, keeping those with a minimum depth of 3 (-d 3), excluding pseudoautosomal regions (-b 5000000 -c 154900000), using the HapMap CEU allele frequencies (Altshuler et al. 2010) for estimation, and discarding variants with allele frequencies below 0.05 (-m 0.05). The estimates were then obtained allowing for up to 1000 blocks for the jackknife procedure (-maxsites=1000).

### Reference panels and dataset merging

Publicly available data were assembled into several datasets for the population genetic analyses (Dataset S19). Some datasets mostly contained data derived from SNP arrays, whereas others are composed of genomes (sequencing reads) from different individuals. Depending on the analysis performed with panels including SNP arrays, the data of the individuals sequenced in this study were merged either by: sampling a random read (MDS, *f_3_*); computing genotype likelihoods (NGSadmix and fastNGSadmix); calling genotypes (qpGraph). For the analyses with whole genome datasets, the external data and the genomes sequenced in this study were processed in the same way: sampling random reads (conditional heterozygosity analysis); calling genotypes (ROH and PSMC); or using all reads as input (error-corrected D-statistics). Genotype likelihoods and genotype calling are described under the “Mapping and variant calling” section above. Below we list and describe the datasets that were compiled for different analyses presented in the main text.

#### Populations names

Three populations mentioned in the main text bear exonyms as population names in the publications in which their genomic data was first published. We replaced those names by their endonyms in the main text and figures: Paiter (previous label: Surui), Yjxa (previous label: Karitiana) and Wixárika (previous label: Huichol).

#### Datasets including SNP array data

Genotype calls from different SNP panels were merged with PLINK (Purcell et al. 2007). The panels were merged on matching autosomal positions, and variants with more than two alleles were removed.

1. MDS_NGSadmix_Wollstein_Xing_Malaspinas. This panel includes 823,805 SNPs (567,544 transitions and 256,261 transversions), and 585 individuals from 20 worldwide populations. The data of 583 modern individuals (from Wollstein et al. 2010; Xing et al. 2010) was assembled in (Malaspinas et al., 2014). The two ancient Polynesian “Botocudos” (Malaspinas et al., 2014) were merged to this panel by sampling a random read. For the MDS analyses, the individuals sequenced in this study were merged by sampling a random read at each of the 823K positions. For NGSadmix analyses, genotype likelihoods were computed for the individuals sequenced in this study and merged to the panel. MDS and NGSadmix analyses were run with all the SNPs (Datasets S5 and S8) or only the transversions of this panel (Fig. 2A and Datasets S6 and S7).
2. NGSadmix_fastNGSadmix_Wollstein_Xing. This panel includes 256,261 SNPs (transversions only), and 144 individuals representing six worldwide present-day populations. A total of 24 individuals were sampled randomly from the following populations: Yoruba (Africa), CEU (Utah residents with Northern and Western European ancestry), Japanese (East Asia), New Guineans (Oceania), Tongans or Samoans (Polynesia), and Totonacs (Americas). The panels from which the individuals were sampled were originally published in (Wollstein et al. 2010; Xing et al. 2010) and assembled in (Malaspinas et al., 2014). Ancestry proportions were inferred for the 144 individuals with NGSadmix (K = 6, i.e. the number of populations included in the analysis); the proportions and inferred allele frequencies per component were then used as input to infer ancestry proportions with fastNGSadmix for the 24 individuals sequenced in this study as well for the two Polynesian “Botocudos” (Malaspinas et al., 2014) (Figs. 2B and S6 and Dataset S7).
3. F3_Raghavan_Skoglund_Castro_AncAmericas. This panel includes 56,646 SNPs (56,134 transitions and 512 transversions) and 775 individuals from 167 Native American populations. The data of 647 Native Americans from different publications were assembled and masked for non-Indigenous American ancestry in (Raghavan et al., 2015). The data of 12 present-day Native Brazilians from (Castro e Silva et al. 2020), as well as 43 present-day Native Brazilians from (Skoglund et al. 2015) were then merged, resulting in 702 individuals and 56,646 SNPs. We then added the data of 73 ancient Indigenous Americans (panel F3_73AncNatAm described under “Whole genome datasets” and Dataset S19) by sampling a random read at each of the 56K positions from trimmed (5 bp on each end) reads. Similarly, 22 “Botocudos” were merged by sampling an allele after trimming 5 bp on each end on the reads (Fig. 3).
4. NGSadmix_Raghavan_Skoglund_Castro. This panel includes 56,646 SNPs (56,134 transitions and 512 transversions) and 333 modern individuals from 38 populations. To perform ancestry estimations, four populations (ten individuals each) out of the Americas (Yoruba, Papuans, French and Dai) and 20 populations (238 individuals) from the Americas were extracted from the panel compiled in (Raghavan et al., 2015). We then merged the data of 14 populations (55 individuals): five populations (12 individuals published by Castro e Silva et al. 2020) and nine populations (43 individuals) from Brazil published by (Skoglund et al. 2015). The genotype calls of the 333 present-day individuals were recoded as genotype likelihoods and merged to the genotype likelihoods computed after trimming 5 bp for the 9 “Botocudos” with a DoC above 0.1× (Fig. 4) and the 22 “Botocudos”, the mummy and the Sambaqui individual from this study (Dataset S10).
5. F3_Raghavan. This panel includes 199,285 SNPs (197,101 transitions and 2,184 transversions) and 647 individuals from 80 Native American populations, assembled and masked for non-Indigenous American ancestry in (Raghavan et al., 2015). To compute outgroup-*f_3_* statistics (Fig. S7), 22 “Botocudos” were merged to this panel by sampling an allele after trimming 5 bp on each end on the reads.
6. F3_NGSadmix_Raghavan_CastroTupGuaUnrel. This panel includes 40,146 SNPs (39,778 transitions and 368 transversions) and 290 present-day individuals from 22 Native American populations. To perform ancestry estimations (Dataset S11), four populations (ten individuals each) out of the Americas (Yoruba, Papuans, French and Dai) and 20 populations (238 individuals) from the Americas were extracted from the panel compiled by (Raghavan et al., 2015). We then merged the data of 2 populations (52 individuals) from Brazil published by (Castro e Silva et al. 2020). The genotype calls of the 330 present-day individuals were recoded as genotype likelihoods and merged to the genotype likelihoods computed after trimming 5 bp for the 22 “Botocudos”, the mummy and the Sambaqui individual from this study. To compute outgroup-*f_3_* statistics (Fig. S8), 22 “Botocudos” were merged to this panel by sampling an allele after trimming 5 bp on each end on the reads.
7. F3_NGSadmix_Raghavan_CastroTupGuaUnrel_Castro12NatAm_Skoglund. This panel includes 13,490 SNPs (13,397 transitions and 93 transversions) and 345 individuals from 43 Native American populations. To perform ancestry estimations, four populations (ten individuals each) out of the Americas (Yoruba, Papuans, French and Dai) and 20 populations (238 individuals) from the Americas were extracted from the panel compiled by (Raghavan et al., 2015). We then merged the data of 14 populations (107 individuals) from Brazil: five populations (64 individuals published by Castro e Silva et al. 2020) and nine populations (43 individuals) published by (Skoglund et al. 2015). The genotype calls of the 345 present-day individuals were recoded as genotype likelihoods and merged to the genotype likelihoods computed after trimming 5 bp on the ends of the reads for the 22 “Botocudos”, the mummy and the Sambaqui individual from this study (Dataset S12). To compute outgroup-*f_3_* statistics (Fig. S9), 22 “Botocudos” were merged to this panel by sampling an allele after trimming 5 bp on each end on the reads.
8. F3_NGSadmix_Skoglund_Castro12NatAm. This panel includes 606,888 SNPs (430,944 transitions and 175,944 transversions), and 95 individuals from 16 populations. To perform ancestry estimations, 40 individuals from two populations out of the Americas (Yoruba and French, 20 individuals each) were randomly selected from the Allen Ancient DNA Resource (https://reich.hms.harvard.edu/allen-ancient-dna-resource-aadr-downloadable-genotypes-present-day-and-ancient-dna-data, version 44.3; genomes originally published in Bergström et al. 2020; Mallick et al. 2016; Meyer et al. 2012; Prüfer et al. 2014). We then merged the data of 14 populations (55 individuals) from Brazil: five populations (12 individuals) published by (Castro e Silva et al. 2020) and nine populations (43 individuals) published by (Skoglund et al. 2015). The genotype calls of the 95 present-day individuals were recoded as genotype likelihoods and merged to the genotype likelihoods computed after trimming 5 bp for the 22 “Botocudos”, the mummy and the Sambaqui individual from this study (Dataset S13). To compute outgroup-*f_3_* statistics (Fig. S10), 22 “Botocudos” were merged to this panel by sampling an allele after trimming 5 bp on each end on the reads.
9. F3_NGSadmix_Skoglund_CastroTupGuaUnrel_Castro12NatAm. This panel includes 77,766 SNPs (63,690 transitions and 14,077 transversions) and 147 individuals. To perform ancestry estimations, forty individuals from two populations out of the Americas (Yoruba and French, twenty individuals each) were randomly selected from the Allen Ancient DNA Resource (https://reich.hms.harvard.edu/allen-ancient-dna-resource-aadr-downloadable-genotypes-present-day-and-ancient-dna-data, version 44.3; genomes originally published in Bergström et al. 2020; Mallick et al. 2016; Meyer et al. 2012; Prüfer et al. 2014). We then merged the data of 14 populations (107 individuals) from Brazil: five populations (64 individuals) published by (Castro e Silva et al. 2020) and nine populations (43 individuals) published by (Skoglund et al. 2015). The genotype calls of the 147 present-day individuals were recoded as genotype likelihoods and merged to the genotype likelihoods computed after trimming 5 bp for the 22 “Botocudos”, the mummy and the Sambaqui individual from this study (Dataset S14). To compute outgroup-*f_3_* statistics (Fig. S11), 22 “Botocudos” were merged to this panel by sampling an allele after trimming 5 bp on each end on the reads.
10. qpGraph_Skoglund_Castro. This panel includes 606,888 SNPs (493,453 transitions and 113,435 transversions), and 95 individuals from 16 populations. The following populations were selected to build admixture graphs with qpGraph: Yoruba (n = 20, Bergström et al. 2020; Mallick et al. 2016; Meyer et al. 2012; Prüfer et al. 2014; Allen Ancient DNA Resource https://reich.hms.harvard.edu/allen-ancient-dna-resource-aadr-downloadable-genotypes-present-day-and-ancient-dna-data, version 44.3), Xavante (n = 11, Skoglund et al. 2015), Yjxa (n = 4, Skoglund et al. 2015), Paiter (n = 4, Skoglund et al. 2015), Guarani Kaiowá (n = 9, Skoglund et al. 2015), and Urubu Ka’apor (n = 3, Skoglund et al. 2015). Genotypes were called for the “Botocudo” MN0008 (USER-treated library only, 2 bp trimmed on each end of the reads) and merged to the panel. Admixture graphs with this panel are shown in Figs. 5 and S12.

#### Whole genome datasets

For the panels ROH, PSMC, and ErrorCorr_Dstat, we integrated the data of the “Botocudo” MN0008, corresponding to the USER-treated extract and with 2 bp trimmed at the end of the reads.

1. Conditional_heterozygosity. This panel includes twenty populations, two individuals per population (Fig. 6A). The 20 populations are distributed as follows: four from Africa (Meyer et al. 2012; Prüfer et al. 2014), two from Europe (Meyer et al. 2012; Prüfer et al. 2014), two from East Asia (Meyer et al. 2012; Prüfer et al. 2014), six from Oceania (Meyer et al. 2012; Prüfer et al. 2014; Malaspinas et al., 2014), five present-day populations from the Americas (Mallick et al. 2016; Meyer et al. 2012; Prüfer et al. 2014; Skoglund et al. 2015), three ancient populations from the Americas (Moreno-Mayar, Vinner, et al. 2018; Moreno-Mayar, Potter, et al. 2018), and the ancient “Botocudos” (this study). A read was sampled per position per individual from the mapped reads as described in the “Conditional heterozygosity” section.
2. ROH. This panel includes fifteen individuals from nine populations (Fig. 6B): “Botocudos” (n = 1, this study), Spirit Cave (n = 1, Moreno-Mayar, Vinner, et al. 2018), Mixe (n = 1, Meyer et al. 2012), Paiter (n = 2, Skoglund et al. 2015), Yjxa (n = 2, Meyer et al. 2012; Prüfer et al. 2014), Papuan (n = 2, Meyer et al. 2012; Prüfer et al. 2014), Dai (n = 2, Meyer et al. 2012; Prüfer et al. 2014), French (n = 2, Meyer et al. 2012; Prüfer et al. 2014), and Yoruba (n = 2, Meyer et al. 2012; Prüfer et al. 2014). Genotypes were called for each individual as described in the “Genotype calling” section. ROH per genome were inferred as described in the “Runs of homozygosity” section.
3. PSMC. This panel includes six individuals (Fig. 6C): MN0008 (“Botocudo”, this study); LP6005441-DNA_A12 (Paiter, Skoglund et al. 2015); HGDP00998 (Yjxa, Meyer et al. 2012); LP6005441-DNA_G07 (Maya, Mallick et al. 2016); HGDP01307 (Dai, Meyer et al. 2012); HGDP00927 (Yoruba, Meyer et al. 2012). Genotypes were called for each individual as described in the “Genotype calling” section.
4. ErrorCorr_Dstat. This panel includes 26 individuals (Figs. 7 and S14). The consensus sequence of one Yoruba individual (Meyer et al. 2012) was used as a proxy for the ancestral allele in the outgroup population. Three Mixe individuals (Prüfer et al. 2014; Skoglund et al. 2015) represent the population in H_2_. For the populations in H_3_ for which gene flow was tested, French (n = 1, Meyer et al. 2012) and Han (n = 1, Meyer et al. 2012) genomes represented the Eurasian populations, whereas Australians (n = 7, Malaspinas et al., 2016), a Papuan (n = 1, Meyer et al. 2012), and a historical Andaman (n = 1, Moreno-Mayar, Vinner, et al. 2018) were chosen as representatives of Australasia. For the Indigenous Americans in H_1_, we included the genomes of the following populations: “Botocudo” MN0008_L3U_trim2 (n = 1, this study); the 10,000 yo Lagoa Santa (n = 1, Moreno-Mayar, Vinner, et al. 2018); Paiter (n = 2, Skoglund et al. 2015); Yjxa (n = 4, Mallick et al. 2016; Prüfer et al. 2014; Rasmussen et al. 2014); Aymara (n = 1, Raghavan et al., 2015); and Wixárika (n = 1, Raghavan et al., 2015). For the ancient Lagoa Santa and the historical Andaman individuals, 5 bp were trimmed at each end of the reads.
5. F3_73AncNatAm. This panel includes 73 ancient individuals from 30 populations in the Americas (Dataset S19 and Fig. 3). The populations span North and South America (Moreno-Mayar, Vinner, et al. 2018; Moreno-Mayar, Potter, et al. 2018; Posth et al. 2018; Raghavan et al., 2014; Raghavan et al., 2015; Rasmussen et al. 2014; Scheib et al. 2018; Schroeder et al. 2018). For each ancient genome, the reads were trimmed (5 bp on each end), and an allele was sampled and merged to the panel F3_Raghavan_Skoglund_Castro_AncAmericas (56,646 SNPs).

### Multidimensional scaling

For the multidimensional scaling (MDS) analyses with worldwide populations the panel MDS_NGSadmix_Wollstein_Xing_Malaspinas was used (described in “Panels and data sets merging” and Dataset S19). Allele sharing distances were computed with PLINK (Purcell et al. 2007) on pseudo-haploid data of the panel (i.e., a random allele per individual per site) and either (i) the 24 sequenced individuals or (ii) each of the sequenced individuals in this study; for both cases the distances were computed using (a) only transversions and (b) all the SNPs. Two individuals (“Botocudo” MN00010 and Sambaqui MN01701) did not have data overlapping with other individuals in (ia), so they were excluded from the projection of (ia). Classic MDS projections were performed in R (Core Team et al. 2020) with the function cmdscale() with the pairwise distances as input. See Fig. 2A and Datasets S5 and S6.

### Clustering analyses - Ancestry estimation

Panels were compiled to assess with clustering analyses whether there were further Polynesians (MDS_NGSadmix_Wollstein_Xing_Malaspinas and NGSadmix_fastNGSadmix_Wollstein_Xing), and to explore the genetic structure within South America (NGSadmix_Raghavan_Skoglund_Castro). To estimate ancestry proportions, we used NGSadmix (Skotte et al. 2013) and fastNGSadmix (Jørsboe et al. 2017). The first approach allows us to estimate ancestry in an unsupervised fashion on the joint data with the classic STRUCTURE model (Pritchard et al. 2000), whereas the second allows us to estimate the ancestry in a single individual using pre-inferred ancestry proportions and allele frequencies in the reference panel. Both methods can take genotype likelihoods as input. Therefore, we converted the SNP calls in the panels to genotype likelihoods, and computed genotype likelihoods for the twenty-four Native Brazilians as described above. For the panels, the genotype likelihoods matrix was prepared by setting a value of one on the entry corresponding to the genotype observed, and zero for the other two alternative genotypes (i.e., the entries for a heterozygous call would be coded as ’0 1 0’). NGSadmix (Skotte et al. 2013) was run with default parameters (i.e., on sites with a minor allele frequency of 0.05,-minMaf 0.05).

#### Panels with Polynesians: NGSadmix_fastNGSadmix_Wollstein_Xing

We first modelled population structure for the six reference populations (Yoruba, CEU, Japanese, New Guineans, Tongans and Samoans, and Totonacs) in the panel NGSadmix_fastNGSadmix_Wollstein_Xing (Dataset S19) considering K = 6 ancestry groups using NGSadmix (Skotte et al. 2013). The ancestry estimates and allele frequencies of the NGSadmix run with the highest likelihood among 100 replicates was chosen as input for fastNGSadmix (Jørsboe et al. 2017) to estimate their proportions in the two Polynesian-“Botocudos”, the 22 “Botocudos”, the mummy and the Sambaqui. Figs. 2B and S6 show the fastNGSadmix results for K = 6 (see Datasets S5 and S6 for NGSadmix analyses). Genotype likelihoods were computed as described above for the 22 “Botocudos”, the mummy, the Sambaqui individual, and the two Polynesian “Botocudos” (Malaspinas et al., 2014). fastNGSadmix (Jørsboe et al. 2017) was run specifying 100 bootstraps, and K = 6 ancestral components (Fig. 2B). See Fig. S6 for the analyses including transitions.

#### Panels with Polynesians: MDS_NGSadmix_Wollstein_Xing_Malaspinas

NGSadmix was also run separately for each of the 24 sequenced individuals with the panel MDS_NGSadmix_Wollstein_Xing_Malaspinas. The data of the 583 individuals of the panel was merged to the genotype likelihoods of each of the 24 Native Brazilians, and sites with missing data in the ancient Brazilian being analyzed were removed. Ten replicates were run with NGSadmix assuming K = 6. The replicate with the highest likelihood was then plotted (see Datasets S8 and S9).

#### Panel enriched in Native American populations: NGSadmix_Raghavan_Skoglund_Castro

To explore the genetic structure in South America, we estimated ancestry proportions using the panel NGSadmix_Raghavan_Skoglund_Castro. As most of the SNPs in this panel are transitions, to reduce the effect of ancient DNA damage, 5 bp were trimmed on each end of the reads of the 22 “Botocudos”, the mummy and the Sambaqui individual prior to genotype likelihoods computation, and only bases with a minimum quality score of 30 were considered to calculate the likelihoods. The genotype likelihoods of the 24 individuals were then merged to the panel, and ten replicates of NGSadmix were run for each value of K from 2 to 15. The run with the highest likelihood for K = 13 and including the nine “Botocudos” with a DoC above 0.1× was plotted in Fig. 4; for other runs including the 22 “Botocudos”, the mummy and the Sambaqui individual, see Dataset S10. For runs with other panels enriched in Native Americans (described in Dataset S19), see Datasets S11 – S14.

### Outgroup-*f*_3_

Outgroup-*f_3_* statistics were computed with ADMIXTOOLS version 5.1 (Patterson et al. 2012). To investigate the genetic affinity between the “Botocudos” and the Native American populations, outgroup-*f_3_* statistics of the form *f_3_*(Yoruba; Botocudo, Native American) were calculated using the panel F3_Raghavan_Skoglund_Castro_AncAmericas described in Dataset S19, which includes 22 Botocudos. We report results for comparisons for which there were at least 5,000 SNPs out of the 56,646 included in the panel. Tests with further panels enriched in Native American populations (Dataset S19) are shown in Figs. S7 – S11.

### Admixture graphs

We used qpGraph (version 6100) to fit *f*-statistics-based admixture graphs (Patterson et al. 2012) for the populations in the panel qpGraph_Skoglund_Castro (Dataset S19). We followed an admixture graph search strategy similar to that in (Moreno-Mayar, Vinner, et al. 2018), where we start with a simple graph modelling the relationship between a few key populations. We then extended the starting graph by including each of the additional populations as either a non-admixed or an admixed leaf in every possible position of the graph. For each additional population, we assessed the fit of each topology using three criteria: 1) its fit score, 2) the Z-score of the worst residual between the predicted and the observed *f*4-statistics, and 3) the presence of trifurcations. We carried out comparisons between non-admixed and admixed models using a likelihood ratio test following (Lipson et al. 2017). Here, fit score differences of ∼3 and ∼4.6 correspond to p-values of 0.05 and 0.01 respectively.

### Conditional heterozygosity estimation

To estimate heterozygosity in the “Botocudo” population, we followed (Skoglund et al. 2014) as the data sequenced here include damaged low-depth genomes. First, we ascertained heterozygous sites from the autosomes of an African individual (Yoruba from Nigeria, SS6004475, Mallick et al. 2016) relying on the genotype calls in the Simons Genome Diversity Project (SGDP) panel. Transitions were filtered out, as well as sites that had more than one alternative allele in any of the individuals from the SGDP panel (Mallick et al. 2016). We then selected two Botocudo genomes and sampled a random read from each genome at every ascertained position in the Yoruba individual. Similarly, we analyzed two individuals from other 19 populations (panel Conditional_heterozygosity, Dataset S19), and drew one allele per individual at a given position. The conditional heterozygosity per population was calculated as the proportion of observed heterozygous sites to the total sites with data. Confidence intervals were computed by applying a jackknife resampling strategy over blocks of 5 Mb along the genome. Results for the 19 populations and the pair of “Botocudos” MN0008 (DoC: 23.9×) and MN00056 (DoC: 3.3×) are shown in Fig. 6A. Estimates for all pairs of “Botocudos” are shown in Fig. S13.

### Runs of homozygosity

For fifteen individuals (panel ”ROH” in Dataset S19), genotypes were called as described in the “Genotype calling” section. Runs of homozygosity were inferred with PLINK (--homozyg) (Purcell et al. 2007) on the genotype calls using default parameters (--homozyg-snp 100--homozyg-density 50--homozyg-gap 1000--homozyg-window-het 1--homozyg-window-het 5) and by selecting runs of at least 500 Kb long (--homozyg-kb 500), following (Gamba et al. 2014).

### Population size estimation

Effective population sizes were estimated with PSMC (Heng Li et al. 2011) for six individuals (dataset “PSMC”, Dataset S19). For each of the six individuals, genotypes were called as described in the “Genotype calling” section and used as input for PSMC. PSMC was run with parameters suggested by the developers for humans: a maximum of 25 iterations (-N 25), a maximum coalescence time of 2N0 = 15 (-t 15), initial theta to rho ratio of 5 (-r 5), and 28 free parameters spanning 64 atomic time intervals (-p “4+25*2+4+6”).

### Error-corrected D-statistics

The genome of a Yoruba individual (HGDP00927, Meyer et al. 2012) was used as an outgroup to indicate the ancestral allele for the D-statistics. The consensus sequence of the Yoruba genome was generated with ANGSD (Korneliussen et al. 2014), selecting the most common base (-doFasta 2 - doCounts 1) with a minimum base quality of 30 (-minQ 30). Error rates were computed as indicated in the “Error rate estimation” section for 25 genomes (dataset ErrorCorr_Dstat, Dataset S19) used in the test as either H_1_, H_2_ or H_3_. D-statistics computation, error correction and weighted jackknife resampling were performed with ANGSD over 5 Mb blocks (-blockSize 5000000), taking all bases at each position (-doAbbababa2 1) with a minimum quality score of 30 (-minQ 30) following (Soraggi et al. 2018).

### Metagenomic analyses

#### Bacteriome classification

The reads that could not be mapped to the human genome were classified into different taxonomic levels by Kraken2 (Wood et al. 2019) using NCBI’s RefSeq database, which included bacterial, archaeal and viral genomes (downloaded on November 3^rd^, 2017). We then extracted the summary statistics for the counts and assigned hits at the species level, using Kraken-biom (https://github.com/smdabdoub/kraken-biom).

#### Authentication of oral bacteria

We decided to focus on bacteria that could be present in the oral cavity. We first identified the species for which at least 10% of the total reads of the individual samples were assigned, resulting in 25 taxa (Dataset S15). Eight out of the twenty-five taxa have been reported among the oral genomes of the expanded Human Oral Microbiome Database: *Actinomyces* sp. oral taxon 414 strain F0588 (NZ_CP012590.1), *Anaerolineaceae* bacterium oral taxon 439 strain W11661 (NZ_CP017039.1), *Fusobacterium nucleatum* strain Fn12230 (NZ_CP053468.1), *Comamonas testosteroni* strain G1 (NZ_CP067086.1), *Tannerella forsythia* 92A2 (NC_016610.1), *Stenotrophomonas maltophilia* strain NCTC10258 (NZ_LS483377.1), *Acinetobacter lwoffii* strain NCTC5866 (NZ_CAADHN010000001.1), *Pseudomonas stutzeri* strain F2a (NZ_AP024722.1).

We then mapped the non-human reads of the 24 sequenced individuals to the reference genomes of the eight oral bacteria with BWA (Li et al., 2009) aln (version 0.7.13) using a seed length of 32 and a maximum edit distance of 0.04 (with the flags -l 32-n 0.04). To filter the mapped reads, we used samtools view (Danecek et al. 2021) (version 0.1.19) and a mapping quality of 37. To remove clonal duplicates, we used samtools rmdup. The DNA damage patterns were generated using mapDamage2 (Jónsson et al. 2013) with default parameters. We used Circos (Krzywinski et al. 2009) (version 0.69-6) to plot the genome depth of coverage within a window size of 1000 bp.

#### Virome classification

The reads per individual that did not map to the human genome were used as input for DIAMOND (0.9.22, Buchfink et al. 2014) to screen for viruses. DIAMOND was run in its alignment mode (*diamond blastx*) with the RefSeq viral protein database (downloaded on November 9^th^ 2018). The alignments retrieved by DIAMOND were filtered to keep only viruses that infect humans. We used the host information stored in Virus-Host DB to do so (downloaded on September 25^th^ 2019, Mihara et al. 2016). A total of 50 different viral candidates infecting humans were obtained after this step, each candidate had at least one alignment for at least one of the 24 individuals.

The non-human reads of all the individuals were then mapped against each of the 50 viral candidates (Dataset S16) independently using BWA (H. Li et al. 2009) (v 0.7.15) (*aln -l 1024*) setting the threshold for mapping quality at 0 to retain reads mapping within repetitive regions as some viruses had palindromic regions (see below). A threshold of 1% for the BoC (at least 1% of the genome covered by at least one sequenced base) was then applied to further reduce the list of candidates. A total 19 individuals with hits to 15 viruses were retained after this step. The viruses include: adeno-associated virus 2, bufavirus-3, hepatitis B virus, human betaherpesvirus 7, human endogenous retrovirus K113, human parvovirus 4, human parvovirus B19, mamastrovirus 1, Merkel cell polyomavirus, torque teno midi virus 14, torque teno virus 10, torque teno virus 12, torque teno virus 16, torque teno virus 28 and torque teno virus 8 (Fig. 8A).

To qualitatively assess whether the mapped reads were ancient as opposed to resulting from either modern contamination or from a bioinformatic artefact, the genome coverage, the read length distribution, and deamination patterns were computed. The distribution of reads across the genome was computed using BEDTools (Quinlan et al. 2010) (v. 2.29.2) (genomecov-d-ibam). The read length distribution and the deamination patterns were computed using the tool bamdamage from bammds (v. 1, Anna-Sapfo Malaspinas et al. 2014). For all but one of those candidates listed above, (human parvovirus B19, see Fig. 8BC) we were not able to assess whether the data was ancient as there were too few mapped reads to assess their ancient origin. These 14 viruses are therefore deemed candidates for future studies, and we analyzed further the data mapping to human parvovirus B19.

#### Human parvovirus B19 reads and phylogenetic analysis

For six individuals (MN00346, MN00067, MN00021, MN00013, MN00119, and MN00039), Human parvovirus B19 RefSeq genome (NC_000883) was the virus with the highest BoC with values above 5%. For those six, the reads that mapped against the Human parvovirus B19 were BLASTed (v. 2.10.1+ [*blastn*], Altschul et al. 1990) to the nt database-i.e., considering a more extensive database including archaea, eukaryotes and humans-to see if human parvovirus B19 was the best hit for all reads that mapped to this genome. Only two reads (one in MN00067 and the other in MN00346) had as best hit another organism (the Japanese rice fish, *Oryzias latipes*) rather than human parvovirus B19, suggesting that most of the reads mapping to human parvovirus B19 are genuine parvovirus sequenced DNA fragments.

To reconstruct the parvovirus genomes and to determine the most likely genotype, the unmapped reads of the six individuals were then mapped with BWA [v 0.7.15] [*aln -l 1024*] against the three different genotypes reported for Human parvovirus B19. To do so, we compiled a set of 11 reference genomes following (Mühlemann, Margaryan, et al. 2018). This set includes the Human parvovirus B19 genome whose proteins are present in DIAMOND’s database (NC_000883.2), three genomes per genotype (genotype 1: FN669502.1, AF113323.1, DQ357065.1; genotype 2: HQ340602.1, AJ717293.1, DQ333427.1; genotype 3: AY083234.1, NC_004295.1 and AJ249437.1) and an external outgroup, the Bovine parvovirus (NC_001540.1).

The highest BoC among all the individuals and across the 11 Human parvovirus B19 sequences was achieved for the MN00346 data mapped against AY083234.1 (genotype 3) with a BoC value of 83.9% (Fig. 8B). The other individuals have less than 30.1% BoC for any Human parvovirus B19 sequence (Fig. 8B).

Phylogenetic analyses were conducted to infer the genotype of the parvovirus likely to have infected MN00346. A consensus sequence was first obtained using ANGSD (Korneliussen et al. 2014) (with the options-doFasta 2 and-doCounts 1) for the MN00346 data mapping against AY083234.1. The set of 11 public genomes mentioned above, and the reconstructed MN00346 parvovirus genome were aligned with MUSCLE (Edgar 2004). The data were then restricted to the coding sequence (CDS) regions (removing in particular the inverted terminal repeats). Finally, a neighbor-joining phylogenetic tree was reconstructed from this multiple alignment with Seaview (Gouy et al. 2010) (v. 5.0.4) (BIO-NJ algorithm) with default parameters (Fig. 8D).

### Diet characterization

The measured δ^13^C and δ^15^N values from permanent second and third molars (n = 14 out of 20 “Botocudos” and n = 1 Sambaqui, Dataset S18) were used to infer the diet of the individuals. Note that the values from deciduous teeth and permanent first molars were not included in this analysis as those teeth are formed during the gestational and breastfeeding period and tend to show significantly increased nitrogen and carbon stable isotope values, compared to teeth formed in later periods, when the individual can obtain food directly from plants and animals (Katzenberg et al. 1996). Furthermore, the mummy (MN1943) presented a δ15N value far above the results usually found in human tissues, likely caused by an analytical error, and was also excluded from the diet analyses below.

The δ^13^C and δ^15^N values of the 15 ancient Brazilians were then compared to isotopic data from human permanent teeth and bones from different archaeological and contemporaneous locations and contexts in Brazil. The reference set included:

1. 10,000-8,000 BP (Early Holocene) hunter-gatherers from the sites Lapa do Santo (Hermenegildo 2009; Strauss et al. 2016) and Lapa das Boleiras (Hermenegildo 2009) from the Lagoa Santa region in Minas Gerais state.
2. 6,700-4,900 BP (Mid-Holocene) hunter-gatherers from a riverine sambaqui, Sítio Moraes, São Paulo state (Colonese et al. 2014).
3. 3,000-1,000 BP fisher-gatherers from three coastal sites in the Santa Catarina state: Sambaqui do Forte Marechal Luz (Bastos et al. 2014), Sambaqui Jabuticabeira II (Colonese et al. 2014) and Praia da Tapera (Bastos et al. 2015).
4. Present-day individuals from Rio de Janeiro and Brasília cities (Tinoco 2019).

The isotopic values were displayed on two dimensions (δ^13^C vs δ^15^N, Fig. S15).

## References

1. Allentoft, Morten E., Matthew Collins, David Harker, James Haile, Charlotte L. Oskam, Marie L. Hale, Paula F. Campos, et al. 2012. “The Half-Life of DNA in Bone: Measuring Decay Kinetics in 158 Dated Fossils.” Proceedings of the Royal Society B: Biological Sciences 279 (1748): 4724–33. https://doi.org/10.1098/rspb.2012.1745.

2. Allentoft, Morten E, Martin Sikora, Karl-Göran Sjögren, Simon Rasmussen, Morten Rasmussen, Jesper Stenderup, Peter B Damgaard, et al. 2015. “Population Genomics of Bronze Age Eurasia.” Nature 522 (7555): 167–72. https://doi.org/10.1038/nature14507.

3. Altschul, Stephen F., Warren Gish, Webb Miller, Eugene W. Myers, and David J. Lipman. 1990. “Basic Local Alignment Search Tool.” Journal of Molecular Biology 215 (3): 403–10. https://doi.org/10.1016/S0022-2836(05)80360-2.

4. Altshuler, David M., Richard M. Durbin, Gonçalo R. Abecasis, David R. Bentley, Aravinda Chakravarti, Andrew G. Clark, Peter Donnelly, et al. 2012. “An Integrated Map of Genetic Variation from 1,092 Human Genomes.” Nature 491 (7422): 56–65. https://doi.org/10.1038/nature11632.

5. Altshuler, David M., Richard A. Gibbs, Leena Peltonen, David M. Altshuler, Richard A. Gibbs, Leena Peltonen, Emmanouil Dermitzakis, et al. 2010. “Integrating Common and Rare Genetic Variation in Diverse Human Populations.” Nature 467 (7311): 52–58. https://doi.org/10.1038/nature09298.

6. Alves-Silva, Juliana, Magda da Silva Santos, Pedro E.M. Guimarães, Alessandro C.S. Ferreira, Hans-Jürgen Bandelt, Sérgio D.J. Pena, and Vania Ferreira Prado. 2000. “The Ancestry of Brazilian MtDNA Lineages.” The American Journal of Human Genetics 67 (2): 444–61. https://doi.org/10.1086/303004.

7. Arizmendi Cárdenas, Yami Ommar, Samuel Neuenschwander, and Anna-Sapfo Malaspinas. 2021. “Benchmarking Metagenomics Classifiers on Ancient Viral DNA: A Simulation Study.” *BioRxiv*, April, 2021.04.30.442132. https://doi.org/10.1101/2021.04.30.442132.

8. Azevedo, Marta Maria. 2018. “Instituto Socioambiental | Povos Indígenas No Brasil.” 2018. https://pib.socioambiental.org/pt/O_Censo_2010_e_os_Povos_Indigenas.

9. Barbieri, Chiara, Rodrigo Barquera, Leonardo Arias, José R. Sandoval, and Oscar Acosta. 2018. “The Genomic Landscape of Western South America : Andes, Amazonia and Pacific Coast.”

10. Barquera, Rodrigo, Thiseas C. Lamnidis, Aditya Kumar Lankapalli, Arthur Kocher, Diana I. Hernández-Zaragoza, Elizabeth A. Nelson, Adriana C. Zamora-Herrera, et al. 2020. “Origin and Health Status of First-Generation Africans from Early Colonial Mexico.” Current Biology 30 (11): 2078–2091.e11. https://doi.org/10.1016/j.cub.2020.04.002.

11. Bastos, Murilo Q. R., Andrea Lessa, Claudia Rodrigues-Carvalho, Robert H. Tykot, and Roberto Ventura Santos. 2014. “Análise de Isótopos de Carbono e Nitrogênio: A Dieta Antes e Após a Presença de Cerâmica No Sítio Forte Marechal Luz.” Revista Do Museu de Arqueologia e Etnologia, no. 24: 137. https://doi.org/10.11606/issn.2448-1750.revmae.2014.109329.

12. Bastos, Murilo Q. R., Roberto V. Santos, Robert H. Tykot, Sheila M. F. Mendonça de Souza, Claudia Rodrigues-Carvalho, and Andrea Lessa. 2015. “Isotopic Evidences Regarding Migration at the Archeological Site of Praia Da Tapera: New Data to an Old Matter.” Journal of Archaeological Science: Reports 4 (December): 588–95. https://doi.org/10.1016/j.jasrep.2015.10.028.

13. Beall, Clifford J., Alisha G. Campbell, Ann L. Griffen, Mircea Podar, and Eugene J. Leys. 2018. “Genomics of the Uncultivated, Periodontitis-Associated Bacterium Tannerella Sp. BU045 (Oral Taxon 808).” *MSystems*, June. https://doi.org/10.1128/mSystems.00018-18.

14. Bergström, Anders, Shane A. McCarthy, Ruoyun Hui, Mohamed A. Almarri, Qasim Ayub, Petr Danecek, Yuan Chen, et al. 2020. “Insights into Human Genetic Variation and Population History from 929 Diverse Genomes.” Science 367 (6484): eaay5012. https://doi.org/10.1126/science.aay5012.

15. Bieber, Judy. 2014. “Mediation Through Militarization: Indigenous Soldiers and Transcultural Middlemen of the Rio Doce Divisions, Minas Gerais, Brazil, 1808-1850” 71 (2): 227–54.

16. Bolnick, Deborah A., Jennifer A. Raff, Lauren C. Springs, Austin W. Reynolds, and Aida T. Miró-Herrans. 2016. “Native American Genomics and Population Histories.” Annual Review of Anthropology 45 (1): 319–40. https://doi.org/10/gfpn95.

17. Bos, Kirsten I., Kelly M. Harkins, Alexander Herbig, Mireia Coscolla, Nico Weber, Iñaki Comas, Stephen A. Forrest, et al. 2014. “Pre-Columbian Mycobacterial Genomes Reveal Seals as a Source of New World Human Tuberculosis.” Nature 2014 514:7523 514 (7523): 494–97. https://doi.org/10.1038/nature13591.

18. Bravo-Lopez, Miriam, Viridiana Villa-Islas, Carolina Rocha Arriaga, Ana B. Villaseñor-Altamirano, Axel Guzmán-Solís, Marcela Sandoval-Velasco, Julie K. Wesp, et al. 2020. “Paleogenomic Insights into the Red Complex Bacteria Tannerella Forsythia in Pre-Hispanic and Colonial Individuals from Mexico.” Philosophical Transactions of the Royal Society B: Biological Sciences 375 (1812): 20190580. https://doi.org/10.1098/rstb.2019.0580.

19. Bronk Ramsey, Christopher. 2009. “Bayesian Analysis of Radiocarbon Dates.” Radiocarbon 51 (01): 337–60. https://doi.org/10.1017/S0033822200033865.

20. Buchfink, Benjamin, Chao Xie, and Daniel H. Huson. 2014. “Fast and Sensitive Protein Alignment Using DIAMOND.” Nature Methods 12 (1): 59–60. https://doi.org/10.1038/nmeth.3176.

21. Castro e Silva, Marcos Araújo, Tiago Ferraz, Maria Cátira Bortolini, David Comas, and Tábita Hünemeier. 2021. “Deep Genetic Affinity between Coastal Pacific and Amazonian Natives Evidenced by Australasian Ancestry.” Proceedings of the National Academy of Sciences 118 (14): e2025739118. https://doi.org/10.1073/pnas.2025739118.

22. Castro e Silva, Marcos Araújo, Kelly Nunes, Renan Barbosa Lemes, Àlex Mas-Sandoval, Carlos Eduardo Guerra Amorim, Jose Eduardo Krieger, José Geraldo Mill, et al. 2020. “Genomic Insight into the Origins and Dispersal of the Brazilian Coastal Natives.” Proceedings of the National Academy of Sciences of the United States of America 117 (5): 2372–77. https://doi.org/10.1073/pnas.1909075117.

23. Ceballos, Francisco C., Peter K. Joshi, David W. Clark, Michèle Ramsay, and James F. Wilson. 2018. “Runs of Homozygosity: Windows into Population History and Trait Architecture.” Nature Reviews Genetics 19 (4): 220–34. https://doi.org/10.1038/nrg.2017.109.

24. Colombo, Giulia, Luca Traverso, Lucia Mazzocchi, Viola Grugni, Nicola Rambaldi Migliore, Marco Rosario Capodiferro, Gianluca Lombardo, et al. 2022. “Overview of the Americas’ First Peopling from a Patrilineal Perspective: New Evidence from the Southern Continent,” 22.

25. Colonese, André Carlo, Matthew Collins, Alexandre Lucquin, Michael Eustace, Y. Hancock, Raquel De Almeida Rocha Ponzoni, Alice Mora, et al. 2014. “Long-Term Resilience of Late Holocene Coastal Subsistence System in Southeastern South America.” PLoS ONE 9 (4): 1–13. https://doi.org/10.1371/journal.pone.0093854.

26. Core Team, R and R Core Team. 2020. R: A Language and Environment for Statistical Computing. R Foundation for Statistical Computing, Vienna, Austria. Vol. 0. Vienna, Austria: R Foundation for Statistical Computing. https://doi.org/10.1038/sj.hdy.6800737.

27. Cox, Trevor F., and Michael A. A. Cox. 2001. “Multidimensional Scaling,” 308.

28. Damgaard, Peter B, Ashot Margaryan, Hannes Schroeder, Ludovic Orlando, Eske Willerslev, and Morten E Allentoft. 2015. “Improving Access to Endogenous DNA in Ancient Bones and Teeth.” Scientific Reports 5 (January): 11184. https://doi.org/10.1038/srep11184.

29. Danecek, Petr, James K Bonfield, Jennifer Liddle, John Marshall, Valeriu Ohan, Martin O Pollard, Andrew Whitwham, et al. 2021. “Twelve Years of SAMtools and BCFtools.” GigaScience 10 (2): giab008. https://doi.org/10.1093/gigascience/giab008.

30. Edgar, R. C. 2004. “MUSCLE: Multiple Sequence Alignment with High Accuracy and High Throughput.” Nucleic Acids Research 32 (5): 1792–97. https://doi.org/10.1093/nar/gkh340.

31. Ehrenreich, Paul. 2014. Índios Botocudos Do Espírito Santo No Século XIX. Vol. 21.

32. Eisenhofer, Raphael, Hideaki Kanzawa-Kiriyama, Ken-ichi Shinoda, and Laura S. Weyrich. 2020. “Investigating the Demographic History of Japan Using Ancient Oral Microbiota.” Philosophical Transactions of the Royal Society B: Biological Sciences 375 (1812): 20190578. https://doi.org/10/gn4cnq.

33. Escapa, Isabel F., Tsute Chen, Yanmei Huang, Prasad Gajare, Floyd E. Dewhirst, and Katherine P. Lemon. 2018. “New Insights into Human Nostril Microbiome from the Expanded Human Oral Microbiome Database (EHOMD): A Resource for the Microbiome of the Human Aerodigestive Tract.” MSystems 3 (6). https://doi.org/10.1128/MSYSTEMS.00187-18/SUPPL_FILE/SYS006182299ST7.XLSX.

34. Ewens, W.J. 2016. “Effective Population Size.” In Encyclopedia of Evolutionary Biology, 494–97. Elsevier. https://doi.org/10.1016/B978-0-12-800049-6.00025-1.

35. Fernandes, Ricardo, Andrew R. Millard, Marek Brabec, Marie-Josée Nadeau, and Pieter Grootes. 2014. “Food Reconstruction Using Isotopic Transferred Signals (FRUITS): A Bayesian Model for Diet Reconstruction.” PLOS ONE 9 (2): e87436. https://doi.org/10.1371/JOURNAL.PONE.0087436.

36. Fleming-Moran, Millicent, Ricardo V. Santos, and Carlos E. A. Coimbra. 1991. “Blood Pressure Levels of the Suruí and Zoró Indians of the Brazilian Amazon: Group- and Sex-Specific Effects Resulting from Body Composition, Health Status, and Age.” Source 63 (6): 835–61.

37. Fu, Qiaomei, Alissa Mittnik, Philip L F Johnson, Kirsten Bos, Martina Lari, Ruth Bollongino, Chengkai Sun, et al. 2013. “A Revised Timescale for Human Evolution Based on Ancient Mitochondrial Genomes.” Current Biology : CB 23 (7): 553–59. https://doi.org/10.1016/j.cub.2013.02.044.

38. Galetti, Mauro, Raisa Reis Rodarte, Carolina Lima Neves, Marcelo Moreira, and Raul Costa-Pereira. 2016. “Trophic Niche Differentiation in Rodents and Marsupials Revealed by Stable Isotopes.” PLOS ONE 11 (4): e0152494. https://doi.org/10.1371/JOURNAL.PONE.0152494.

39. Galucio, Ana Vilacy, Sérgio Meira, Joshua Birchall, Denny Moore, Nilson Gabas Júnior, Sebastian Drude, Luciana Storto, Gessiane Picanço, and Carmen Reis Rodrigues. 2015. “Genealogical Relations and Lexical Distances within the Tupian Linguistic Family.” Boletim Do Museu Paraense Emílio Goeldi. Ciências Humanas 10 (2): 229–74. https://doi.org/10.1590/1981-81222015000200004.

40. Gamba, Cristina, Eppie R Jones, Matthew D Teasdale, Russell L McLaughlin, Gloria Gonzalez-Fortes, Valeria Mattiangeli, László Domboróczki, et al. 2014. “Genome Flux and Stasis in a Five Millennium Transect of European Prehistory.” Nature Communications 5 (1): 5257. https://doi.org/10.1038/ncomms6257.

41. Goncalves, V. F., Jesper Stenderup, Cláudia Rodrigues-Carvalho, Hilton P Silva, H. Goncalves-Dornelas, A. Liryo, Toomas Kivisild, et al. 2013. “Identification of Polynesian MtDNA Haplogroups in Remains of Botocudo Amerindians from Brazil.” Proceedings of the National Academy of Sciences 110 (16): 6465–69. https://doi.org/10.1073/pnas.1217905110.

42. Gonçalves, Vanessa F, Flavia C Parra, Higgor Gonçalves-Dornelas, Claudia Rodrigues-Carvalho, Hilton P Silva, and Sergio DJ Pena. 2010. “Recovering Mitochondrial DNA Lineages of Extinct Amerindian Nations in Extant Homopatric Brazilian Populations.” Investigative Genetics 1 (1): 13. https://doi.org/10.1186/2041-2223-1-13.

43. Gouy, Manolo, Stéphane Guindon, and Olivier Gascuel. 2010. “Sea View Version 4: A Multiplatform Graphical User Interface for Sequence Alignment and Phylogenetic Tree Building.” Molecular Biology and Evolution 27 (2): 221–24. https://doi.org/10.1093/molbev/msp259.

44. Green, Richard E, Johannes Krause, Adrian W Briggs, Tomislav Maricic, Udo Stenzel, Martin Kircher, Nick Patterson, et al. 2010. “A Draft Sequence of the Neandertal Genome.” *Science (New York*, N.Y.) 328 (5979): 710–22. https://doi.org/10.1126/science.1188021.

45. Green, Richard E., Anna Sapfo Malaspinas, Johannes Krause, Adrian W. Briggs, Philip L. F. Johnson, Caroline Uhler, Matthias Meyer, et al. 2008. “A Complete Neandertal Mitochondrial Genome Sequence Determined by High-Throughput Sequencing.” Cell 134 (3): 416–26. https://doi.org/10.1016/j.cell.2008.06.021.

46. Guzmán-Solís, Axel A., Viridiana Villa-Islas, Miriam J. Bravo-López, Marcela Sandoval-Velasco, Julie K. Wesp, Jorge A. Gómez-Valdés, María de la Luz Moreno-Cabrera, et al. 2021. “Ancient Viral Genomes Reveal Introduction of Human Pathogenic Viruses into Mexico during the Transatlantic Slave Trade.” ELife 10 (August). https://doi.org/10.7554/eLife.68612.

47. Heaton, Timothy J., Peter Köhler, Martin Butzin, Edouard Bard, Ron W. Reimer, William E. N. Austin, Christopher Bronk Ramsey, et al. 2020. “Marine20—The Marine Radiocarbon Age Calibration Curve (0–55,000 Cal BP).” Radiocarbon 62 (4): 779–820. https://doi.org/10.1017/RDC.2020.68.

48. Henn, Brenna M., Laura R. Botigué, Stephan Peischl, Isabelle Dupanloup, Mikhail Lipatov, Brian K. Maples, Alicia R. Martin, et al. 2016. “Distance from Sub-Saharan Africa Predicts Mutational Load in Diverse Human Genomes.” Proceedings of the National Academy of Sciences 113 (4): E440–49. https://doi.org/10.1073/PNAS.1510805112.

49. Hermenegildo, Tiago. 2009. “Reconstituição Da Dieta e Dos Padrões de Subsistência Das Populações Pré-Históricas de Caçadores-Coletores Do Brasil Central Através Da Ecologia Isotópica.” Universidade de São Paulo.

50. Hofstad, Tor. 2006. “The Genus Fusobacterium.” In The Prokaryotes, edited by Martin Dworkin, Stanley Falkow, Eugene Rosenberg, Karl-Heinz Schleifer, and Erko Stackebrandt, 1016–27. New York, NY: Springer New York. https://doi.org/10.1007/0-387-30747-8_51.

51. Hogg, Alan G., Timothy J. Heaton, Quan Hua, Jonathan G. Palmer, Chris SM Turney, John Southon, Alex Bayliss, et al. 2020. “SHCal20 Southern Hemisphere Calibration, 0–55,000 Years Cal BP.” Radiocarbon 62 (4): 759–78. https://doi.org/10.1017/RDC.2020.59.

52. Hughey, Jeffery R., Juan C. Braga, Julio Aguirre, William J. Woelkerling, and Jody M. Webster. 2008. “Analysis of Ancient DNA from Fossil Corallines (Corallinales, Rhodophyta).” Journal of Phycology 44 (2): 374–83. https://doi.org/10.1111/j.1529-8817.2008.00462.x.

53. IBGE. 2010. “Censo Demográfico : 2010 : Características Gerais Dos Indígenas : Resultados Do Universo.” INSTITUTO BRASILEIRO DE GEOGRAFIA E ESTATÍSTICA.

54. Imbelloni, José. 1938. Tabla Clasificatoria de Los Indios; Regiones Biológicas y Grupos Raciales Humanos de América. Physis. https://www.worldcat.org/title/tabla-clasificatoria-de-los-indios-regiones-biologicas-y-grupos-raciales-humanos-de-america/oclc/28305199.

55. Jónsson, Hákon, Aurélien Ginolhac, Mikkel Schubert, Philip L F Johnson, and Ludovic Orlando. 2013. “MapDamage2.0: Fast Approximate Bayesian Estimates of Ancient DNA Damage Parameters.” Bioinformatics 29 (13): 1682–84.

56. Jørsboe, Emil, Kristian Hanghøj, and Anders Albrechtsen. 2017. “FastNGSadmix: Admixture Proportions and Principal Component Analysis of a Single NGS Sample.” Edited by Oliver Stegle. Bioinformatics 33 (19): 3148–50. https://doi.org/10.1093/bioinformatics/btx474.

57. Kanindé Associação de Defesa Etnoambiental, Organização Metareilá do Povo Indígena Paiter, and Betty Mindlin. 2021. “Instituto Socioambiental | Povos Indígenas No Brasil.” 2021. https://pib.socioambiental.org/pt/Povo:Surui_Paiter.

58. Karolchik, Donna, Angela S Hinrichs, Terrence S Furey, Krishna M Roskin, Charles W Sugnet, David Haussler, and W James Kent. 2004. “The UCSC Table Browser Data Retrieval Tool.” Nucleic Acids Research 32 (Database issue): D493-6. https://doi.org/10.1093/nar/gkh103.

59. Katoh, Kazutaka, and Daron M. Standley. 2013. “MAFFT Multiple Sequence Alignment Software Version 7: Improvements in Performance and Usability.” Molecular Biology and Evolution 30 (4): 772–80. https://doi.org/10.1093/molbev/mst010.

60. Katzenberg, M Anne, D Ann Herring, and Shelley R Saunders. 1996. “Weaning and Infant Mortality: Evaluating the Skeletal Evidence.” YEARBOOK OF PHYSICAL ANTHROPOLOGY 39: 177–99. https://doi.org/10.1002/(SICI)1096-8644(1996)23.

61. Kazarina, Alisa, Elina Petersone-Gordina, Janis Kimsis, Jevgenija Kuzmicka, Pawel Zayakin, Žans Griškjans, Guntis Gerhards, and Renate Ranka. 2021. “The Postmedieval Latvian Oral Microbiome in the Context of Modern Dental Calculus and Modern Dental Plaque Microbial Profiles.” Genes 12 (2): 309. https://doi.org/10/gn4cns.

62. Kircher, Martin, Susanna Sawyer, and Matthias Meyer. 2012. “Double Indexing Overcomes Inaccuracies in Multiplex Sequencing on the Illumina Platform.” Nucleic Acids Research 40 (1): 1–8.

63. Kirin, Mirna, Ruth McQuillan, Christopher S. Franklin, Harry Campbell, Paul M. McKeigue, and James F. Wilson. 2010. “Genomic Runs of Homozygosity Record Population History and Consanguinity.” PLOS ONE 5 (11): 1–7. https://doi.org/10.1371/journal.pone.0013996.

64. Kivisild, Toomas. 2017. “The Study of Human Y Chromosome Variation through Ancient DNA.” *Human Genetics*, March, 1–18. https://doi.org/10.1007/s00439-017-1773-z.

65. Korneliussen, Thorfinn Sand, Anders Albrechtsen, and Rasmus Nielsen. 2014. “ANGSD: Analysis of Next Generation Sequencing Data.” BMC Bioinformatics 15 (1): 356.

66. Krzywinski, Martin, Jacqueline Schein, Inanç Birol, Joseph Connors, Randy Gascoyne, Doug Horsman, Steven J. Jones, and Marco A. Marra. 2009. “Circos: An Information Aesthetic for Comparative Genomics.” Genome Research 19 (9): 1639–45. https://doi.org/10.1101/gr.092759.109.

67. Lacerda, F., and R. Peixoto. 1876. “Contribuições Para o Estudo Antropológico Das Raças Indígenas Do Brazil.” In . Archivos do Museu Nacional do Rio de Janeiro.

68. Langfur, Hal. 2002. “Uncertain Refuge: Frontier Formation and the Origins of the Botocudo War in Late Colonial Brazil.” Hispanic American Historical Review 82 (2): 215–56. https://doi.org/10.1215/00182168-82-2-215.

69. León, De, Toetik Koesbardiati, John David Weissmann, Carlos S. Reyna-blanco, Gen Suwa, Osamu Kondo, Sapfo Malaspinas, et al. 2018. “Labyrinth Is an Indicator of Population History and Dispersal from Africa.” Proceedings of the National Academy of Sciences 115 (23): E5429–E5429. https://doi.org/10.1073/pnas.1808125115.

70. Li, H., and R. Durbin. 2009. “Fast and Accurate Short Read Alignment with Burrows-Wheeler Transform.” Bioinformatics 25 (14): 1754–60.

71. Li, Heng, and Richard Durbin. 2011. “Inference of Human Population History from Individual Whole-Genome Sequences.” Nature 475 (7357): 493–96. https://doi.org/10.1038/nature10231.

72. Lipson, Mark, and David Reich. 2017. “A Working Model of the Deep Relationships of Diverse Modern Human Genetic Lineages Outside of Africa.” Molecular Biology and Evolution 34 (4): 889–902. https://doi.org/10.1093/MOLBEV/MSW293.

73. Majander, Kerttu, Saskia Pfrengle, Arthur Kocher, Denise Kü, Johannes Krause, Verena J. Schuenemann Correspondence, Judith Neukamm, et al. 2020. “Ancient Bacterial Genomes Reveal a High Diversity of Treponema Pallidum Strains in Early Modern Europe.” Current Biology 30: 3788–3803.e10. https://doi.org/10.1016/j.cub.2020.07.058.

74. Malaspinas, Anna-Sapfo, Ole Tange, José Víctor Moreno-Mayar, Morten Rasmussen, Michael DeGiorgio, Yong Wang, Cristina E. Valdiosera, Gustavo Politis, Eske Willerslev, and Rasmus Nielsen. 2014. “Bammds: A Tool for Assessing the Ancestry of Low- Depth Whole-Genome Data Using Multidimensional Scaling (MDS).” Bioinformatics 30 (20): 2962–64. https://doi.org/10.1093/bioinformatics/btu410.

75. Malaspinas, Anna-Sapfo, Michael C. Westaway, Craig Muller, Vitor C. Sousa, Oscar Lao, Isabel Alves, Anders Bergström, et al. 2016. “A Genomic History of Aboriginal Australia.” Nature 538 (7624): 207–14. https://doi.org/10.1038/nature18299.

76. Malaspinas, A.-S, O Lao, H Schroeder, M Rasmussen, M Raghavan, I Moltke, P F Campos, et al. 2014. “Two Ancient Human Genomes Reveal Polynesian Ancestry among the Indigenous Botocudos of Brazil.” Current Biology 29 (8): 12711–16. https://doi.org/10.1016/j.cub.2014.09.035.

77. Mallick, Swapan, Heng Li, Mark Lipson, Iain Mathieson, Melissa Gymrek, Fernando Racimo, Mengyao Zhao, et al. 2016. “The Simons Genome Diversity Project: 300 Genomes from 142 Diverse Populations.” Nature 538 (7624): 201–6.

78. Mann, Allison E., Susanna Sabin, Kirsten Ziesemer, Åshild J. Vågene, Hannes Schroeder, Andrew T. Ozga, Krithivasan Sankaranarayanan, et al. 2018. “Differential Preservation of Endogenous Human and Microbial DNA in Dental Calculus and Dentin.” Scientific Reports 2018 8:1 8 (1): 1–15. https://doi.org/10.1038/s41598-018-28091-9.

79. McKenna, Aaron, Matthew Hanna, Eric Banks, Andrey Sivachenko, Kristian Cibulskis, Andrew Kernytsky, Kiran Garimella, et al. 2010. “The Genome Analysis Toolkit: A MapReduce Framework for Analyzing next-Generation DNA Sequencing Data.” Genome Research 20 (9): 1297–1303. https://doi.org/10.1101/gr.107524.110.

80. McQuillan, Ruth, Anne-Louise Leutenegger, Rehab Abdel-Rahman, Christopher S. Franklin, Marijana Pericic, Lovorka Barac-Lauc, Nina Smolej-Narancic, et al. 2008. “Runs of Homozygosity in European Populations.” The American Journal of Human Genetics 83 (3): 359–72. https://doi.org/10.1016/j.ajhg.2008.08.007.

81. Meyer, Matthias, Martin Kircher, Marie Theres Gansauge, Heng Li, Fernando Racimo, Swapan Mallick, Joshua G. Schraiber, et al. 2012. “A High-Coverage Genome Sequence from an Archaic Denisovan Individual.” Science 338 (6104): 222–26. https://doi.org/10.1126/science.1224344.

82. Mihara, Tomoko, Yosuke Nishimura, Yugo Shimizu, Hiroki Nishiyama, Genki Yoshikawa, Hideya Uehara, Pascal Hingamp, Susumu Goto, and Hiroyuki Ogata. 2016. “Linking Virus Genomes with Host Taxonomy.” Viruses 8 (3). https://doi.org/10.3390/v8030066.

83. Mindlin, Betty. 1985. “Nós Paiter : Os Suruí de Rondônia.” www.etnolinguistica.org.

84. Missagia De Mattos, Izabel. 2017. “A Guerra Ofensiva Aos ‘Botocudos Antropófagos’ Nas Minas Oitocentistas e Seus Significados Para a Nacionalidade Brasileira Em Formação: Uma Abordagem Comparativa,” 87–105.

85. Monteiro, Maria Elizabeth Brêa. 1984. “Relatório Sobre Os Índios Karitiana, Estado de Rondônia.” Minter-FUNAI. https://acervo.socioambiental.org/sites/default/files/documents/KTD00007.pdf https://acervo.socioambiental.org/acervo/documentos/relatorio-sobre-os-indios-karitiana-estado-de-rondonia.

86. Moore, D. 2006. “Brazil: Language Situation.” *Encyclopedia of Language & Linguistics*, January, 117–28. https://doi.org/10.1016/B0-08-044854-2/01855-1.

87. Moreno-Mayar, J. Víctor, Thorfinn Sand Korneliussen, Jyoti Dalal, Gabriel Renaud, Anders Albrechtsen, Rasmus Nielsen, and Anna-Sapfo Malaspinas. 2020. “A Likelihood Method for Estimating Present-Day Human Contamination in Ancient Male Samples Using Low-Depth X-Chromosome Data.” Edited by Russell Schwartz. Bioinformatics 36 (3): 828–41. https://doi.org/10.1093/bioinformatics/btz660.

88. Moreno-Mayar, J Víctor, Ben A Potter, Lasse Vinner, Matthias Steinrücken, Simon Rasmussen, Jonathan Terhorst, John A Kamm, et al. 2018. “Terminal Pleistocene Alaskan Genome Reveals First Founding Population of Native Americans.” Nature 553 (7687): 203–7. https://doi.org/10.1038/nature25173.

89. Moreno-Mayar, J. Víctor, Lasse Vinner, Peter de Barros Damgaard, Constanza de la Fuente, Jeffrey Chan, Jeffrey P. Spence, Morten E. Allentoft, et al. 2018. “Early Human Dispersals within the Americas.” Science 362 (6419): eaav2621. https://doi.org/10.1126/science.aav2621.

90. Mühlemann, Barbara, Terry C Jones, Peter De Barros Damgaard, Morten E Allentoft, Irina Shevnina, Andrey Logvin, Bazartseren Boldgiv, et al. 2018. “Ancient Hepatitis B Viruses from the Bronze Age to the Medieval Period.” https://doi.org/10.1038/s41586-018-0097-z.

91. Mühlemann, Barbara, Ashot Margaryan, Peter de Barros Damgaard, Morten E. Allentoft, Lasse Vinner, Anders J. Hansen, Andrzej Weber, et al. 2018. “Ancient Human Parvovirus B19 in Eurasia Reveals Its Long-Term Association with Humans.” Proceedings of the National Academy of Sciences 115 (29): 7557–62. https://doi.org/10.1073/pnas.1804921115.

92. Neel, J. V., and F. M. Salzano. 1967. “Further Studies on the Xavante Indians. X. Some Hypotheses-Generalizations Resulting from These Studies.” American Journal of Human Genetics 19 (4): 554.

93. Neves, Pedro Da-gloria Walter A. 2017. Archaeological and Paleontological Research in Lagoa Santa. https://doi.org/10.1007/978-3-319-57466-0.

94. Nielsen, Rasmus, Joshua M. Akey, Mattias Jakobsson, Jonathan K. Pritchard, Sarah Tishkoff, and Eske Willerslev. 2017. “Tracing the Peopling of the World through Genomics.” Nature 541 (7637): 302–10. https://doi.org/10.1038/nature21347.

95. Oliveira, Tatiana Gonçalves. 2016. “O Aldeamento Dos Índios De Itambacuri E A Política Indigenista Na Província De Minas Gerais (1873-1889).” PPGH, UFJF.

96. Orlando, Ludovic, Aurélien Ginolhac, Guojie Zhang, Duane Froese, Anders Albrechtsen, Mathias Stiller, Mikkel Schubert, et al. 2013. “Recalibrating Equus Evolution Using the Genome Sequence of an Early Middle Pleistocene Horse.” Nature 499 (7456): 74– 78. https://doi.org/10.1038/nature12323.

97. Ottoni, Claudio, Dušan Bori, Olivia Cheronet, Vitale Sparacello, Irene Dori, Alfredo Coppa, Dragana Antonovi C. H, et al. n.d. “Tracking the Transition to Agriculture in Southern Europe through Ancient DNA Analysis of Dental Calculus.” https://doi.org/10.1073/pnas.2102116118/-/DCSupplemental.

98. Pagliaro, H, M Azevedo, and R V Santos. 2005. Demografia Dos Povos Indígenas No Brasil: Um Panorama Crítico. Editora FIOCRUZ. https://doi.org/10.7476/9788575412541.

99. Palleroni, Norberto J. 2010. “The Pseudomonas Story.” Environmental Microbiology 12 (6): 1377–83. https://doi.org/10.1111/j.1462-2920.2009.02041.x.

100. Paraíso, Maria Hilda B. 1992. “Os Botocudos e Sua Trajetória Histórica.” In História Dos Índios No Brasil, edited by Manuela Carneiro da Cunha, 413–30. São Paulo: Companhia das Letras.

101. Patterson, Nick, Priya Moorjani, Yontao Luo, Swapan Mallick, Nadin Rohland, Yiping Zhan, Teri Genschoreck, Teresa Webster, and David Reich. 2012. “Ancient Admixture in Human History.” Genetics 192 (3): 1065–93. https://doi.org/10.1534/genetics.112.145037.

102. Paul, Rivet. 1942. Les Origines de l’Homme Américain. Les Éditions de l’Arbre.

103. Pemberton, Trevor J., Devin Absher, Marcus W. Feldman, Richard M. Myers, Noah A. Rosenberg, and Jun Z. Li. 2012. “Genomic Patterns of Homozygosity in Worldwide Human Populations.” American Journal of Human Genetics 91 (2): 275–92. https://doi.org/10.1016/j.ajhg.2012.06.014.

104. Pezo-Lanfranco, Luis, Sabine Eggers, Cecilia Petronilho, Alice Toso, Dione da Rocha Bandeira, Matthew Von Tersch, Adriana M. P. dos Santos, Beatriz Ramos da Costa, Roberta Meyer, and André Carlo Colonese. 2018. “Middle Holocene Plant Cultivation on the Atlantic Forest Coast of Brazil?” Royal Society Open Science 5 (9). https://doi.org/10.1098/RSOS.180432.

105. Philips, Anna, Ireneusz Stolarek, Bogna Kuczkowska, Anna Juras, Luiza Handschuh, Janusz Piontek, Piotr Kozlowski, and Marek Figlerowicz. 2017. “Comprehensive Analysis of Microorganisms Accompanying Human Archaeological Remains.” GigaScience 6 (7): 1–13. https://doi.org/10/gbg8vd.

106. Phillips, Nicky. 2019. “Indigenous Groups Look to Ancient DNA to Bring Their Ancestors Home.” Nature 568 (7752): 294–97. https://doi.org/10.1038/D41586-019-01167-W.

107. Pickrell, Joseph K., and David Reich. 2014. “Toward a New History and Geography of Human Genes Informed by Ancient DNA.” Trends in Genetics 30 (9): 377–89. https://doi.org/10/f6hqjz.

108. Posth, Cosimo, Nathan Nakatsuka, Iosif Lazaridis, Pontus Skoglund, Swapan Mallick, Thiseas C. Lamnidis, Nadin Rohland, et al. 2018. “Reconstructing the Deep Population History of Central and South America.” Cell 175 (5): 1185–1197.e22. https://doi.org/10.1016/j.cell.2018.10.027.

109. Poznik, G. David. 2016. “Identifying Y-Chromosome Haplogroups in Arbitrarily Large Samples of Sequenced or Genotyped Men.” BioarXiv, 1–5. https://doi.org/10.1101/088716.

110. Pritchard, J. K., M. Stephens, and P. Donnelly. 2000. “Inference of Population Structure Using Multilocus Genotype Data.” Genetics 155 (2): 945–59. https://doi.org/10.1111/j.1471-8286.2007.01758.x.

111. Prüfer, Kay, Fernando Racimo, Nick Patterson, Flora Jay, Sriram Sankararaman, Susanna Sawyer, Anja Heinze, et al. 2014. “The Complete Genome Sequence of a Neanderthal from the Altai Mountains.” Nature 505 (7481): 43–49. https://doi.org/10.1038/nature12886.

112. Pucciarelli, Héctor M., Marina L. Sardi, José C. Jimenez López and Carlos Serrano Sanchez,. 2003. “Early Peopling and Evolutionary Diversification in America.” Quaternary International 109–110 (January): 123–32. https://doi.org/10.1016/S1040-6182(02)00208-2.

113. Purcell, Shaun, Benjamin Neale, Kathe Todd-Brown, Lori Thomas, Manuel A R Ferreira, David Bender, Julian Maller, et al. 2007. “PLINK: A Tool Set for Whole-Genome Association and Population-Based Linkage Analyses.” American Journal of Human Genetics 81 (3): 559–75. https://doi.org/10.1086/519795.

114. Qiu, Jianming, Maria Söderlund-Venermo, and Neal S. Young. 2017. “Human Parvoviruses.” Clinical Microbiology Reviews 30 (1): 43– 113. https://doi.org/10.1128/CMR.00040-16.

115. Quinlan, Aaron R, and Ira M Hall. 2010. “BEDTools: A Flexible Suite of Utilities for Comparing Genomic Features.” *Bioinformatics (Oxford*, England) 26 (6): 841–42. https://doi.org/10.1093/bioinformatics/btq033.

116. Raghavan, M., M. Steinrucken, K. Harris, S. Schiffels, S. Rasmussen, M. DeGiorgio, A. Albrechtsen, et al. 2015. “Genomic Evidence for the Pleistocene and Recent Population History of Native Americans.” Science 349 (6250): aab3884–aab3884. https://doi.org/10.1126/science.aab3884.

117. Raghavan, Maanasa, Michael DeGiorgio, Anders Albrechtsen, Ida Moltke, Pontus Skoglund, Thorfinn S. Korneliussen, Bjarne Grønnow, et al. 2014. “The Genetic Prehistory of the New World Arctic.” Science 345 (6200). https://doi.org/10.1126/science.1255832.

118. Ramallo, Virginia, Rafael Bisso-Machado, Claudio Bravi, Michael D. Coble, Francisco M. Salzano, Tábita Hünemeier, and Maria Cátira Bortolini. 2013. “Demographic Expansions in South America: Enlightening a Complex Scenario with Genetic and Linguistic Data.” American Journal of Physical Anthropology 150 (3): 453–63. https://doi.org/10.1002/ajpa.22219.

119. Rascovan, Nicolás, Karl Göran Sjögren, Kristian Kristiansen, Rasmus Nielsen, Eske Willerslev, Christelle Desnues, and Simon Rasmussen. 2019. “Emergence and Spread of Basal Lineages of Yersinia Pestis during the Neolithic Decline.” Cell 176 (1–2): 295–305.e10. https://doi.org/10.1016/J.CELL.2018.11.005.

120. Rasmussen, Morten, Sarah L. Anzick, Michael R. Waters, Pontus Skoglund, Michael DeGiorgio, Thomas W. Stafford, Simon Rasmussen, et al. 2014. “The Genome of a Late Pleistocene Human from a Clovis Burial Site in Western Montana.” Nature 506 (7487): 225–29. https://doi.org/10.1038/nature13025.

121. Rasmussen, Morten, Martin Sikora, Anders Albrechtsen, Thorfinn Sand Korneliussen, J. Víctor Moreno-Mayar, G. David Poznik, Christoph P. E. Zollikofer, et al. 2015. “The Ancestry and Affiliations of Kennewick Man.” Nature, 1–10.

122. Reimer, Paula, Stephen Hoper, James Mcdonald, Ron Reimer, Svetlana Svyatko, and Michelle Thompson. 2015. “The Queen’s University, Belfast Laboratory Protocols Used Fro AMS Radiocarbon Dating at the 14Chrono Centre Scientific Dating Report.” Scientific Dating Report Copyright English Heritage 5.

123. Ryan, Robert P., Sebastien Monchy, Massimiliano Cardinale, Safiyh Taghavi, Lisa Crossman, Matthew B. Avison, Gabriele Berg, Daniel van der Lelie, and J. Maxwell Dow. 2009. “The Versatility and Adaptation of Bacteria from the Genus Stenotrophomonas.” Nature Reviews Microbiology 7 (7): 514–25. https://doi.org/10.1038/nrmicro2163.

124. Scheel-Ybert, Rita, Sabine Eggers, Veronica Wesolowski, and Paulo DeBlasis. 2003. “Novas Perspectivas Na Reconstituição Do Modo de Vida Dos Sambaquieiros: Uma Abordagem Multidisciplinar.” Revista de Arqueologia 16 (1): 109–37. https://doi.org/10.24885/sab.v16i1.182.

125. Scheib, C. L., Hongjie Li, Tariq Desai, Vivian Link, Christopher Kendall, Genevieve Dewar, Peter William Griffith, et al. 2018. “Ancient Human Parallel Lineages within North America Contributed to a Coastal Expansion.” Science 360 (6392): 1024–27. https://doi.org/10.1126/science.aar6851.

126. Schroeder, Hannes, Martin Sikora, Shyam Gopalakrishnan, Lara M. Cassidy, Pierpaolo Maisano Delser, Marcela Sandoval Velasco, Joshua G. Schraiber, et al. 2018. “Origins and Genetic Legacies of the Caribbean Taino.” Proceedings of the National Academy of Sciences of the United States of America 115 (10): 2341–46. https://doi.org/10.1073/pnas.1716839115.

127. Schubert, Mikkel, Aurelien Ginolhac, Stinus Lindgreen, John F Thompson, Khaled A S Al-Rasheid, Eske Willerslev, Anders Krogh, and Ludovic Orlando. 2012. “Improving Ancient DNA Read Mapping against Modern Reference Genomes.” BMC Genomics 13 (1): 178. https://doi.org/10.1186/1471-2164-13-178.

128. Schubert, Mikkel, Stinus Lindgreen, and Ludovic Orlando. 2016. “AdapterRemoval v2: Rapid Adapter Trimming, Identification, and Read Merging.” BMC Research Notes 9 (1): 88.

129. Sikora, Martin, Andaine Seguin-Orlando, Vitor C. Sousa, Anders Albrechtsen, Thorfinn Korneliussen, Amy Ko, Simon Rasmussen, et al. 2017. “Ancient Genomes Show Social and Reproductive Behavior of Early Upper Paleolithic Foragers.” Science 358 (6363): 659–62. https://doi.org/10.1126/science.aao1807.

130. Silverman, Helaine, and William H. Isbell, eds. 2008. The Handbook of South American Archaeology. The Handbook of South American Archaeology. Springer New York. https://doi.org/10.1007/978-0-387-74907-5.

131. Skoglund, Pontus, Swapan Mallick, Maria Cátira Bortolini, Niru Chennagiri, Tábita Hünemeier, Maria Luiza Petzl-Erler, Francisco Mauro Salzano, Nick Patterson, and David Reich. 2015. “Genetic Evidence for Two Founding Populations of the Americas.” Nature 525 (7567): 104–8. https://doi.org/10.1038/nature14895.

132. Skoglund, Pontus, Helena Malmström, Ayça Omrak, Maanasa Raghavan, Cristina Valdiosera, Torsten Günther, Per Hall, et al. 2014. “Genomic Diversity and Admixture Differs for Stone-Age Scandinavian Foragers and Farmers.” Science 344 (6185): 747–50. https://doi.org/10.1126/science.1253448.

133. Skoglund, Pontus, and David Reich. 2016. “A Genomic View of the Peopling of the Americas.” Current Opinion in Genetics and Development 41: 27–35. https://doi.org/10.1016/j.gde.2016.06.016.

134. Skoglund, Pontus, Jan Stor\a a, Anders Götherström, and Mattias Jakobsson. 2013. “Accurate Sex Identi Fi Cation of Ancient Human Remains Using DNA Shotgun Sequencing.” Journal of Archaeological Science 40 (12): 4477–82.

135. Skotte, Line, Thorfinn Sand Korneliussen, and Anders Albrechtsen. 2013. “Estimating Individual Admixture Proportions from Next Generation Sequencing Data.” Genetics 195 (3): 693–702. https://doi.org/10.1534/genetics.113.154138.

136. Smith, Colin I., Andrew T. Chamberlain, Michael S. Riley, Chris Stringer, and Matthew J. Collins. 2003. “The Thermal History of Human Fossils and the Likelihood of Successful DNA Amplification.” Journal of Human Evolution 45 (3): 203–17. https://doi.org/10.1016/S0047-2484(03)00106-4.

137. Soraggi, Samuele, Carsten Wiuf, and Anders Albrechtsen. 2018. “Powerful Inference with the D-Statistic on Low-Coverage Whole- Genome Data.” G3&#58; Genes|Genomes|Genetics 8 (2): 551–66. https://doi.org/10.1534/g3.117.300192.

138. Steward, J. H. 1949. The Native Population of South America. Steward, J. H. (1949). The native population of South America. US Government Printing Office.

139. Storto, Luciana, and Felipe Ferreira Vander Velden. 2021. “Karitiana- Indigenous Peoples in Brazil.” April 1, 2021. https://pib.socioambiental.org/en/Povo:Karitiana#Denomination_and_population.

140. Strauss, André, Rodrigo Elias Oliveira, Ximena S. Villagran, Danilo V. Bernardo, Domingo C. Salazar-García, Marcos César Bissaro, Francisco Pugliese, et al. 2016. “Early Holocene Ritual Complexity in South America: The Archaeological Record of Lapa Do Santo (East-Central Brazil).” Antiquity 90 (354): 1454–73. https://doi.org/10.15184/aqy.2016.220.

141. Tinoco, Rachel Lima Ribeiro. 2019. “Análise Isotópica Em Dentes de Brasileiros Contemporâneos Com Dieta e Origem Conhecidas: Interpretações Forenses e Arqueológicas.” Universidade Federal do Rio de Janeiro.

142. Towner, K. J., E. Bergogne-Bérézin, and C. A. Fewson, eds. 1991. The Biology of Acinetobacter: Taxonomy, Clinical Importance, Molecular Biology, Physiology, Industrial Relevance. Boston, MA: Springer US. https://doi.org/10.1007/978-1-4899-3553-3.

143. Vågene, Åshild J., Alexander Herbig, Michael G. Campana, Nelly M. Robles García, Christina Warinner, Susanna Sabin, Maria A. Spyrou, et al. 2018. “Salmonella Enterica Genomes from Victims of a Major Sixteenth-Century Epidemic in Mexico.” Nature Ecology & Evolution 2018 2:3 2 (3): 520–28. https://doi.org/10.1038/s41559-017-0446-6.

144. Velden, Felipe Ferreira Vander, 1978-. 2004. “Por Onde o Sangue Circula : Os Karitiana e a Intervenção Biomedica.” http://repositorio.unicamp.br/jspui/handle/REPOSIP/279104.

145. Velsko, Irina M., James A. Fellows Yates, Franziska Aron, Richard W. Hagan, Laurent A. F. Frantz, Louise Loe, Juan Bautista Rodriguez Martinez, et al. 2019. “Microbial Differences between Dental Plaque and Historic Dental Calculus Are Related to Oral Biofilm Maturation Stage.” Microbiome 7 (1): 1–20. https://doi.org/10.1186/s40168-019-0717-3.

146. Wada, Gregory H, Jelmer W Eerkens, Beth Shapiro, and Ruth V Nichols. 2018. “INSIGHTS INTO THE ANCIENT ORAL MICROBIOME: SURVEY OF MICROBIAL DIVERSITY FROM THE DENTAL CALCULUS OF TWO PRECONTACT SITES BY THE SAN FRANCISCO BAY” 32: 16.

147. Wagner, Jennifer K., Chip Colwell, Katrina G. Claw, Anne C. Stone, Deborah A. Bolnick, John Hawks, Kyle B. Brothers, and Nanibaa’ A. Garrison. 2020. “Fostering Responsible Research on Ancient DNA.” American Journal of Human Genetics 107 (2): 183–95. https://doi.org/10/gn4c5q.

148. Wahlund, Sten. 1928. “ZUSAMMENSETZUNG VON POPULATIONEN UND KORRELATIONSERSCHEINUNGEN VOM STANDPUNKT DER VERERBUNGSLEHRE AUS BETRACHTET.” Hereditas 11 (1): 65–106. https://doi.org/10.1111/j.1601-5223.1928.tb02483.x.

149. Walker, Robert S., Søren Wichmann, Thomas Mailund, and Curtis J. Atkisson. 2012. “Cultural Phylogenetics of the Tupi Language Family in Lowland South America.” PLOS ONE 7 (4): e35025. https://doi.org/10.1371/JOURNAL.PONE.0035025.

150. Warinner, Christina, João F. Matias Matias Rodrigues, Rounak Vyas, Christian Trachsel, Natallia Shved, Jonas Grossmann, Anita Radini, et al. 2014. “Pathogens and Host Immunity in the Ancient Human Oral Cavity.” Nature Genetics 46 (4): 336–44. https://doi.org/10.1038/ng.2906.

151. Wesolowski, Verônica, Sheila Maria Ferraz Mendonça de Souza, Karl J. Reinhard, and Gregório Ceccantini. 2010. “Evaluating Microfossil Content of Dental Calculus from Brazilian Sambaquis.” Journal of Archaeological Science 37 (6): 1326–38. https://doi.org/10.1016/J.JAS.2009.12.037.

152. Wied, Maximilian. 1820. Reise Nach Brasilien in Den Jahren 1815 Bis 1817 /. Reise Nach Brasilien in Den Jahren 1815 Bis 1817 /. Gedruckt und verlegt bey H.L. Brönner; https://doi.org/10.5962/bhl.title.85967.

153. Willems, Anne, and Paul De Vos. 2006. “Comamonas.” In The Prokaryotes: Volume 5: Proteobacteria: Alpha and Beta Subclasses, edited by Martin Dworkin, Stanley Falkow, Eugene Rosenberg, Karl-Heinz Schleifer, and Erko Stackebrandt, 723–36. New York, NY: Springer. https://doi.org/10.1007/0-387-30745-1_31.

154. Willerslev, Eske, and David J. Meltzer. 2021. “Peopling of the Americas as Inferred from Ancient Genomics.” Nature 594 (November 2020). https://doi.org/10.1038/s41586-021-03499-y.

155. Wollstein, Andreas, Oscar Lao, Christian Becker, Silke Brauer, Ronald J. Trent, Peter Nürnberg, Mark Stoneking, and Manfred Kayser. 2010. “Demographic History of Oceania Inferred from Genome-Wide Data.” Current Biology 20 (22): 1983–92. https://doi.org/10.1016/j.cub.2010.10.040.

156. Wood, Derrick E., Jennifer Lu, and Ben Langmead. 2019. “Improved Metagenomic Analysis with Kraken 2.” Genome Biology 20 (1). https://doi.org/10.1186/s13059-019-1891-0.

157. Wright, Joanne L., Sally Wasef, Tim H. Heupink, Michael C. Westaway, Simon Rasmussen, Colin Pardoe, Gudju Gudju Fourmile, et al. 2018. “Ancient Nuclear Genomes Enable Repatriation of Indigenous Human Remains.” Science Advances 4 (12). https://doi.org/10.1126/SCIADV.AAU5064.

158. Xing, Jinchuan, W. Scott Watkins, Adam Shlien, Erin Walker, Chad D. Huff, David J. Witherspoon, Yuhua Zhang, et al. 2010. “Toward a More Uniform Sampling of Human Genetic Diversity: A Survey of Worldwide Populations by High-Density Genotyping.” Genomics 96 (4): 199–210. https://doi.org/10.1016/j.ygeno.2010.07.004.

159. Yengo, Loic, Naomi R. Wray, and Peter M. Visscher. 2019. “Extreme Inbreeding in a European Ancestry Sample from the Contemporary UK Population.” Nature Communications 10 (1): 3719. https://doi.org/10.1038/s41467-019-11724-6.

