## Appendix for "Indigenous peoples in eastern Brazil: insights from 19^th^ century genomes and metagenomes"

Figures S1 to S15
Legends for Datasets S1 to S19

Other supplementary materials for this manuscript include the following:
Datasets S1 to S19

### *Figures (S1 to S15)*

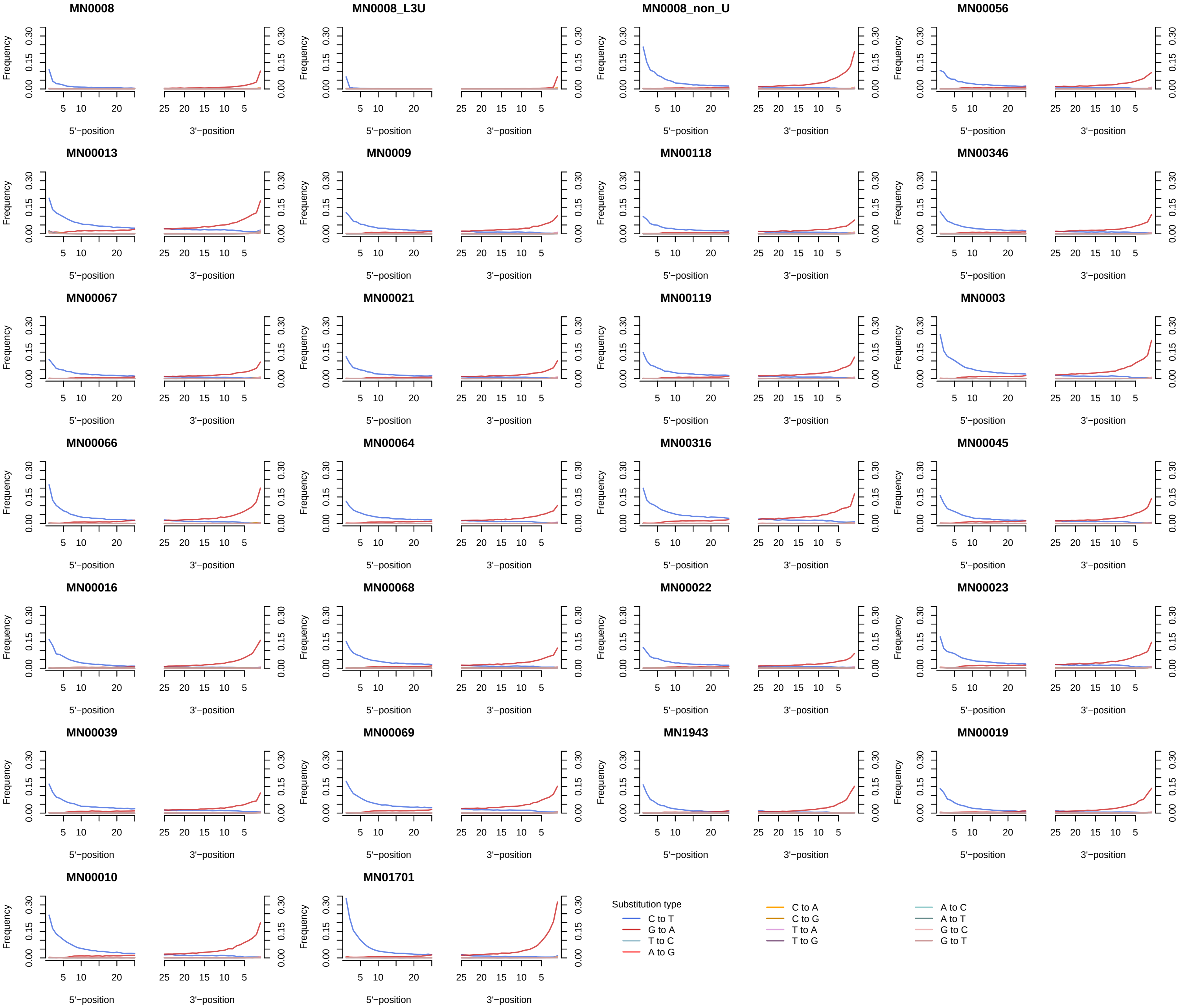


**Figure S1. Substitution frequencies at the read termini across 24 individual samples.** For the individual sample MN0008, substitutions are shown for the data of all the libraries (MN0008), as well as the USER-treated only (MN0008_L3U) and the standard non-USER treated libraries (MN0008_non_U). See Dataset S3 for the plots corresponding to the substitution rates for each of the libraries sequenced per individual

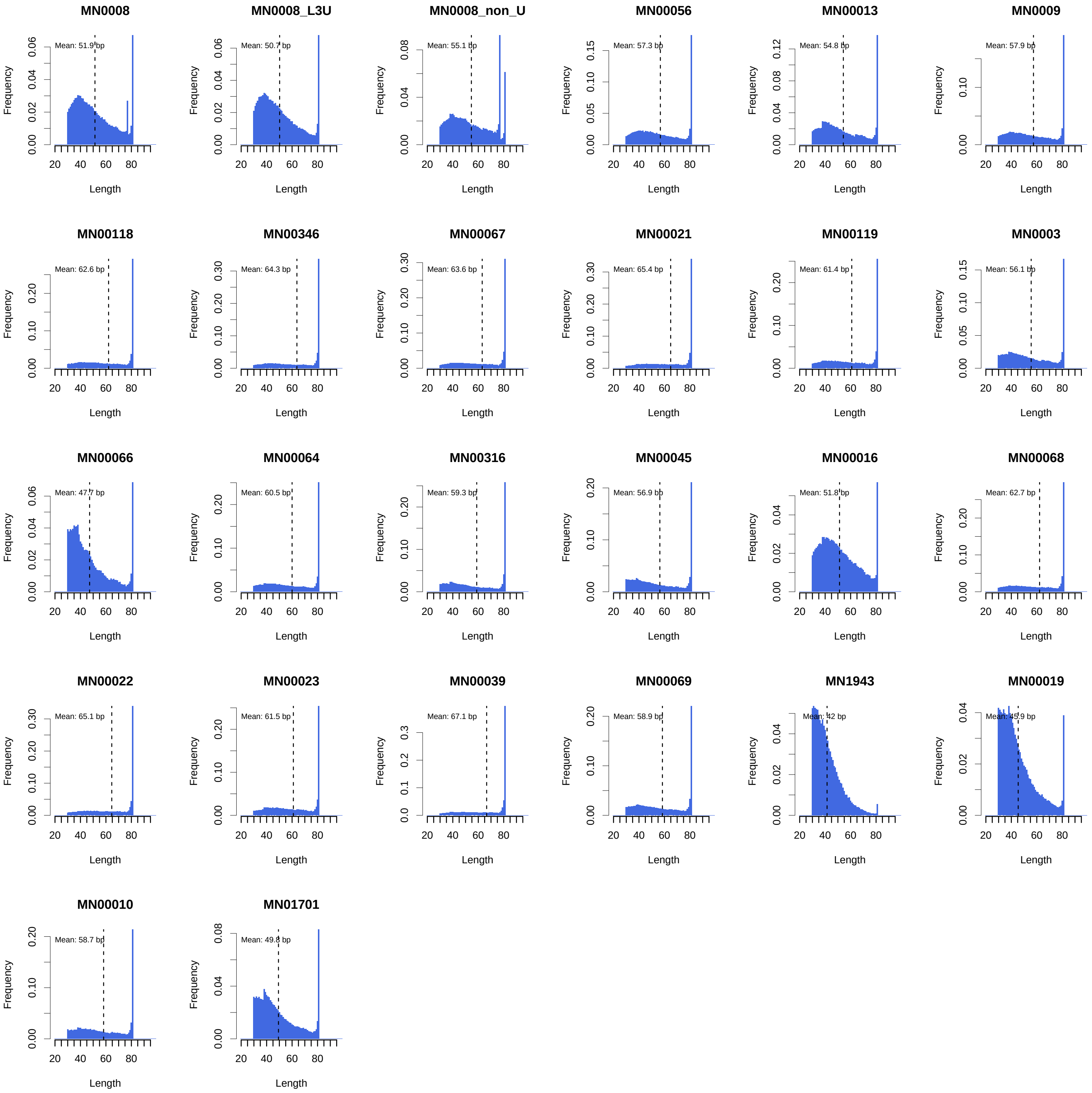


**Figure S2. Read length distribution in base pairs (bp) for mapped reads.** For the individual sample MN0008, read lengths are shown for the data of all the libraries (MN0008), as well as the USER-treated only (MN0008_L3U) and the standard non-USER treated libraries (MN0008_non_U). See Dataset S4 for the read length distribution plots corresponding to the substitution rates for each of the libraries sequenced per individual.

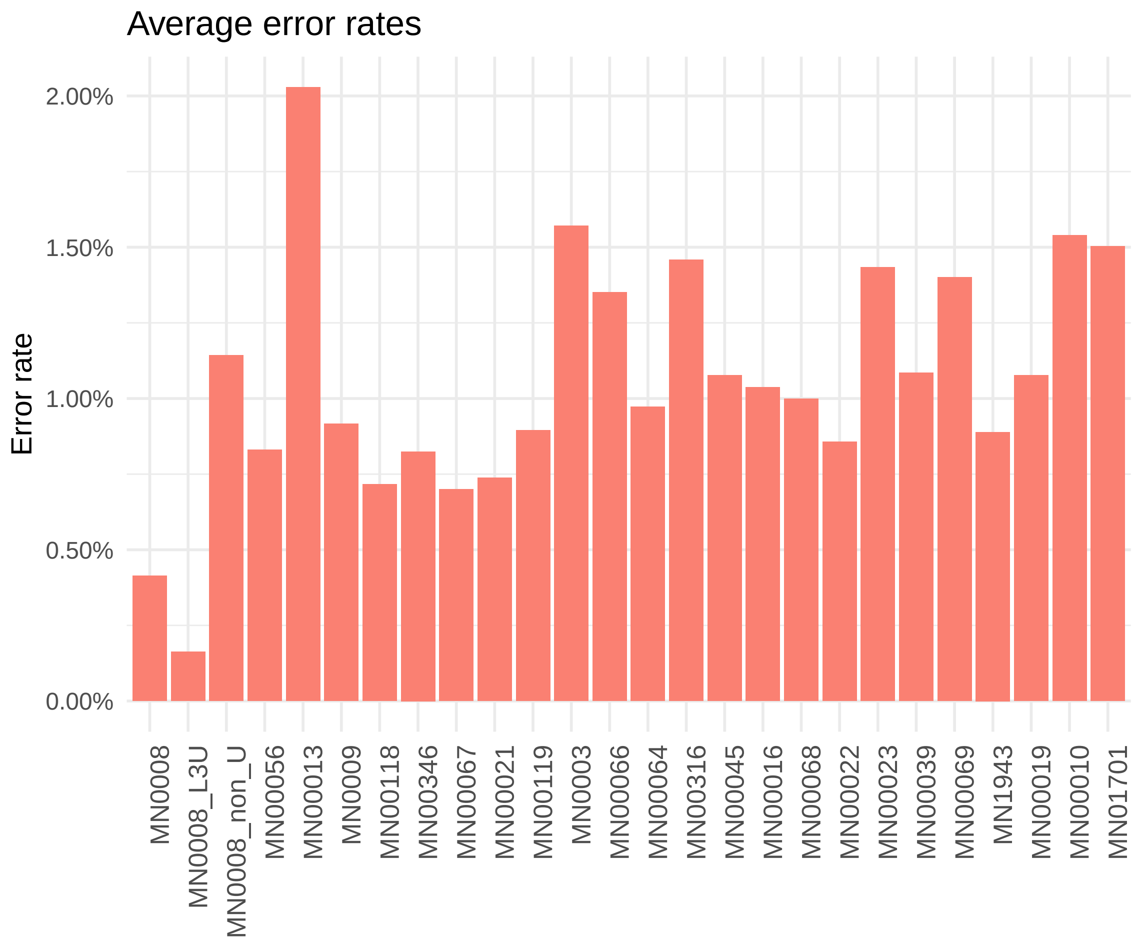


**Figure S3. Average error rates estimated with ANGSD.** For the sample MN0008, error rates are shown for the data of all the libraries (MN0008), as well as the USER-treated only (MN0008_L3U) and the standard non-USER treated libraries (MN0008_non_U). See “Error rate estimation” in Materials and Methods for a description of the estimation.

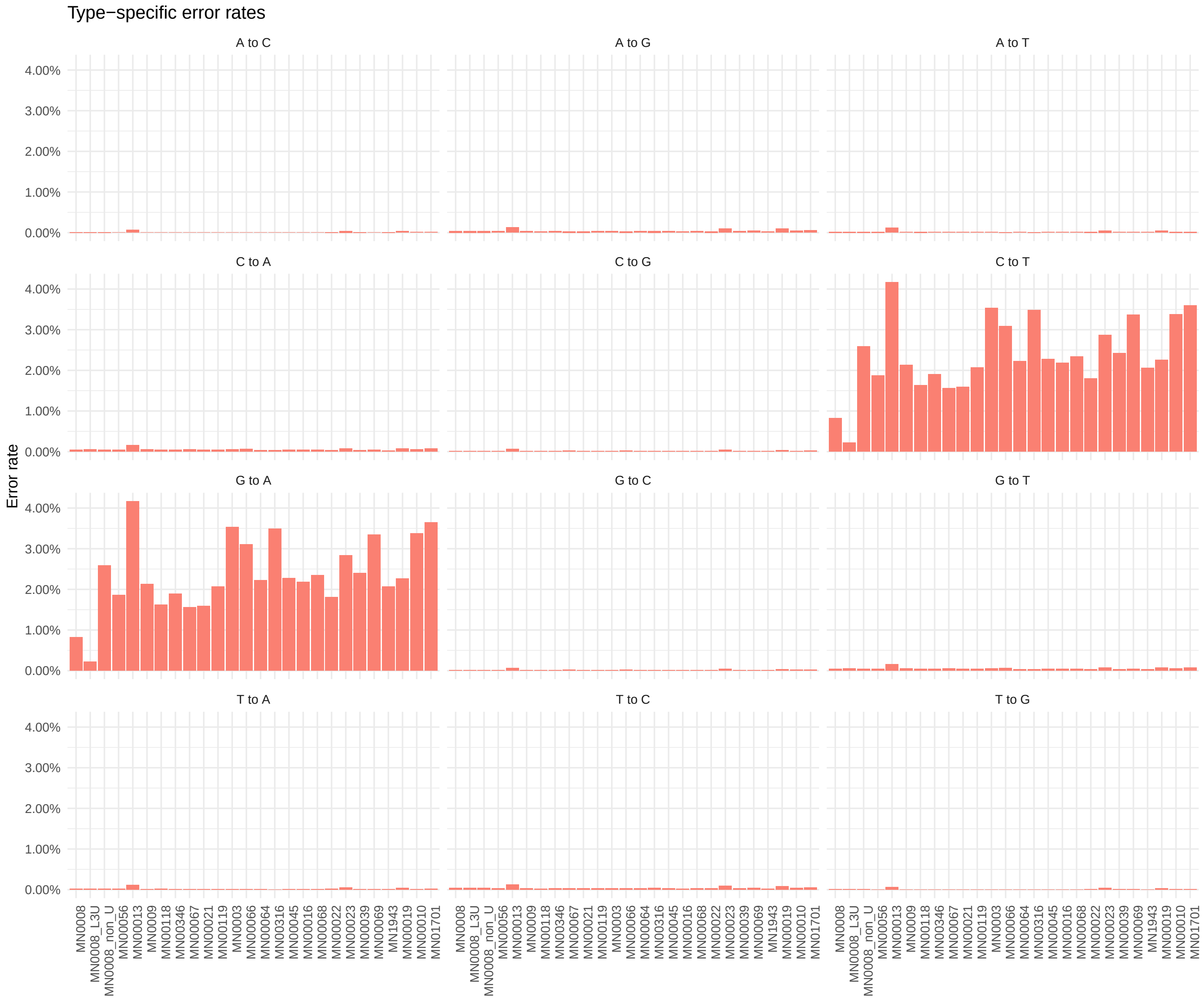


**Figure S4. Average error rates by substitution type.** The title of each panel indicates the substitution type for which error rates were estimated with ANGSD. See "Error rate estimation" in Materials and Methods for a description of the estimation.

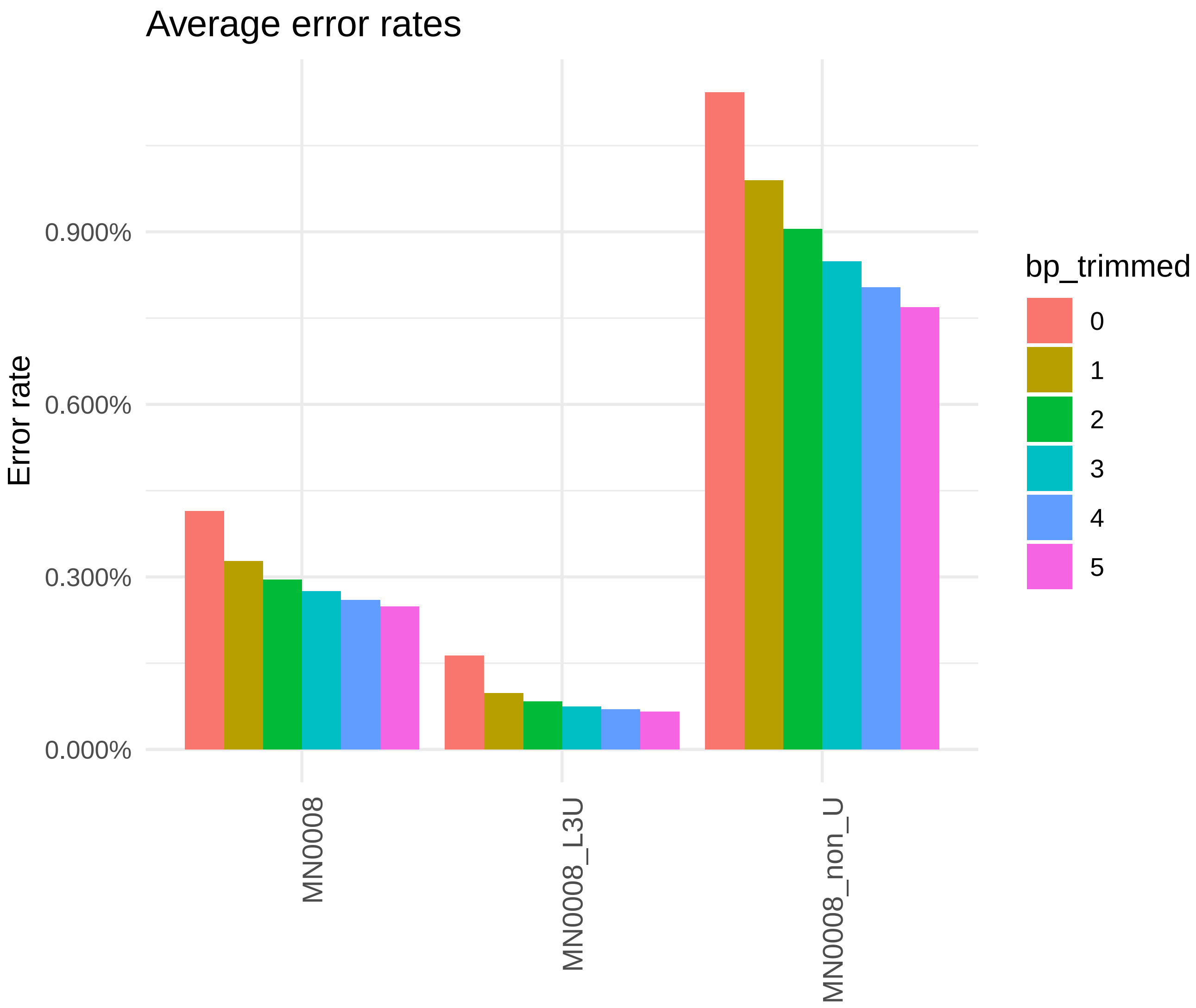


**Figure S5. Average error rates for MN0008.** Error rates estimated with ANGSD for the individual MN0008 for the data of all the libraries (MN0008), as well as the USER-treated only (MN0008_L3U) and the standard non-USER treated libraries (MN0008_non_U). The estimation was done using the complete reads (i.e., 0 bp trimmed) or trimming between 1 and 5 bp on each end of the mapped reads. See “Error rate estimation” in Materials and Methods for a full description of the estimation.

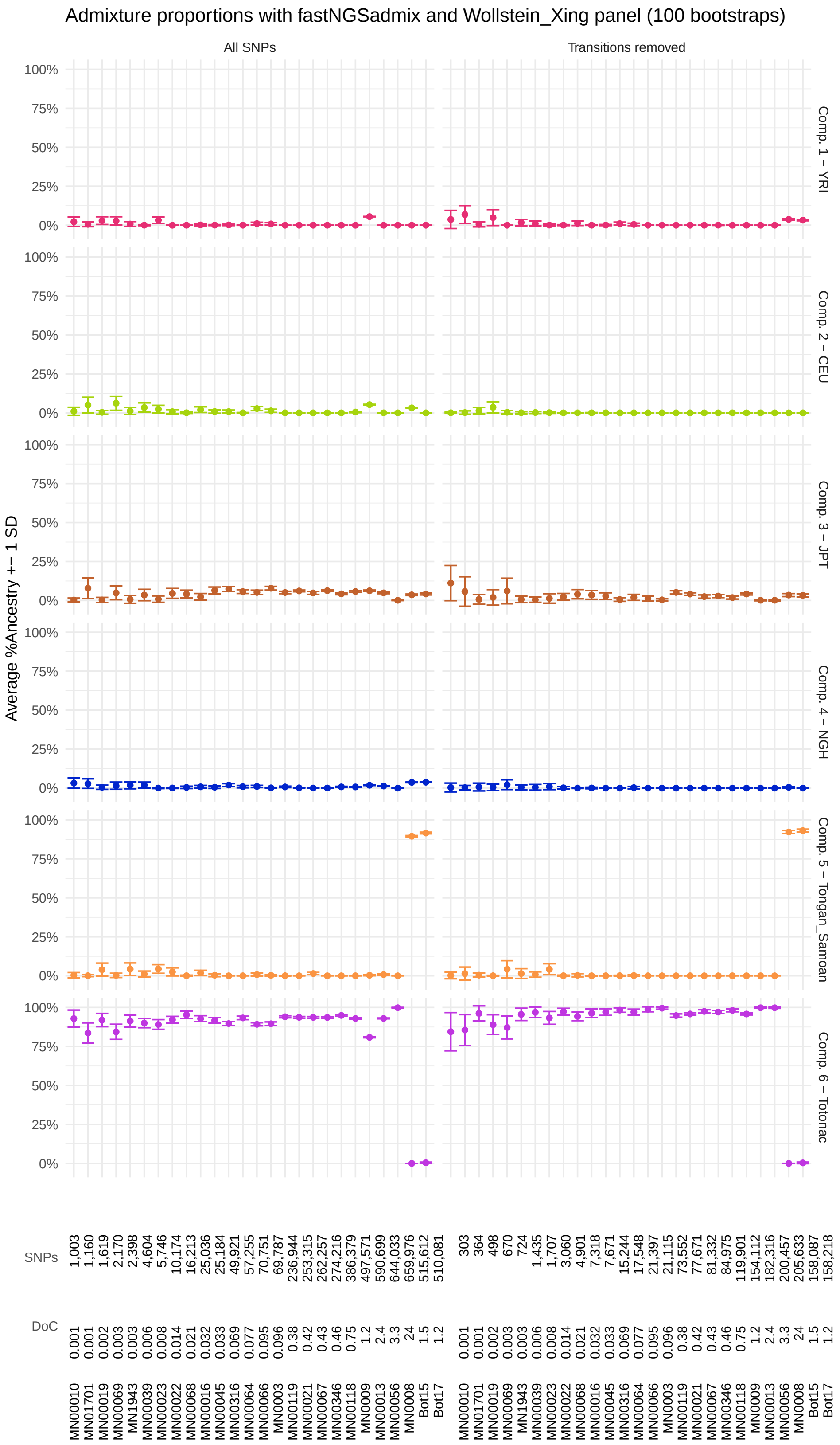


**Figure S6. Average ancestry proportions estimated with fastNGSadmix and the panel NGSadmix_fastNGSadmix_Wollstein_Xing, assuming K = 6 ancestral components.** Ancestry proportions were estimated using all available SNPs in an individual (left column) or only transversion SNPs (right column). The depth of coverage and number of SNPs analyzed are indicated above the ID of the individual. Error bars indicate one standard deviation for the estimation with 100 bootstraps. See "Clustering analyses – Ancestry estimation" and "Reference panels and dataset merging" in Materials and Methods for details on the fastNGSadmix analyses run and the reference panel used (Dataset S19), respectively.

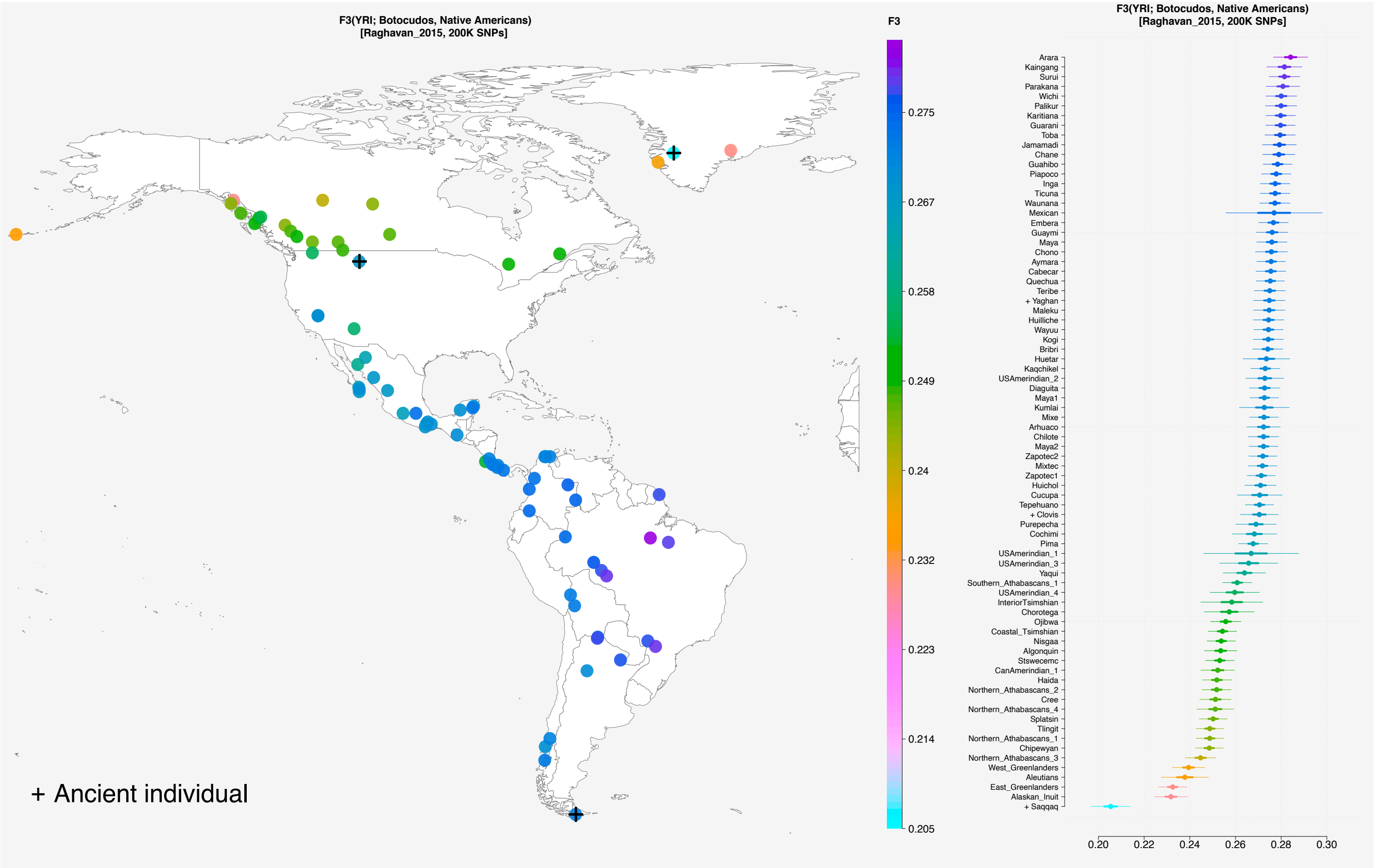


**Figure S7. Scores for outgroup-F3 tests of the form F3(YRI; "Botocudos", Native American) with F3_NGSadmix_Raghavan panel (199,285 SNPs).** The location of the populations is shown in the map with a dot colored according to their score, as indicated in the color bar. Populations with ancient individuals are labeled with a plus mark ("+"). The populations are listed on the right panel, with the highest scoring populations on the top, and their score plotted on the x-axis. Thick and thin bars represent one and three standard errors, respectively. Scores are shown for tests performed on at least 5,000 SNPs. The data of the 22 "Botocudos" sequenced in this study were included here. See "Reference panels and dataset merging" in Materials and Methods and Dataset S19 for a description of the panel used in this analysis.

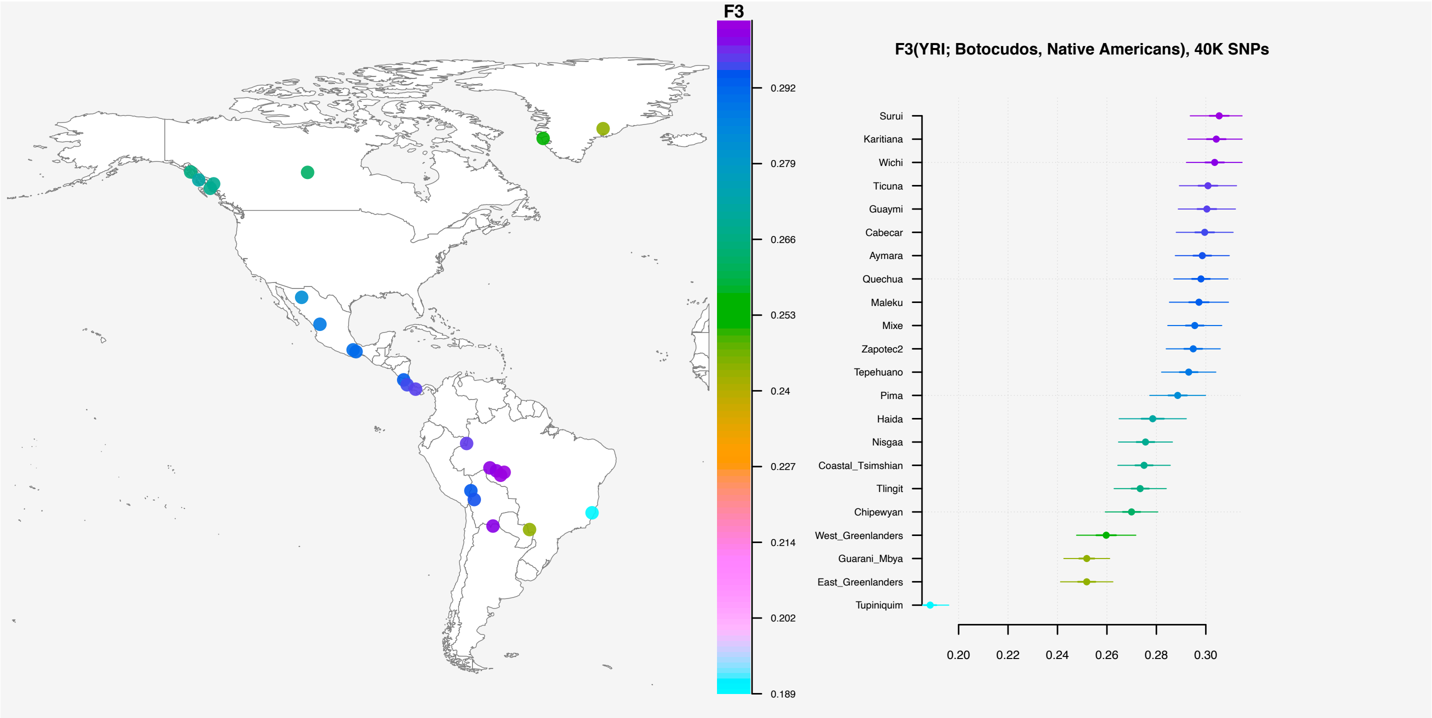


**Figure S8. Scores for outgroup-F3 tests of the form F3(YRI; "Botocudos", Native American) with F3_NGSadmix_Raghavan_CastroTupGuaUnrel panel (40,146 SNPs).** The location of the populations is shown in the map with a dot colored according to their score, as indicated in the color bar. Populations with ancient individuals are labeled with a plus mark ("+"). The populations are listed on the right panel, with the highest scoring populations on the top, and their score plotted on the x-axis. Thick and thin bars represent one and three standard errors, respectively. Scores are shown for tests performed on at least 5,000 SNPs. Individuals from the Guarani Mbyá and Tupiniquim populations have recent non-Native American admixture (not masked in this dataset). See Dataset S11 for NGSadmix runs of this panel. The data of the 22 "Botocudos" sequenced in this study were included here. See "Reference panels and dataset merging" in Materials and Methods and Dataset S19 for a description of the panel used in this analysis.

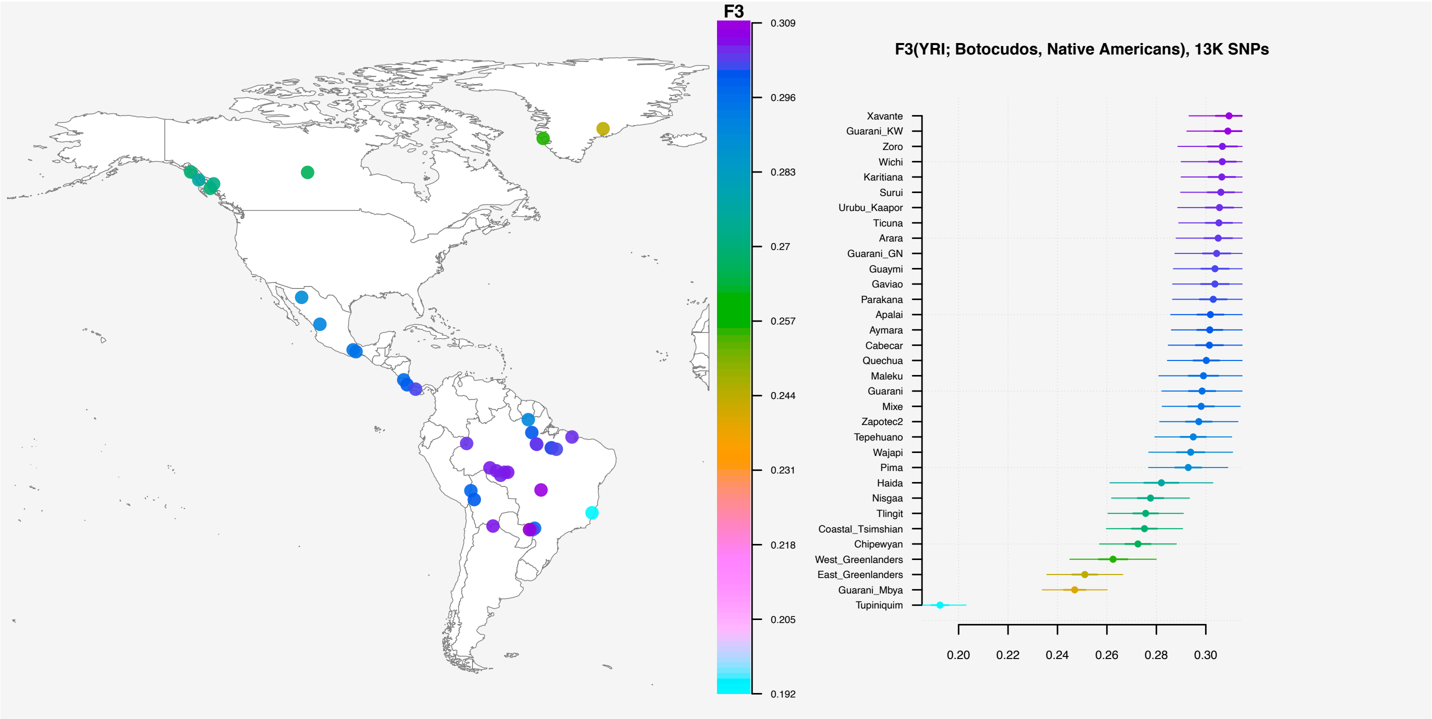


**Figure S9. Scores for outgroup-F3 tests of the form F3(YRI; "Botocudos", Native American) with F3_NGSadmix_Raghavan_CastroTupGuaUnrel_Castro12NatAm_Skoglund panel (13,490 SNPs).** The location of the populations is shown in the map with a dot colored according to their score, as indicated in the color bar. Populations with ancient individuals are labeled with a plus mark ("+"). The populations are listed on the right panel, with the highest scoring populations on the top, and their score plotted on the x-axis. Thick and thin bars represent one and three standard errors, respectively. Scores are shown for tests performed on at least 5,000 SNPs. Individuals from the Guarani Mbyá and Tupiniquim populations have recent non-Native American admixture (not masked in this dataset. See Dataset S12 for NGSadmix runs of this panel. The data of the 22 "Botocudos" sequenced in this study were included here. See "Reference panels and dataset merging" in Materials and Methods and Dataset S19 for a description of the panel used in this analysis.

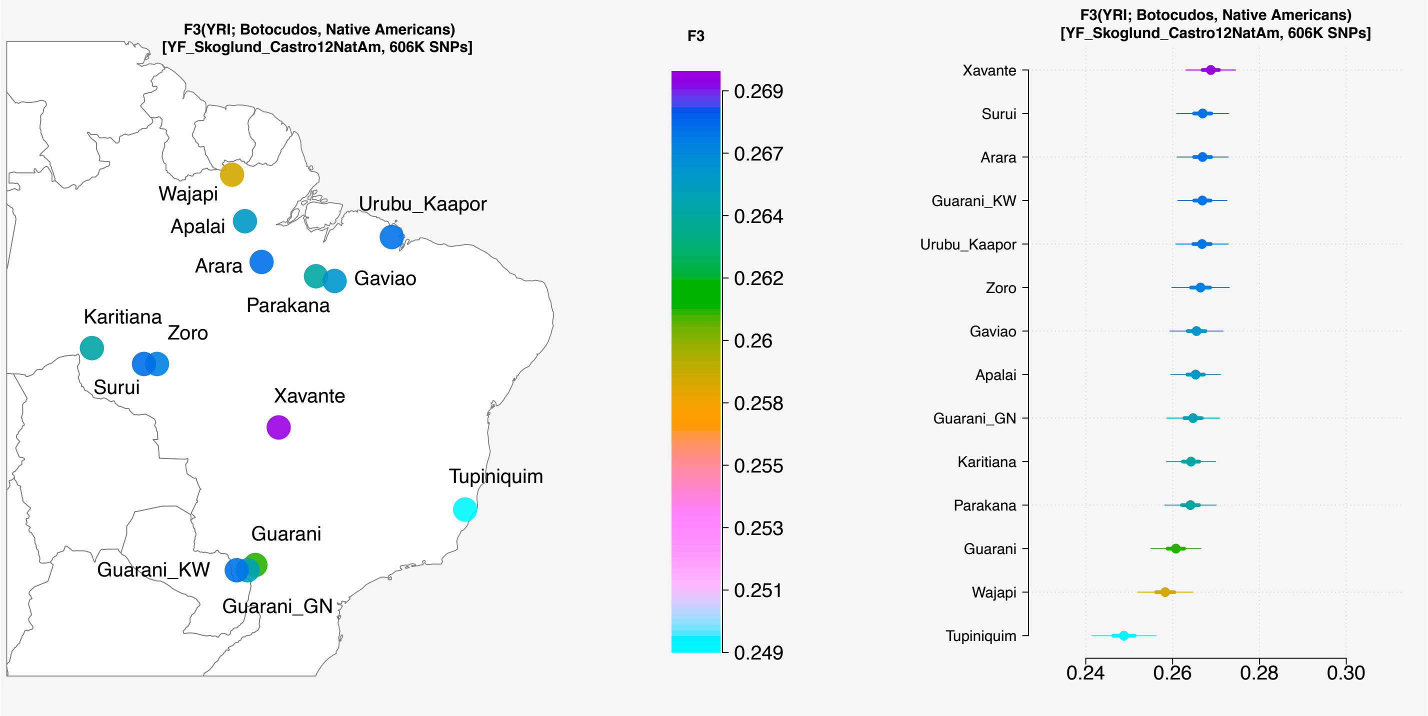


**Figure S10. Scores for outgroup-F3 tests of the form F3(YRI; "Botocudos", Native American) with F3_NGSadmix_Skoglund_Castro12NatAm panel (606,888 SNPs).** The location of the populations is shown in the map with a dot colored according to their score, as indicated in the color bar. Populations with ancient individuals are labeled with a plus mark ("+"). The populations are listed on the right panel, with the highest scoring populations on the top, and their score plotted on the x-axis. Thick and thin bars represent one and three standard errors, respectively. Scores are shown for tests performed on at least 5,000 SNPs. Individuals from the Tupiniquim population have recent non-Native American admixture (not masked in this dataset). See Dataset S13 for NGSadmix runs of this panel.

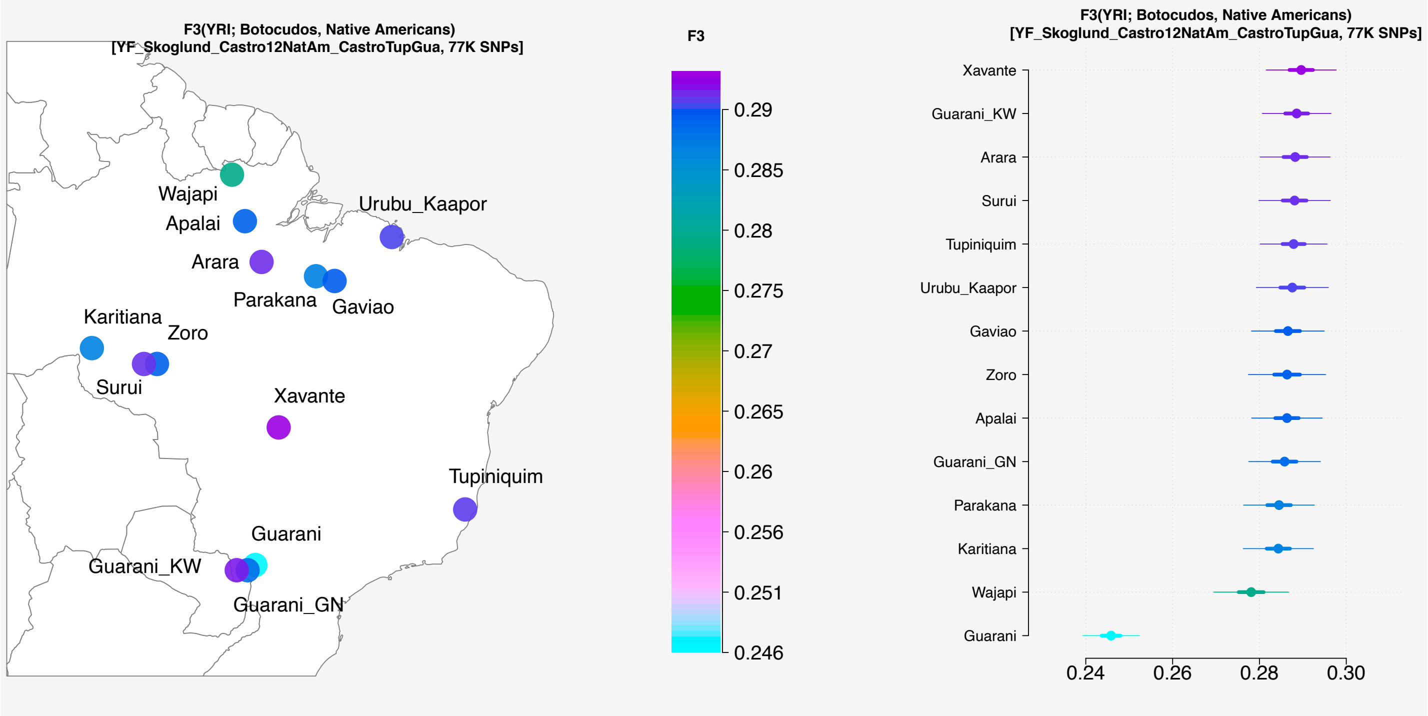


**Figure S11. Scores for outgroup-F3 tests of the form F3(YRI; "Botocudos", Native American) with F3_NGSadmix_Skoglund_CastroTupGuaUnrel_Castro12NatAm panel (77,766 SNPs).** The location of the populations is shown in the map with a dot colored according to their score, as indicated in the color bar. Populations with ancient individuals are labeled with a plus mark ("+"). The populations are listed on the right panel, with the highest scoring populations on the top, and their score plotted on the x-axis. Thick and thin bars represent one and three standard errors, respectively. Scores are shown for tests performed on at least 5,000 SNPs. Individuals from the Guarani population have recent non-Native American admixture (not masked in this dataset). See Dataset S14 for NGSadmix runs of this panel.

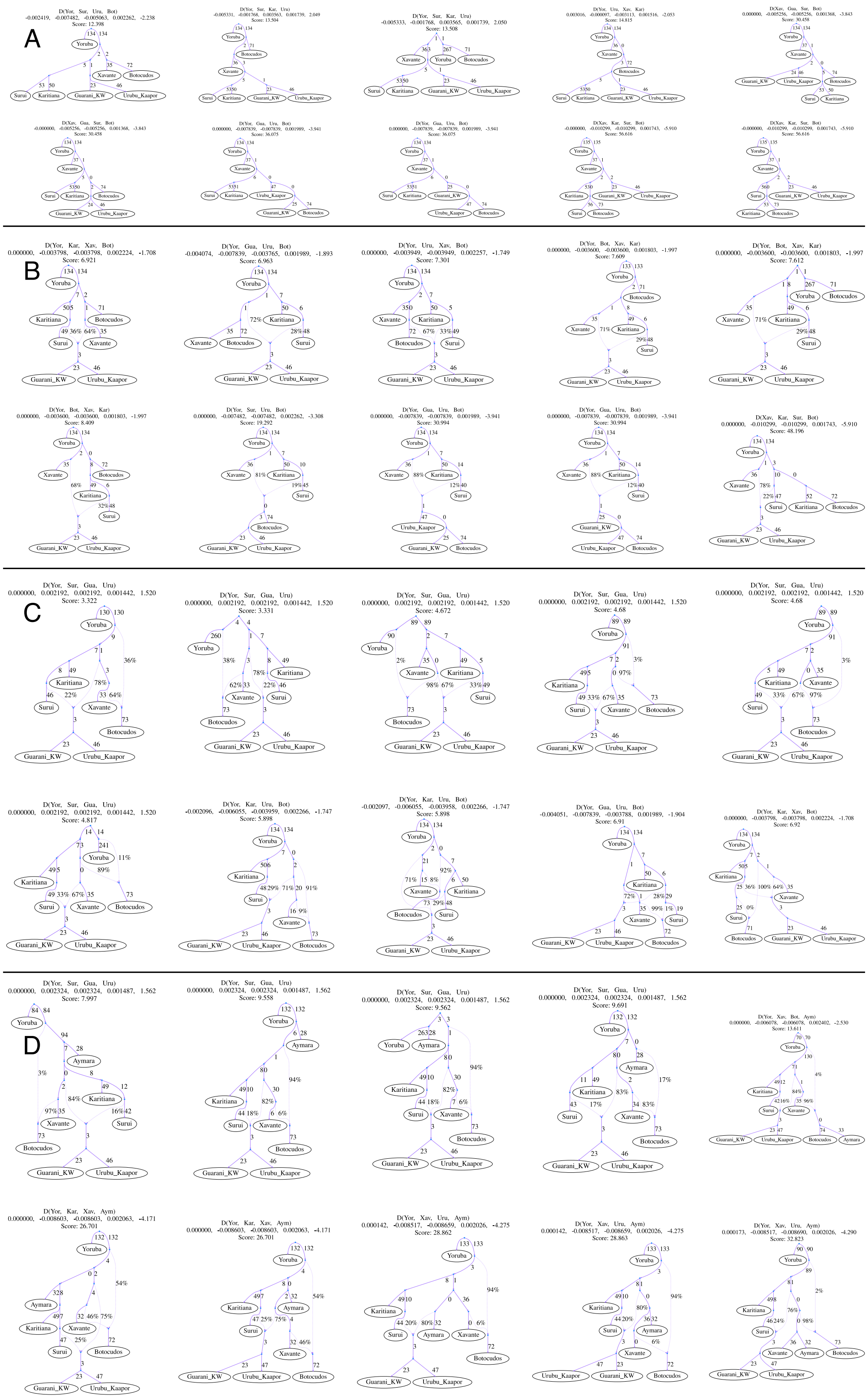


**Figure S12. Admixture graphs modelling the ancestry of the ancient "Botocudos", Jê- (Xavante), Tupi-speaking groups (Karitiana, Surui, Urubu Kaápor, Guarani Kaiowá).** For each of the models in Figure 5, we show the ten best-fitting graphs. Additionally, in D., we show the ten best fitting graphs after adding the Aymara as a leaf to all possible branches of the graph in Figure 5C. For each graph, we show the four populations producing the worst residual between the observed and the expected f-statistics; the observed value for such statistic, the expected value under the model, the residual, the standard error of the residual and a Z-score for such residual; and the model fit score. See "Reference panels and dataset merging" and Dataset S19 for a description of the panel qpGraph_Skoglund_Castro used in this analysis.

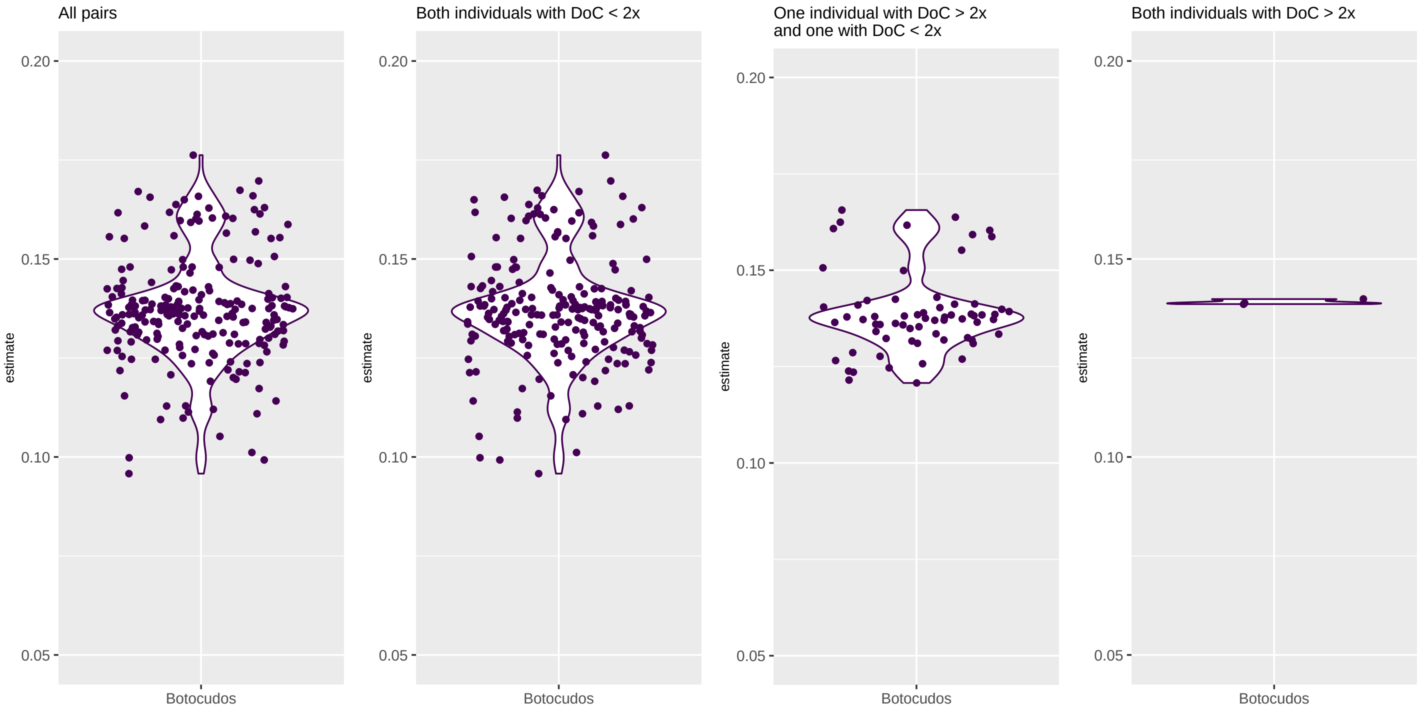


**Figure S13. Conditional heterozygosity for all pairs of "Botocudo" individuals.** Each dot corresponds to the point estimate for a single test involving, from left to right: any two "Botocudos"; pairs with a DoC below 2x each; one with DoC below and one with DoC above 2x; and both with DoC above 2x. The violin plots represent the smoothed density of the data points (heterozygosity estimates), with wider intervals corresponding to higher densities. See "Conditional heterozygosity estimation" in Materials and Methods for a description of the heterozygosity analysis.

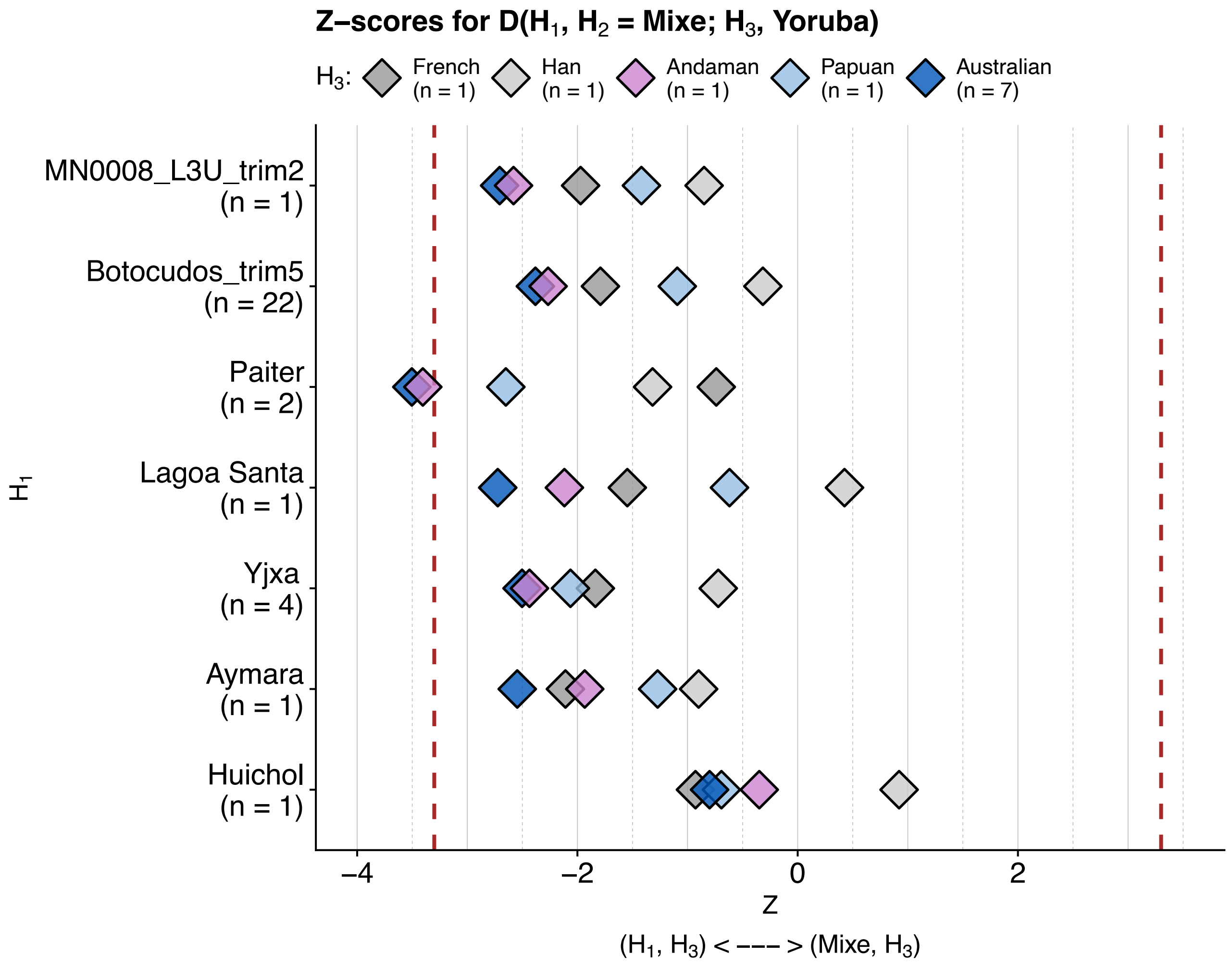


**Figure S14. Error-corrected D-statistics.** Z-scores for error-corrected D-statistics D(H1, H2 = Mixe; H3, Yoruba). Y-axis: population used in the test as H1. Color code: population used in the test as H3. When H3 is closer to H1 than to Mixe, the Z-scores become negative (left-hand side of the plot). In contrast, positive Z-scores (right-hand side of the plot) indicate a stronger proximity between Mixe and H3 than between H1 and H3. The number of genomes per population is indicated between parentheses. D-statistics were corrected according to the error rates estimated per genome following (Soraggi, Wiuf, & Albrechtsen, 2018). Z-scores were obtained after a jackknife resampling approach of D-statistic for 5 Mb-blocks. Vertical dotted lines are placed at |3.3| values and correspond to a p-value of approximately 0.005 (Green et al., 2010). For the "Botocudo" individual MN0008, only reads from the USER-treated library were analyzed. Reads from ancient genomes were trimmed either 2 bp (MN0008) or 5 bp (LagoaSanta, Andaman, 22 "Botocudos") on each end. See "Error-corrected D-statistics" and "Reference panels and dataset merging" in Materials and methods for a description of the analysis run and the panel ErrorCorr_Dstat (Dataset S19) used here.

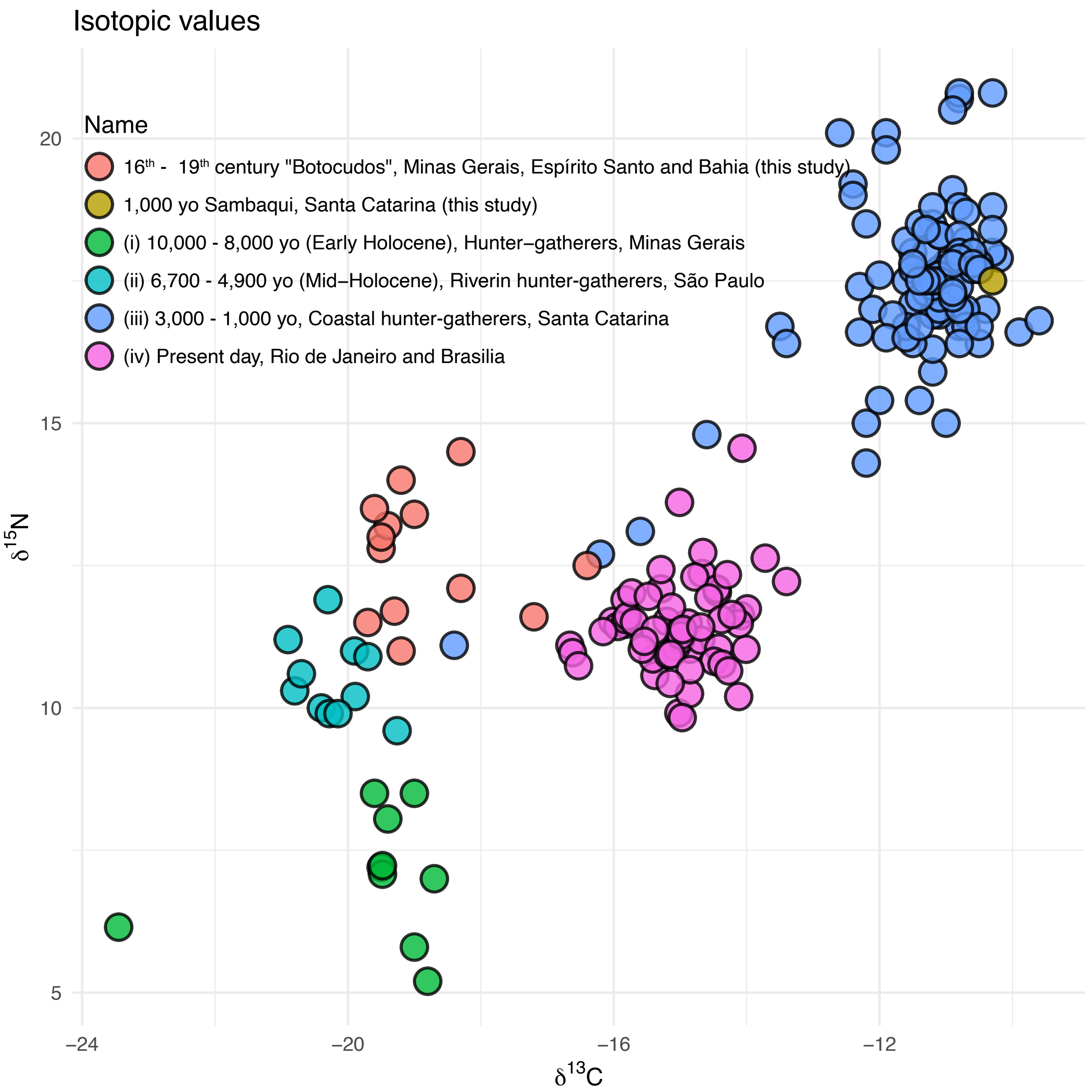


**Figure S15. Isotopic values measured for the individuals in this study and external datasets.** Each dot corresponds to d δ13C and δ15N values measured per individual. Only bones and permanent second and third molars are included in this plot. See “Diet characterization” in Materials and Methods and Dataset S18 for a description of the values plotted in this figure.

#### *Datasets (S1 to S19)*

**Dataset S1 (DS1_Ethics.pdf).** Permit granted by the IPHAN (Instituto do Patrimônio Histórico e Artístico Nacional) to study the skeletal remains of 24 individuals from the Anthropological collection of the Museu Nacional, and registration of microbial species at the Sistema Nacional De Gestão Do Patrimônio Genético E Do Conhecimento Tradicional Associado.

**Dataset S2 (DS2_Sequencing_stats.xlsx).** Excel workbook containing sample information and sequencing statistics per individual (sheet 1) and per library (sheet 2), as well as contamination estimates per individual (sheet 3).

**Dataset S3 (DS3_Damage_lib.pdf).** Plots with substitution rates on mapped reads. Each row of plots corresponds to the data of a single individual, and each panel shows the substitution frequencies per library. Libraries are labeled as mtCapture, L1, L2 and L3U, the latter being USER-treated (individual MN0008 only).

**Dataset S4 (DS4_Length_dist_libs.pdf).** Length distributions of mapped reads per individual and library. Each row of plots corresponds to the data of a single individual, and each panel shows the read length per library. Libraries are labeled as mtCapture, L1, L2 and L3U, the latter being USER-treated (individual MN0008 only).

**Dataset S5 (DS5_MDS_Wollstein_AllSamples.pdf).** First two dimension of multidimensional scaling analyses for each of the individual sequenced in this study, with all SNPs overlapping the MDS_NGSadmix_Wollstein_Xing_Malaspinas panel described in Materials and Methods. The title of each plot indicates the individual that was merged to the panel, represented with a black dot on the plot. Each plot is labeled with the number of SNPs overlapping with the panel in the individual that was merged, the fraction of the panel covered and the depth of coverage of the individual.

**Dataset S6 (DS6_MDS_Wollstein_AllSamples_rmTrans.pdf).** First two dimension of multidimensional scaling analyses for each of the individual sequenced in this study, with transversion SNPs overlapping the MDS_NGSadmix_Wollstein_Xing_Malaspinas panel described in Materials and Methods. The title of each plot indicates the individual that was merged to the panel, represented with a black dot on the plot. Each plot is labeled with the number of SNPs overlapping with the panel in the individual that was merged, the fraction of the panel covered and the depth of coverage of the individual.

**Dataset S7 (DS7_Ancestry_fastNGSadmix_Wollstein.xlsx).** Average ancestry proportions estimated with fastNGSadmix and assuming K = 6 ancestral components. Ancestry proportions were estimated using all available SNPs in an individual (rmTrans = FALSE) or only transversion SNPs (rmTrans = TRUE). The depth of coverage and number of SNPs analyzed are indicated next to the ID of the individual. Ancestry proportions are reported as average +- 1 standard deviation (100 bootstraps). These values are shown in Fig. S6.

**Dataset S8 (DS8_NGSadmix_Wollstein_k6_24samples.pdf).** Ancestry proportions estimated with NGSadmix assuming K = 6 ancestral components. Each of the 24 individuals sequenced in this study was merged to the panel MDS_NGSadmix_Wollstein_Xing_Malaspinas, and all the SNPs with data on the target individual were selected for each of the NGSadmix runs. Rows correspond to one run on the panel plus the target individual. The target individual, number of SNPs analyzed and depth of coverage of the target individual is indicated in the title of each plot.

**Dataset S9 (DS9_NGSadmix_Wollstein_rmTrans_k6_24samples.pdf).** Ancestry proportions estimated with NGSadmix assuming K = 6 ancestral components. Each of the 24 individuals sequenced in this study was merged to the panel MDS_NGSadmix_Wollstein_Xing_Malaspinas, and all the transversion SNPs with data on the target individual were selected for each of the NGSadmix runs. Rows correspond to one run on the panel plus the target individual. The target individual, number of SNPs analyzed and depth of coverage of the target individual is indicated in the title of each plot.

**Dataset S10 (DS10_NGSadmix_Raghavan_Skoglund_Castro_k2-16.pdf).** Ancestry proportions estimated with NGSadmix for the 24 individuals sequenced in this study and the panel NGSadmix_Raghavan_Skoglund_Castro (56,646 SNPs). The reads of the 24 individuals were trimmed 5 bp on each end prior to genotype likelihoods calculation. Each page shows the best run among 10 replicates assuming between K = 2 and K = 16 ancestral components. Fig. 4 shows an NGSadmix run with this panel and only 9 "Botocudos" and assuming K = 13 components.

**Dataset S11 (DS11_NGSadmix_Raghavan_CastroTupGuaUnrel_k2-17.pdf).** Ancestry proportions estimated with NGSadmix for the 24 individuals sequenced in this study and the panel F3_NGSadmix_Raghavan_CastroTupGuaUnrel (40,146 SNPs). The reads of the 24 individuals were trimmed 5 bp on each end prior to genotype likelihoods calculation. Each page shows the best run among 10 replicates assuming between K = 2 and K = 17 ancestral components. See Fig. S8 for outgroup-F3 analysis in this panel.

**Dataset S12 (DS12_NGSadmix_Raghavan_CastroTupGuaUnrel_Castro12NatAm_Skoglund_k2-17.pdf).** Ancestry proportions estimated with NGSadmix for the 24 individuals sequenced in this study and the panel F3_NGSadmix_Raghavan_CastroTupGuaUnrel_Castro12NatAm_Skoglund (13,490 SNPs). The reads of the 24 individuals were trimmed 5 bp on each end prior to genotype likelihoods calculation. Each page shows the best run among 10 replicates assuming between K = 2 and K = 17 ancestral components. See Fig. S9 for outgroup-F3 analysis in this panel.

**Dataset S13 (DS13_NGSadmix_Skoglund_Castro12NatAm_k2-6.pdf).** Ancestry proportions estimated with NGSadmix for the 24 individuals sequenced in this study and the panel F3_NGSadmix_Skoglund_Castro12NatAm (606,888 SNPs). The reads of the 24 individuals were trimmed 5 bp on each end prior to genotype likelihoods calculation. Each page shows the best run among 10 replicates assuming between K = 2 and K = 17 ancestral components. See Fig. S10 for outgroup-F3 analysis in this panel.

**Dataset S14 (DS14_NGSadmix_Skoglund_CastroTupGuaUnrelated_Castro12NatAm_k2-6.pdf).** Ancestry proportions estimated with NGSadmix for the 24 individuals sequenced in this study and the panel F3_NGSadmix_Skoglund_CastroTupGuaUnrel_Castro12NatAm (77,766 SNPs). The reads of the 24 individuals were trimmed 5 bp on each end prior to genotype likelihoods calculation. Each page shows the best run among 10 replicates assuming between K = 2 and K = 17 ancestral components. See Fig. S11 for outgroup-$f_3$ analysis in this panel.

**Dataset S15 (DS15_Metagenomics.xlsx).** Taxonomic classifications by Kraken2 for the 24 individuals sequenced in this study (sheet 1), most abudant bacteria identified across individuals (sheet 2), mapping statistics for eight oral microbes for all individuals (sheet 3) and mapping statistics for eight oral microbes with a depth of coverage above 1× (sheet 4).

**Dataset S16 (DS16_DIAMOND_hits_by_virus.xlsx).** Number of reads classified by DIAMOND for each virus per individual, after selecting only viruses that can be hosted by humans.

**Dataset S17 (DS17_BWA_candidate_virus_q0.xlsx).** Statistics corresponding to the mappings with BWA of the non-human reads for each of the 24 individuals to the reference genomes of the 50 viruses listed in Dataset S16.

**Dataset S18 (DS18_isotopic_values.xlsx).** Isotopic values measured for 22 individuals in this study and four previously published datasets. The values are displayed on Fig. S15. Dataset described in "Diet reconstruction" in Materials and Methods.

**Dataset S19 (DS19_Datasets_methods.xlsx).** Description of genomic datasets described in Materials and Methods. Sheets 1 to 3 list datasets used to produce the main and supplementary figures, and the datasets in Sheet 4 were used only for supplementary figures. See "Reference panels and datasets merging" for a description of the panels included in this workbook.
